# Calcitriol, the active form of vitamin D, is a promising candidate for COVID-19 prophylaxis

**DOI:** 10.1101/2020.06.21.162396

**Authors:** Chee Keng Mok, Yan Ling Ng, Bintou Ahmadou Ahidjo, Regina Ching Hua Lee, Marcus Wing Choy Loe, Jing Liu, Kai Sen Tan, Parveen Kaur, Wee Joo Chng, John Eu-Li Wong, De Yun Wang, Erwei Hao, Xiaotao Hou, Yong Wah Tan, Tze Minn Mak, Cui Lin, Raymond Lin, Paul Tambyah, JiaGang Deng, Justin Jang Hann Chu

## Abstract

COVID-19, the disease caused by SARS-CoV-2 (1), was declared a pandemic by the World Health Organization (WHO) in March 2020 (2). While awaiting a vaccine, several antivirals are being used to manage the disease with limited success (3, 4). To expand this arsenal, we screened 4 compound libraries: a United States Food and Drug Administration (FDA) approved drug library, an angiotensin converting enzyme-2 (ACE2) targeted compound library, a flavonoid compound library as well as a natural product library. Of the 121 compounds identified with activity against SARS-CoV-2, 7 were shortlisted for validation. We show for the first time that the active form of Vitamin D, calcitriol, exhibits significant potent activity against SARS-CoV-2. This finding paves the way for consideration of host-directed therapies for ring prophylaxis of contacts of SARS-CoV-2 patients.

---

Despite implementation of physical distancing, mask wearing, quarantine and the tireless efforts expended for contact tracing, the rapid transmissibility of SARS-CoV-2 even during the asymptomatic phase has made containment of this virus extremely difficult. The main proposed strategy to curb this pandemic is the implementation of mass vaccination programs. Once a suitable vaccine is discovered, the significant challenges associated with vaccination programs e.g. limitations in manufacturing capabilities and associated costs, are anticipated to significantly affect uptake of vaccinations globally. We therefore propose that ring prophylaxis, which had been previously proposed for influenza pandemics and involves treating close contacts of a confirmed case with an antiviral prophylaxis to further curb community spread(5), be considered as a viable strategy to reduce transmission of SARS-CoV-2.

In an effort to identify potential candidates for SARS-CoV-2 chemoprophylaxis, we performed a virus-induced cytopathic effect (CPE) based screen of several small molecule libraries in SARS-CoV-2-infected Vero E6 cells (Fig. 1a). The African green monkey kidney epithelial Vero E6 cells were used for the screen as these cells are highly susceptible to coronaviruses and exhibit obvious CPE upon infection. A 57-compound natural product library and a library of 462 ACE2 targeted inhibitors (the ACE2 receptor was identified to be necessary for SARS-CoV-2 infection (*6*)) were used in a pre-infection treatment screen to identify potential viral entry inhibitors, while a post-infection treatment screen was performed using both a 500 compound flavonoid library and a 1172 FDA-approved compound libraries in order to identify potential inhibitors targeting post-entry steps of the SARS-CoV-2 replication cycle. For the pre-infection treatment screen, Vero E6 cells were treated with compounds for two hours prior to infection with SARS-CoV-2. The post-infection treatment screen on the other hand was performed by adding compounds to the Vero E6 cells 1 hour post-infection with SARS-CoV-2. Compounds which showed less than 50% CPE compared to the 0.1%DMSO vehicle control with SARS-CoV-2 infection were identified as hits (Tables S1-S3). Using this method, we identified 31 compounds from the pre-infection treatment screen and 90 compounds from the post-infection treatment screen with activity against SARS-CoV-2 (Table S4). As expected, our hit list included the tyrosine kinase inhibitors masitinib and imatinib mesylate (*7*), the antiretroviral drug lopinavir(*8*) and the calpain inhibitor calpeptin (*9*) – all compounds reported to inhibit SARS-CoV, SARS-CoV-2 or MERS-CoV. This provides the robustness and confidence of our primary screen for potential antivirals against SARS-CoV-2.

**Figure 1:**
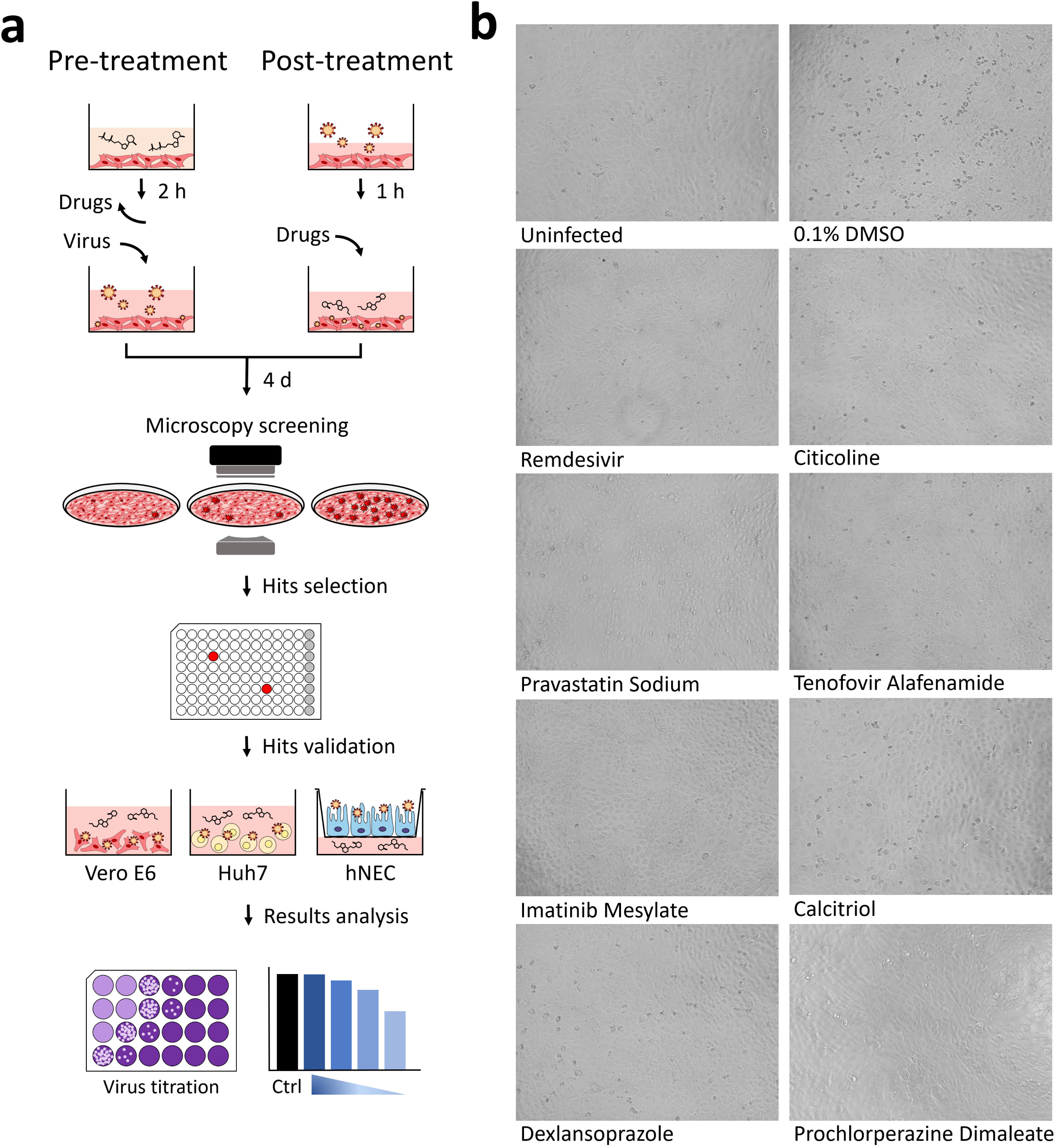
Schematic overview of the anti-SARS-CoV-2 drug screening procedure. (a) ACE2 targeted inhibitors and compounds from the natural product library were applied prior to infection with SARS-CoV-2 (pre-infection treatment) while compounds from the FDA-approved library and the flavonoid library were applied post infection (post-infection treatment). Out of 2191 compounds tested, 121 displayed viral CPE reductions and 7 were selected for further validations using Vero E6, HuH7, and hNEC cell lines. (b) Uninfected cells, infected cells treated with 0.1% DMSO and infected cells treated with remdesivir were included as controls.

Of these primary hits, 7 compounds were selected for downstream validation (Table 1). These included 3 compounds from the pre-infection treatment screen (citicoline, pravastatin sodium and tenofovir alafenamide) and four compounds from the post-infection treatment screen (imatinib mesylate, calcitriol, dexlansoprazole, and prochlorperazine dimaleate). These compounds were selected based on level of CPE inhibition in the primary screens (Fig. 1b), known mechanism of action and existing FDA approval or Generally Recognized as Safe (GRAS) status. FDA approval was considered an important factor as pre-existing data on safety and dosage would allow expedited decisions to be made regarding the potential use of these compounds in vulnerable populations to stymie the current pandemic. Validation assays to determine changes in infectious virus titres upon treatment was carried out by testing selected hit compounds in dose-dependent assays in Vero E6 to confirm the primary screen observation and also in the human hepatocarcinoma HuH7 cell line as the latter cell line expresses high levels of the ACE2 receptor (*10*) and supports replication of coronaviruses (*11*). Cell viability assays were also carried out to ensure that reduction of SARS-CoV-2 titres was not due to cytotoxic effects of the compounds on host cells. CC50, IC50 were obtained for each of the 7 compounds in Vero E6 cells (Table S5) and HuH7 cells (Table S6) and where possible selectivity index values were calculated. Pre-infection treatment of Vero E6 cells with citicoline and pravastatin sodium resulted in dose-dependent inhibition of SARS-CoV-2 (Fig. 2a & 2b), while a non-dose-dependent inhibition of SARS-CoV-2 at lower concentrations was observed with tenofovir alafenamide (Fig. 2c). These observations were however not recapitulated in HuH7 cell line (Fig. 2d, 2e & 2f). Post-infection treatment with ≥ 10 μM imatinib mesylate resulted in a reduction of SARS-CoV-2 titres to non-detectable levels in both Vero E6 and HuH7 cells (Fig. 2g & 2k), and a similar finding was observed with ≥ 10 μM prochlorperazine dimaleate in HuH7 cells (Fig. 2n). Significant reductions of at least 0.4 log_10_ of SARS-CoV-2 were also noted upon treatment with ≥10 μM dexlansoprazole (Fig. 2i, 2m) and post-infection treatment with 10 μM calcitriol resulted in a 1.3 log_10_ reduction of SARS-CoV-2 titre in Vero E6 cells (Fig. 2h). This finding could not be recapitulated in HuH7 cells (Fig. 2l) most likely because the CC50 value of calcitriol was 4.7 μM in HuH7 cells (Table S6). Given that the HuH7 cell line is a hepatocarcinoma cell line and therefore not the first point of entry for SARS-CoV-2 in humans, we decided to test the three most promising compounds (imatinib mesylate, citicoline and calcitriol) against SARS-CoV-2 in the primary human nasal epithelial cell line (hNEC) that is a known *in vivo* target of SARS-CoV-2 (*12*) (Fig. 1a). Despite its significant activity in the continuous cell lines (Vero E6 and HuH7), in hNECs, imatinib mesylate only displayed a 0.2 log_10_ reduction in viral titre (Fig. 3). Interestingly, out of the three compounds only calcitriol proved effective against SARS-CoV-2 with a reduction of 0.69 log_10_ in viral titre (Fig. 3). While recent data has shown that vitamin D levels are negatively associated with morbidity and mortality of COVID-19 cases (*13, 14*), this is the first report of a direct inhibitory effect of calcitriol on SARS-CoV-2.

**Table 1:**
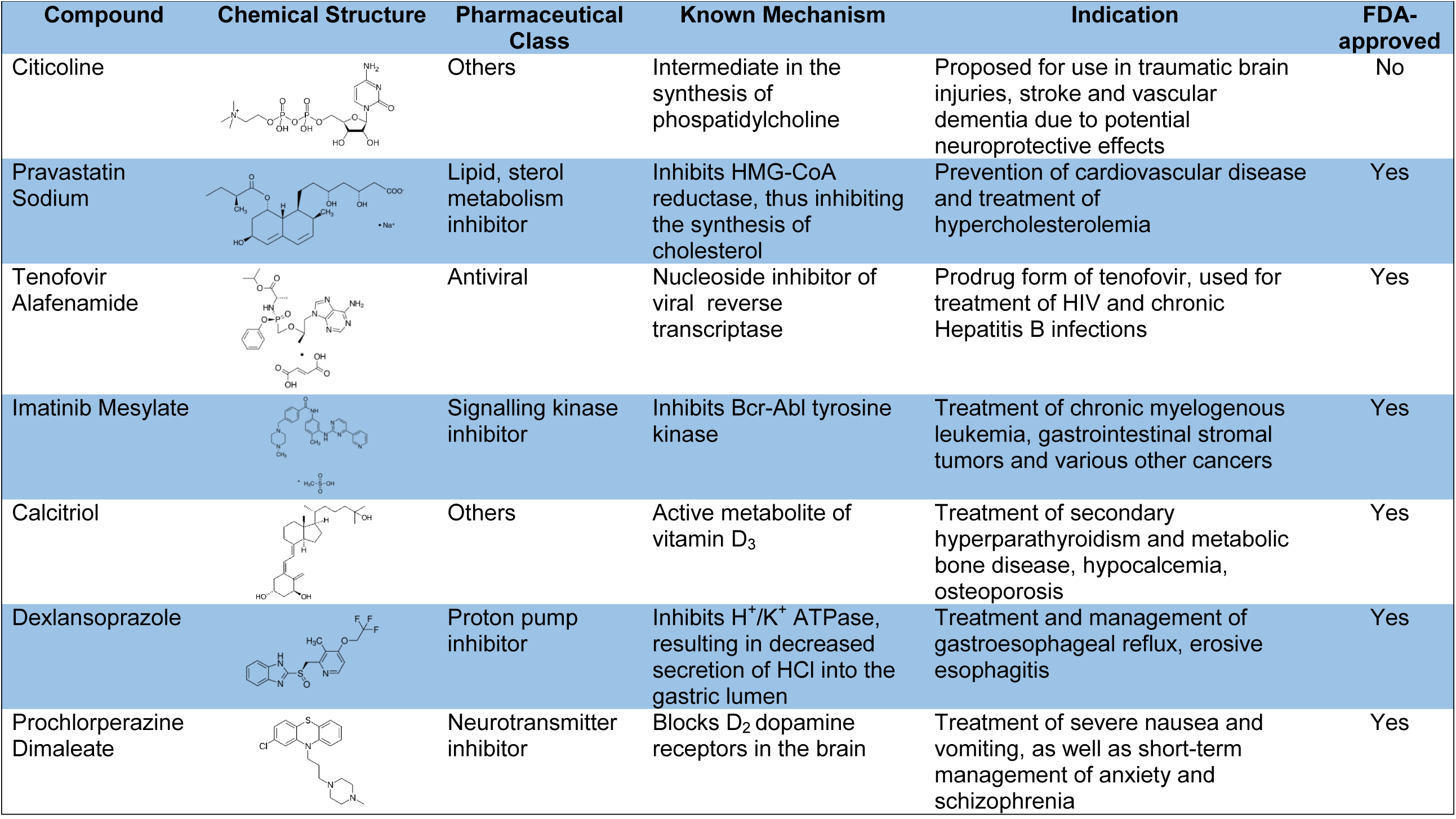
Summary of compounds identified from the primary screen that exhibit potential antiviral effects against SARS-CoV-2 infection.

| Compound | Chemical Structure | Pharmaceutical Class | Known Mechanism | Indication | FDA-approved |
| --- | --- | --- | --- | --- | --- |
| Citicoline |  | Others | Intermediate in the synthesis of phosphatidylcholine | Proposed for use in traumatic brain injuries, stroke and vascular dementia due to potential neuroprotective effects | No |
| Pravastatin Sodium |  | Lipid, sterol metabolism inhibitor | Inhibits HMG-CoA reductase, thus inhibiting the synthesis of cholesterol | Prevention of cardiovascular disease and treatment of hypercholesterolemia | Yes |
| Tenofovir Alafenamide |  | Antiviral | Nucleoside inhibitor of viral reverse transcriptase | Prodrug form of tenofovir, used for treatment of HIV and chronic Hepatitis B infections | Yes |
| Imatinib Mesylate |  | Signalling kinase inhibitor | Inhibits Bcr-Abl tyrosine kinase | Treatment of chronic myelogenous leukemia, gastrointestinal stromal tumors and various other cancers | Yes |
| Calcitriol |  | Others | Active metabolite of vitamin D<sub>3</sub> | Treatment of secondary hyperparathyroidism and metabolic bone disease, hypocalcemia, osteoporosis | Yes |
| Dexlansoprazole |  | Proton pump inhibitor | Inhibits H<sup>+</sup>/K<sup>+</sup> ATPase, resulting in decreased secretion of HCl into the gastric lumen | Treatment and management of gastroesophageal reflux, erosive esophagitis | Yes |
| Prochlorperazine Dimaleate |  | Neurotransmitter inhibitor | Blocks D<sub>2</sub> dopamine receptors in the brain | Treatment of severe nausea and vomiting, as well as short-term management of anxiety and schizophrenia | Yes |

**Figure 2:**
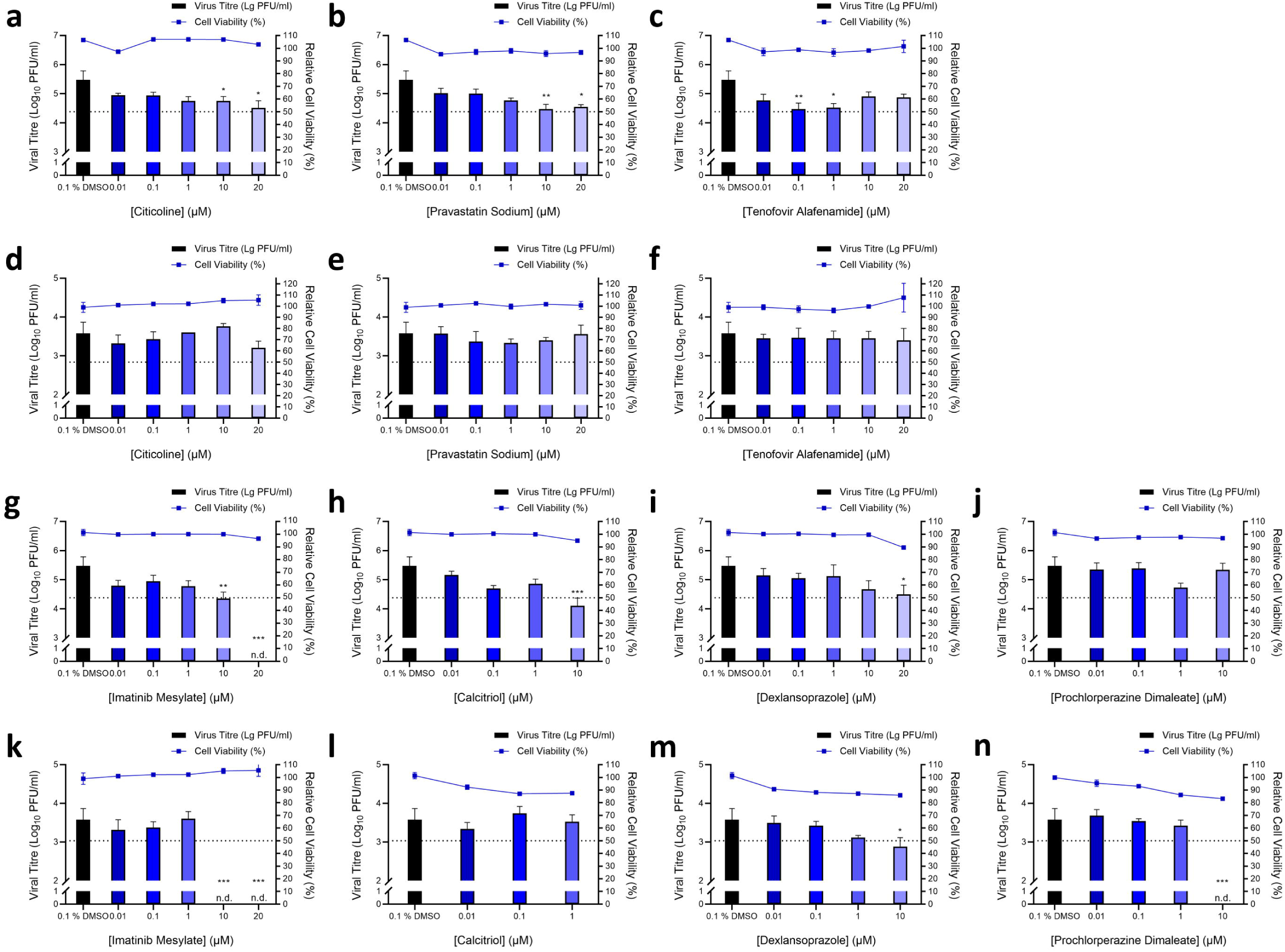
Validation of primary hits. For validating compounds with activity pre-infection, Vero E6 cells were first pre-treated with increasing concentrations of (a) citicoline, (b) pravastatin sodium and (c) tenofovir alafenamide prior to infection with SARS-CoV-2. Similarly, Huh7 cells were also pre-treated with (d) citicoline, (e) pravastatin sodium and (f) tenofovir alafenamid and subsequently infected with SARS-CoV-2. For post-infection treatment validation, Vero E6 cells were infected with SARS-CoV-2 and treated with increasing concentrations of (g) imatinib mesylate, (h) calcitriol, (i) dexlansoprazole and (j) prochlorperazine dimaleate. Similarly, HuH7 cells were also infected and treated with a range of concentrations of (k) imatinib mesylate, (l) calcitriol, (m) dexlansoprazole and (n) prochlorperazine dimaleate. The primary and secondary exes correspond to the viral titre and relative cell viability, and the dashed line represents the CC_50_ cut-off for cell viability. * denotes *p* < 0.05, ** *p* < 0.01 and *** *p* < 0.001. Error bars represent the standard deviation observed from the means of triplicates performed for both cell viability and dose-dependent inhibition studies.

**Figure 3:**
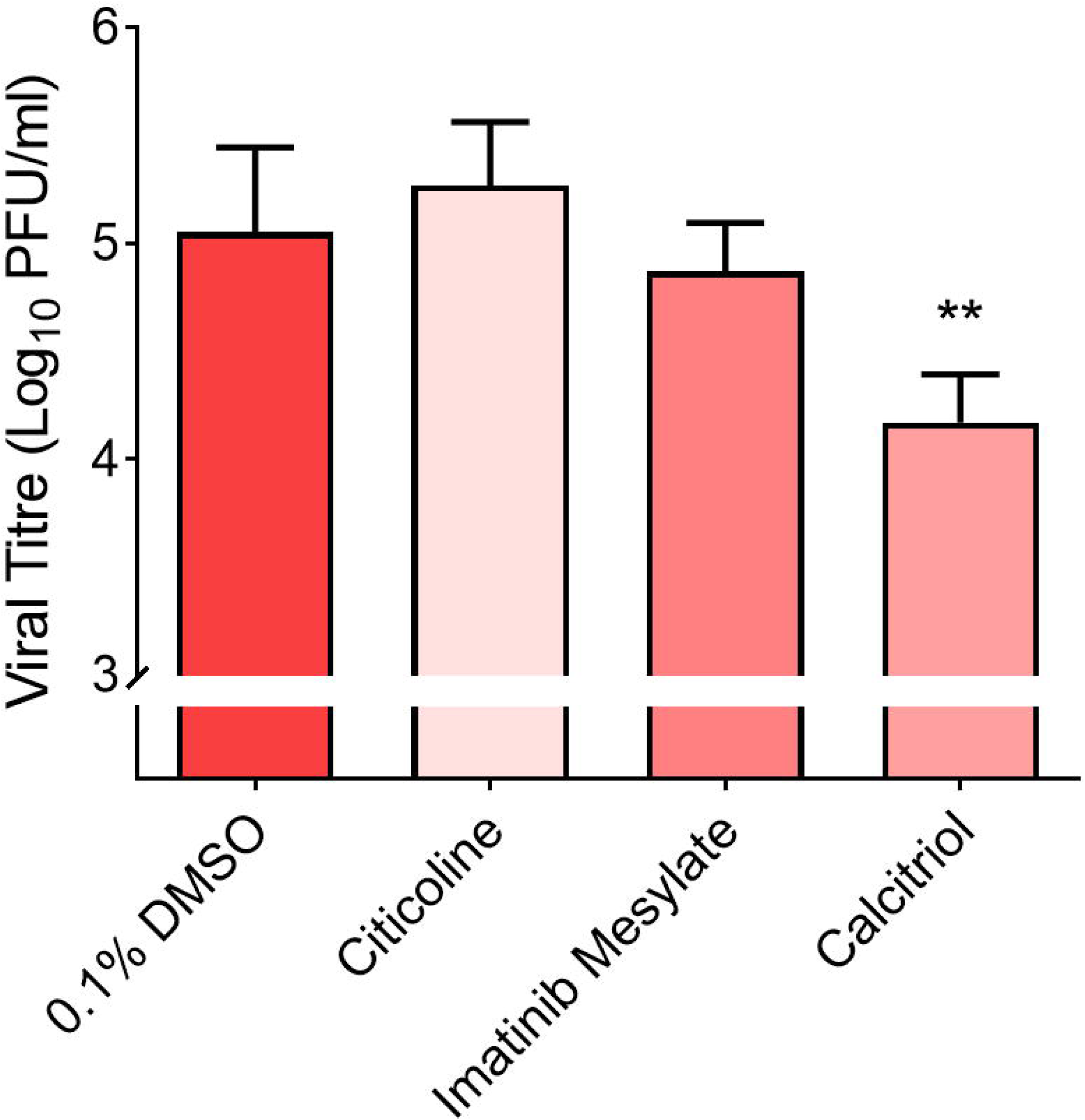
Validation of primary hits in hNECs. hNECs were either pre-treated with 10 μM citicoline prior to SARS-CoV-2 infection or treated following SARS-CoV-2 infection with 10 μM imatinib mesylate and 10 μM calcitriol. ** denotes that *p* < 0.01. Error bars represent the standard deviation observed from the means of triplicates performed for dose-dependent inhibition studies.

Vitamin D is well known to modulate host immune responses through the production of the antimicrobial peptides such as cathelicidin to promote autophagy (*15*). It has proven essential for host defenses against many intracellular pathogens including respiratory pathogens such as *Mycobacterium tuberculosis*, and has been shown to also possess anti-inflammatory properties(*15*). A recent study by Smith and colleagues (*16*) showed an association between vitamin D deficiency and SARS-CoV-2 infection and COVID-19 associated mortality. The authors speculated that vitamin D supplementation could protect against SARS-CoV-2 infection and improve patient disease outcomes(*16*), and our finding certainly provides credence to this hypothesis. Given that calcitriol-mediated inhibition occurred upon post-treatment of Vero E6 cells and hNECs, it is likely that its mechanism of antiviral action targets the post-entry phase of viral replication.

Use of Host-directed therapies (HDTs) for prevention of infections is certainly not a new idea. Most of these therapies however rely mainly on the use of vaccines, convalescent plasma and monoclonal antibodies (*17, 18*). Small molecule HDTs have been used adjunctively for diseases such as tuberculosis (*19*) and have been proposed for viral pandemics (*5*). This strategy would overcome some of the costs and challenges associated with antiviral production, including the emergence of drug resistance (*20*). Vitamin D is a dietary supplement that is cheap and widely available even in low and middle income countries and is converted by the liver and kidneys into the active compound calcitriol (*21, 22*). It is however important that our findings be confirmed in vivo as well as in clinical trials in order to assess efficacy, optimal dosage, treatment duration, toxicity and safety of calcitriol. Given the high transmissibility of SARS-CoV-2 globally (*23*), if these findings can be replicated in clinical trials, calcitriol may certainly prove to be an effective tool in the effort to control the pandemic while waiting for an effective vaccine to be rolled out globally. At the very least, these findings certainly pave the way for consideration of host-directed therapies for ring prophylaxis of contacts of SARS-CoV-2 patients.

## Methods

### Cell lines and Viruses

African green monkey kidney cells (Vero E6; ATCC CRL-1586™), human hepatoma cells (HuH7; Dr Priscilla Yang, Harvard Medical School, USA) and human nasal epithelial cells (hNECs; Dr Wang De Yun, National University of Singapore, Singapore) were utilised in this study. Vero E6 and HuH7 cells were cultured in Dulbecco’s Modified Eagle’s Medium (DMEM) (Sigma-Aldrich) supplemented with 10% heat-inactivated fetal calf serum (FCS) and buffered with 2 g sodium hydrogen carbonate.

hNECs were derived from in vitro differentiation of human nasal epithelial stem/progenitor cells (hNESPCs) obtained from healthy adult donors scheduled for septal plastic surgery. Approval to collect the tissue biopsies was obtained from the National Healthcare Group Domain-Specific Board of Singapore (DSRB Ref: D/11/228) and the institutional review board of the National University of Singapore (IRB Ref: 13-509). Written consent was also obtained from the donors prior to tissue biopsies collection. At the time of biopsies collection, all subjects were free of upper respiratory tract infection and rhinitis symptoms. Once the hNESPCs were isolated and enriched using standardised protocols (24, 25), the hNESPCs were expanded and subjected to ALI culture in Transwells for in vitro differentiation (24, 25). The expanded hNESPCs were then transferred onto 12-well 0.4 μm Transwell inserts (Corning). Once the cells became confluent, growth medium was discarded and 700 μl of PneumaCult™-ALI Medium with inducer supplements (STEMCELL Technologies) was added to the basal chamber to establish ALI conditions. The cells were cultured in ALI culture for 4 weeks, with media change every 2-3 days. hNECs were obtained after 3-4 weeks of differentiation were then subjected to SARS-CoV-2 infection. All cultures were maintained at 37°C, 5% CO_2_.

SARS-CoV-2 (hCoV-19/Singapore/10/2020) was isolated from a nasopharyngeal swab of a COVID-19 patient in February 2020. The virus isolate was validated by qRT-PCR and propagated in Vero E6 cells with no more than 3 passages prior to drug screening. The virus genome (EPI_ISL_410716) has been released to GISAID by the National Public Health Laboratory, National Centre for Infectious Diseases, Singapore and belongs to Clade O. All virus work was performed in a biosafety level 3 (BSL-3) laboratory and all protocols were approved by the BSL-3 Biosafety Committee and Institutional Biosafety Committee of the National University of Singapore.

### Preparation of compound libraries for primary screen

A primary screen to identify novel compounds with potential antiviral effects against SARS-CoV-2 infection was performed on four different libraries, namely, a 462-compound ACE2-targeted compound library (CADD) (TargetMol), 57-compounds natural product library, 500-compound flavonoids library (TimTec) and 1172-compound FDA-approved drug library (Selleckchem). Compounds were first dissolved in 100% dimethyl sulfoxide (DMSO) to a stock concentration of 10 mM, followed by dilution to 100 μM in serum-free media. Compounds were stored at -20°C till further use.

### Primary screen with pre-treatment using ACE2-targeted inhibitors and 57-compound natural product libraries

Vero E6 cells were seeded onto 96-well plates (Corning) at a seeding density of 1×10^4^ cells per 100 µl and incubated overnight prior to drug inhibitory assay. Pre-infection treatment drug assays were performed by treating cells with 10 µM of ACE2-targeted inhibitors and the 57-compound natural product libraries for 2 h at 37°C. 0.1% DMSO and 100 μM remdesivir were included as vehicle and positive controls, respectively. Treated cells were then washed twice with phosphate-buffered saline (PBS) prior to SARS-CoV-2 infection at a multiplicity of infection (MOI) of 1 and incubated for 4 days at 37°C, 5% CO_2_, before formalin fixation, analysis and hit selection.

### Primary screen with post-treatment using FDA-approved drugs and flavonoids library

Vero E6 cells were seeded onto 96-well plates (Corning) at a seeding density of 1×10^4^ cells per 100 µl and incubated overnight. Cells were infected with SARS-CoV-2 infection at a MOI of 1 for 1 h. Final concentrations of 10 µM of FDA-approved drugs and 50 µM of flavonoids were added to SARS-CoV-2-infected cells, respectively. 0.1% DMSO and 100 μM remdesivir were included as vehicle and positive controls, respectively. Cells were then incubated for 4 days at 37°C, 5% CO_2_ before formalin fixation, analysis and hit selection.

### Selection of hits via analysis of cytopathic effects (CPE)

For selection of hits at 4 days post infection, differences in cell viability caused by virus-induced CPE and/or compound-specific toxicity were visually analysed using light microscopy (Olympus). Hits were identified and selected based on the reduction of CPE and > 50% inhibition in duplicate wells in comparison to vehicle control.

### Validation of hits

Hits identified from the primary screens were validated using cell viability and dose-dependent inhibition assays on Vero E6 and HuH7 cells. To determine the cell viability profiles of the selected compounds, Vero E6 and HuH7 cells were seeded into 96-well plates at seeding densities of 1×10^4^ cells and 7.5×10^3^ cells per 100 µl, respectively. Both cell lines were pre-treated with the hit compounds from the ACE2-targeted library, namely citicoline, pravastatin sodium and tenofovir alafenamide for 2 h at a range of concentrations (0.01, 0.1, 1, 10 and 20 µM). Hits identified from the FDA-approved drug library, namely, imatinib mesylate, calcitriol, dexlansoprazole and prochlorperazine dimaleate were added to the cells with the same concentration range and incubated for 4 days. After incubation, the media was removed from the plates and washed once with PBS before addition of alamarBlue Cell Viability Reagent (Thermo Fisher Scientific) diluted 1:10 in media with 2% FCS. Fluorescence was detected after 2.5 h incubation at an excitation wavelength of 570 nm and an emission wavelength of 600 nm on an Infinite 200 Pro multiplate reader (Tecan). Data obtained from compound-treated cells and vehicle-treated cells were normalised against those obtained from untreated cells.

Validation of the primary hits was performed via dose-dependent inhibition assays. Vero E6 and HuH7 cells were seeded onto 96-well plates and incubated overnight prior to drug inhibitory assays. Pre-infection treatment with citicoline, pravastatin sodium and tenofovir alafenamide at working concentrations of 0.01, 0.1, 1, 10, 20 µM was carried out for 2 h at 37°C. The inhibitors were removed and washed twice with PBS prior to SARS-CoV-2 infection at an MOI of 1. For validation of imatinib mesylate, calcitriol, dexlansoprazole and prochlorperazine dimaleate, Vero E6 and HuH7 cells were infected with SARS-CoV-2 at an MOI of 1 for 1 h at 37°C and then incubated with the compounds at working concentrations. All dose-dependent inhibition assay plates were incubated for 4 days at 37°C, 5% CO_2_, prior to harvesting of viral supernatants for virus titration.

### Validations of imantib mesylate, calcitrol and citicoline with primary cell line, hNEC at 10 µM

hNECs were seeded in apical chambers of the Transwell inserts for 4 weeks prior to drug inhibitory assays. Pre-infection treatment with citicoline at 10 µM was added to the basal chamber and carried out for 2 h at 37°C. The inhibitors were removed and replaced with fresh PneumaCult™-ALI Medium (STEMCELL Technologies) prior to SARS-CoV-2 infection at an MOI of 0.1. For validation of imatinib mesylate and calcitriol, hNECs were infected with SARS-CoV-2 at a MOI of 0.1 for 1 h at 37°C before incubation with the compounds at 10 µM. All inhibition assay plates were incubated for 4 days at 37°C, 5% CO_2_, prior to harvesting of viral supernatants for virus titration.

### Plaque Assay

To determine the virus titer, viral supernatants harvested were 10-fold serially diluted in DMEM. 200 µl of each serial diluted supernatant were applied to confluent Vero E6 cells. After 1 h of absorption, the inoculum was removed and 500 µl of 0.5% agarose overlay was added to each well and incubated for 3 days at 37°C, 5% CO_2_ to facilitate plaque formation. The cells were fixed with formalin overnight and the agarose was removed before staining with crystal violet for 5 minutes. The number of plaques were counted and the virus titer of individual samples were expressed in the logarithm of plaque forming units (pfu) per ml.

### Statistical Analyses

One-way analyses of variance (ANOVA) was used to evaluate the statistical significance of data obtained. Drug-treated samples expressing statistical difference when compared to control samples were subsequently subjected to a Dunnett’s post-test, with * denoting that *p* < 0.05, ** denoting that *p* < 0.01 and *** denoting that *p* < 0.001.

## Author contributions

Conceived and designed the experiments: CKM, JJHC. Performed the experiments: CKM, YLN, BAA, RCHL. Analyzed the data: CKM, YLN, BAA, RCHL, MWCL, PK, PT, WJC, JEW, JJHC. Contributed reagents/materials: JL, KST, DYW, EH, XH JD, TMM, CL, RL. Wrote the paper: BAA, MWCL, YLN, RCHL, PK, CKM, YWT, JJHC. All the authors read the manuscript and approved it before submission

## Acknowledgments

We are grateful to the Yong Loo Lin School of Medicine BSL-3 Core Facility for their support with this work. This work was supported through the following funding mechanisms: NUHSRO/2020/066/NUSMedCovid/01/BSL3 Covid Research Work; NUHSRO/2020/050/RO5+5/NUHS-COVID/4; Ministry of Education, Singapore, MOE Tier 2, MOE2017-T2-2-014; Singapore NMRC Centre Grant Program – Diabetes, Tuberculosis and Neuroscience (CGAug16M009); MOH-COVID19RF2-0001; Sino-Singapore Cooperation for Evaluating the Effectiveness and Application of Guangxi Zhuang/Yao Medicines Against COVID-19 (No. GUIKE AB20036001)

## Notes

### Competing Interest Statement

The authors have declared no competing interest.

