## Supplementary info for "Calcitriol, the active form of vitamin D, is a promising candidate for COVID-19 prophylaxis"

**Table S1**: List of primary hits identified from the primary screen performed using an FDA-approved library.

| **Compound/Treatment** | **Image** | **Compound/Treatment** | **Image** |
| --- | --- | --- | --- |
| Uninfected | 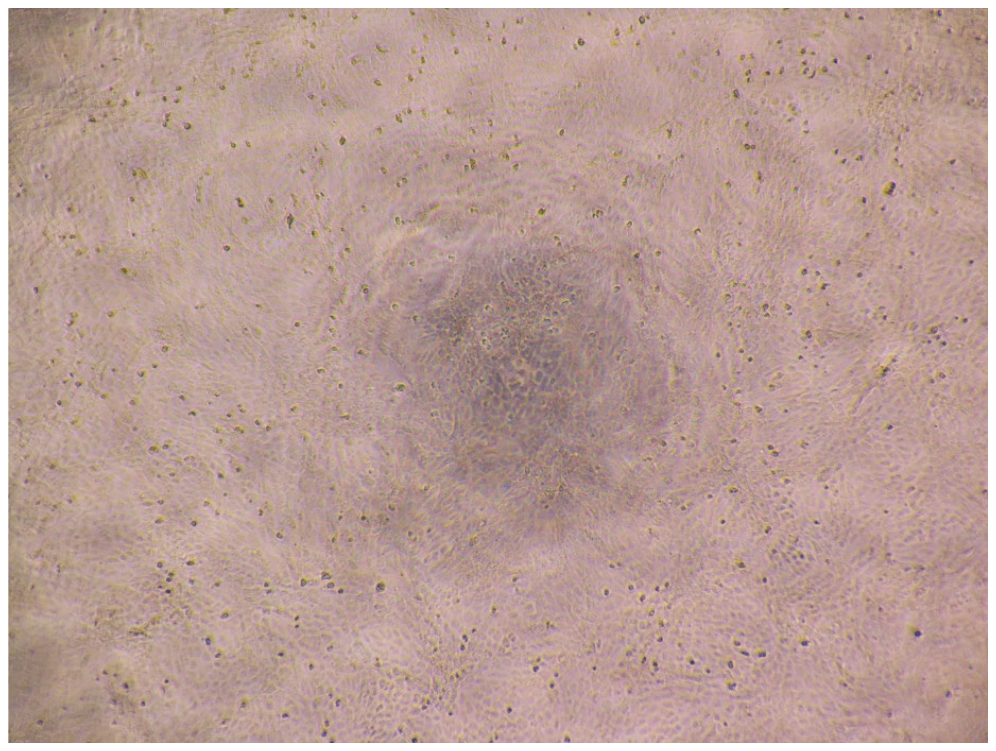 | 0.1% DMSO | 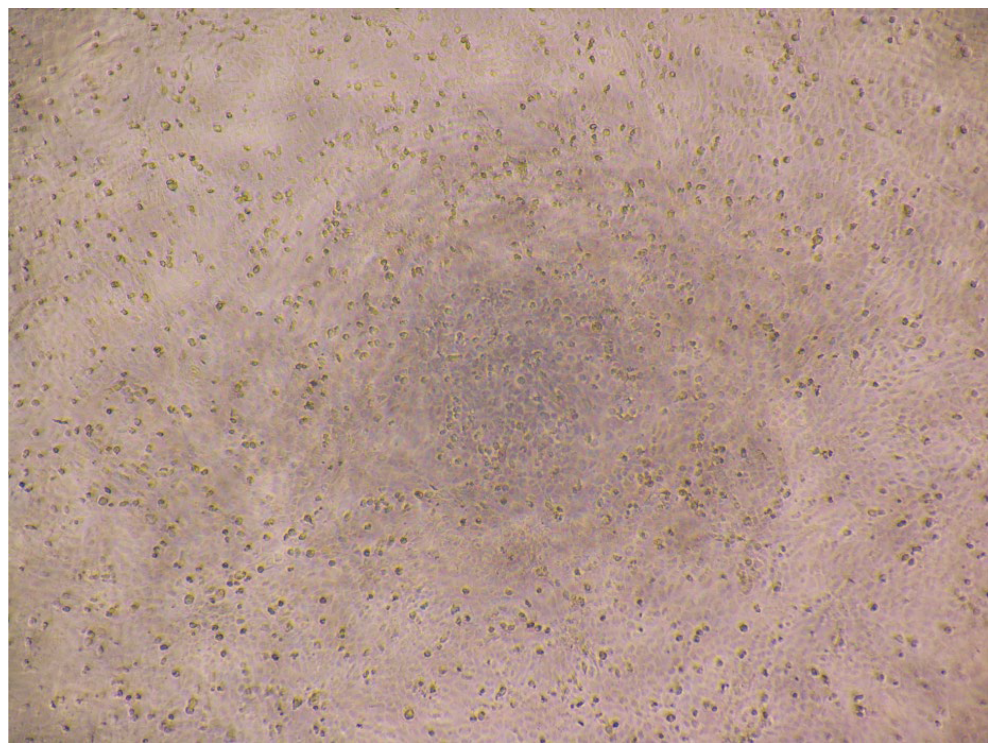 |
| Remdesivir | 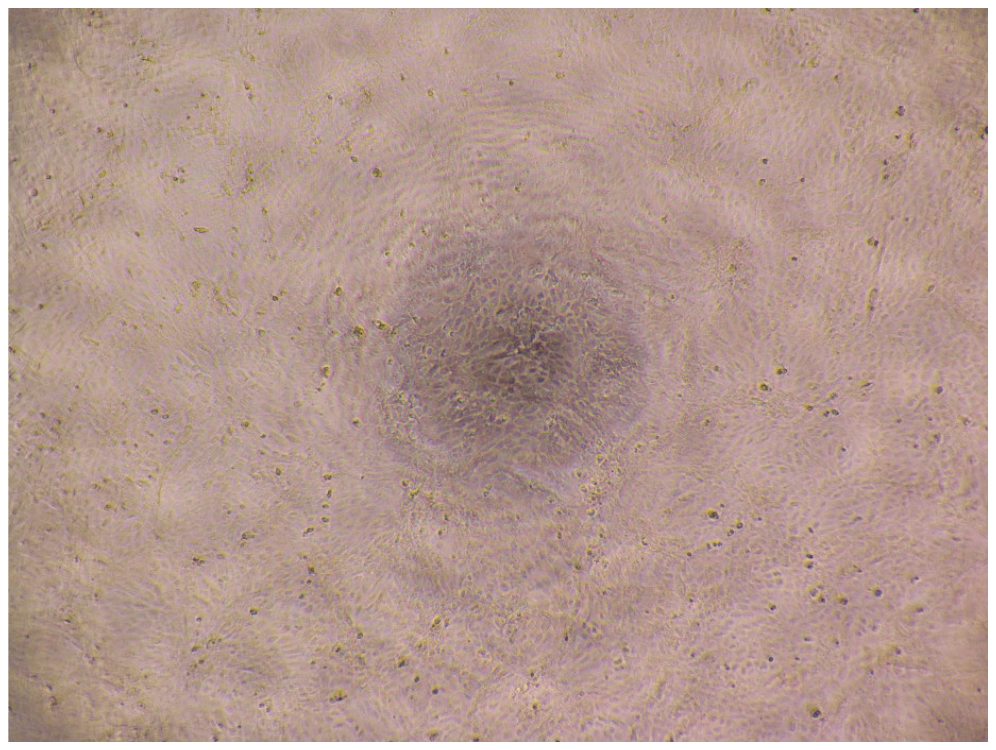 |  |  |
| Ramelteon (TAK-375) | 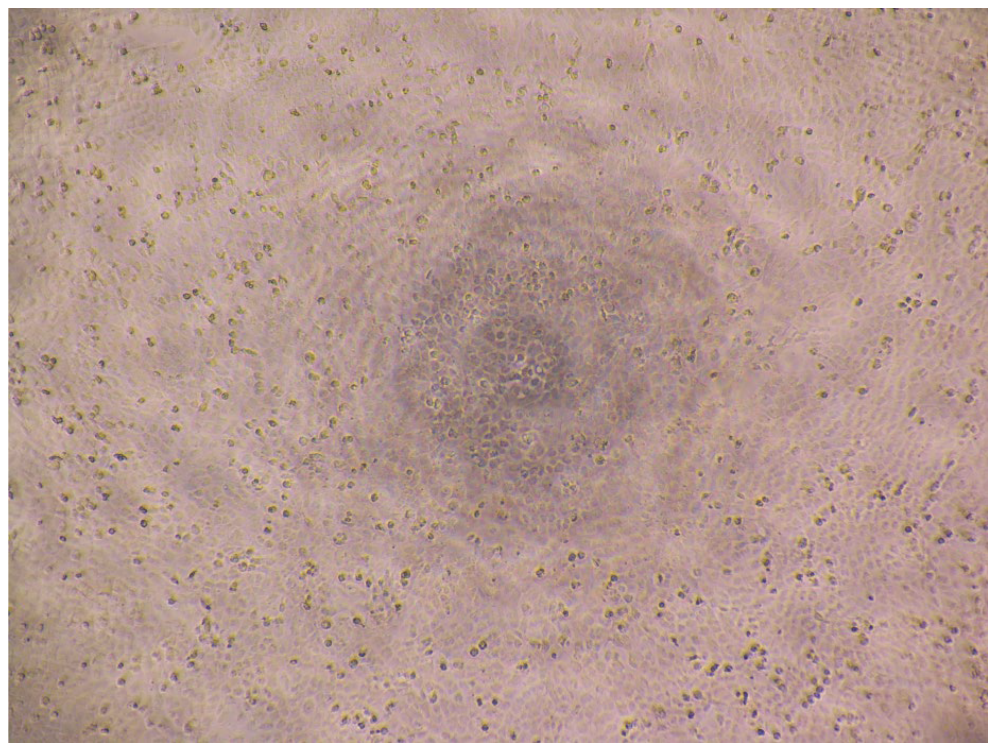 | Telmisartan | 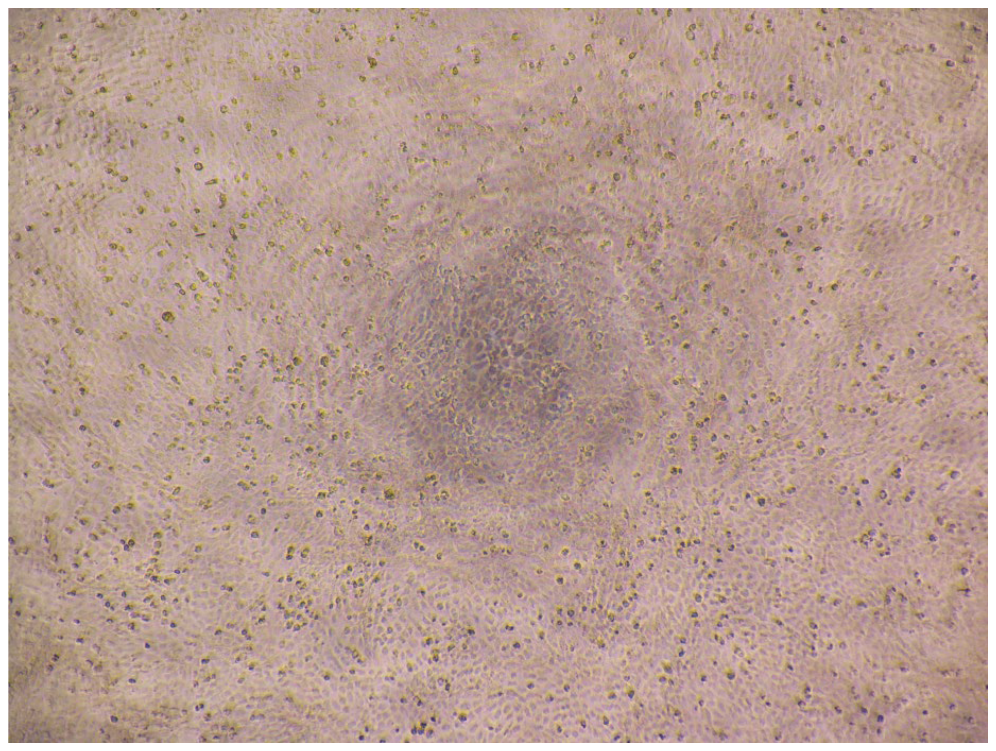 |
| Masitinib | 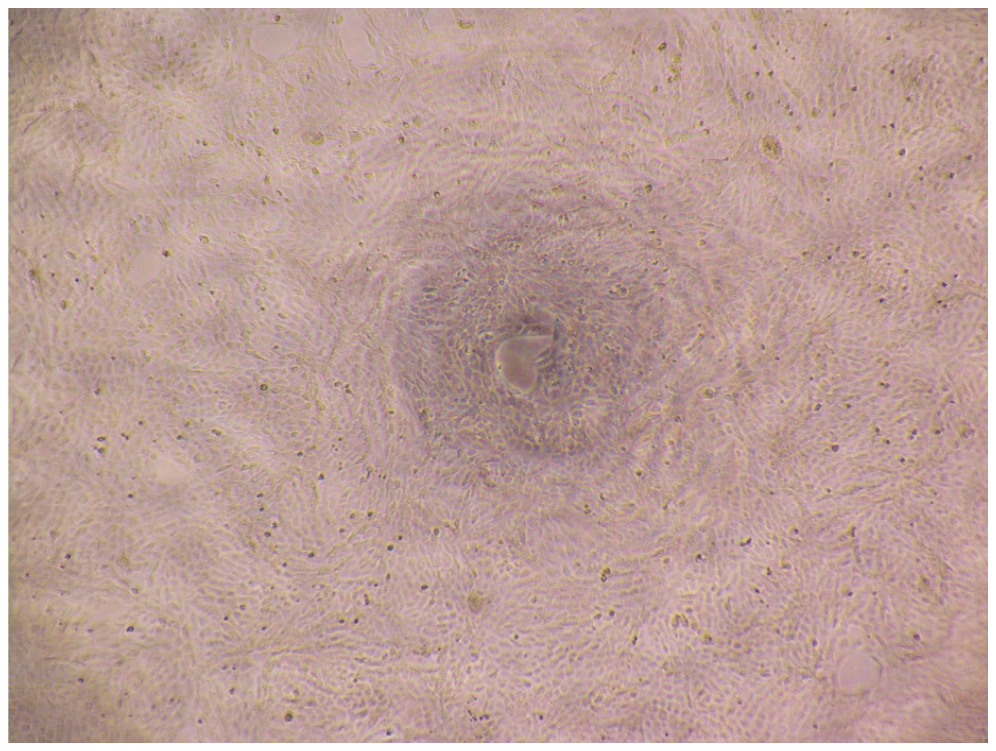 | Rifampin | 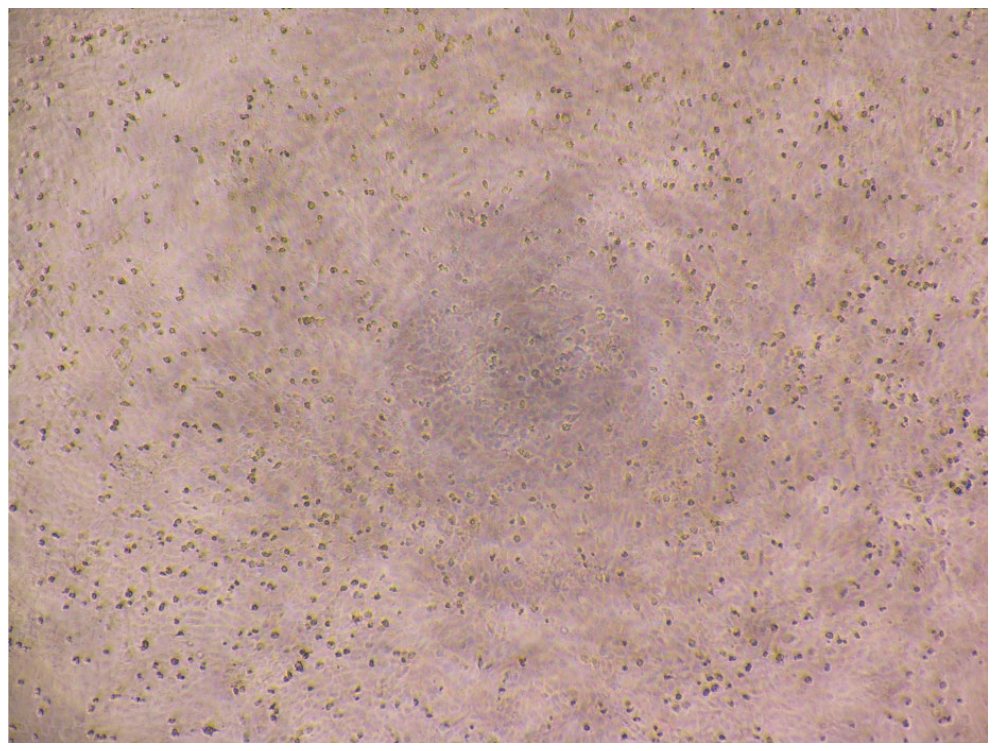 |
| MDV3100 (Enzalutamide) | 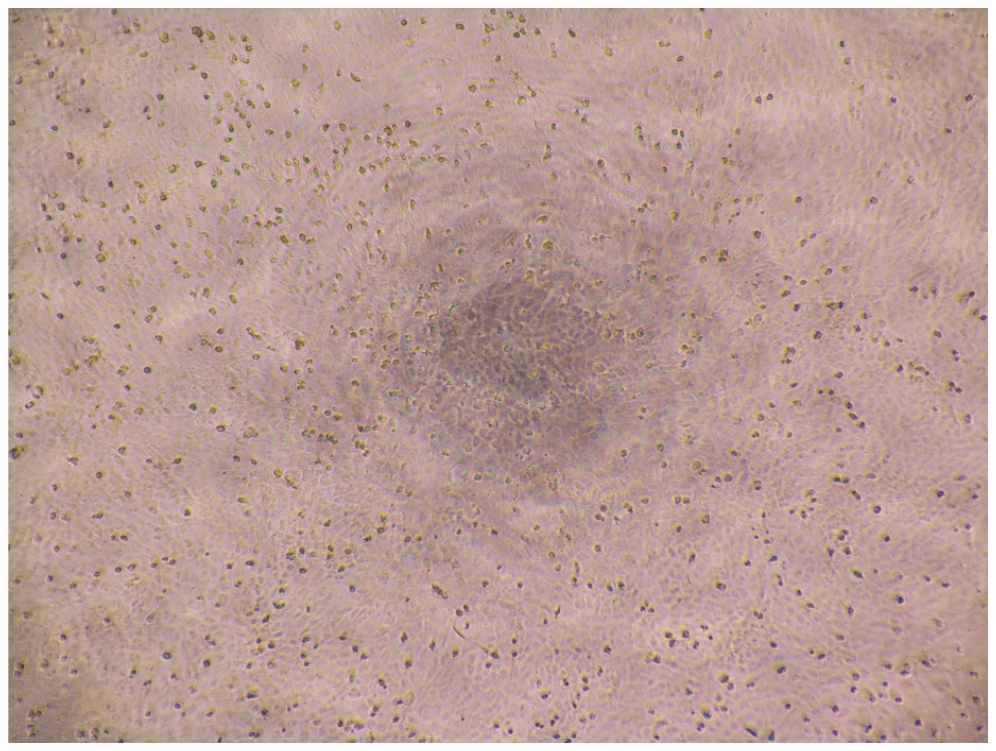 | Amorolfine Hydrochloride | 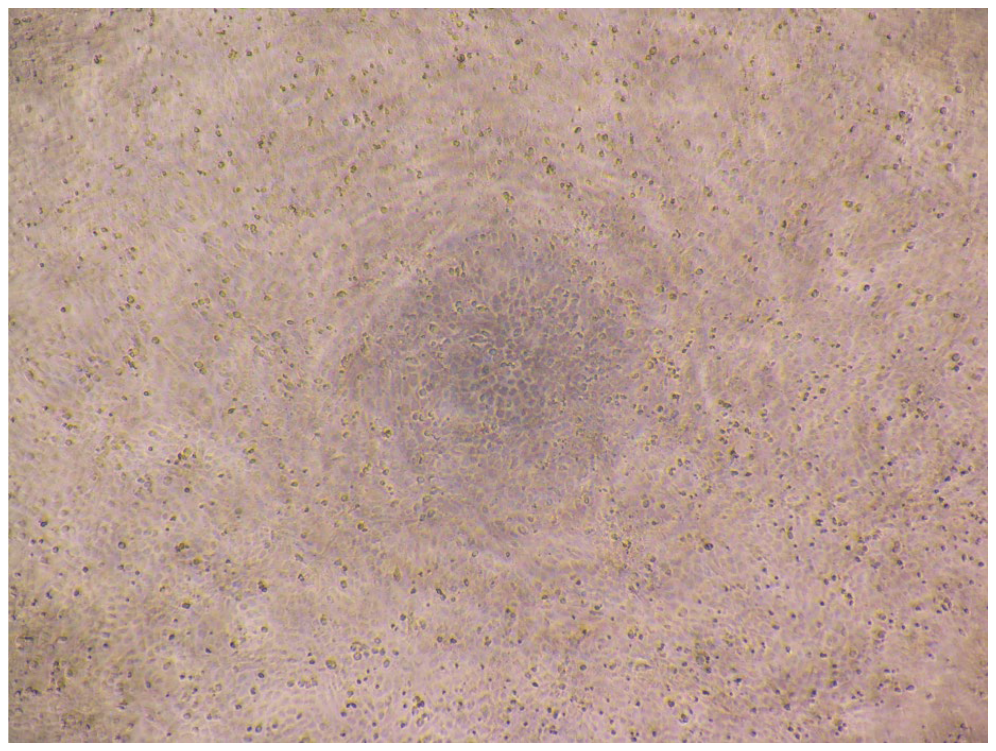 |
| Everolimus | 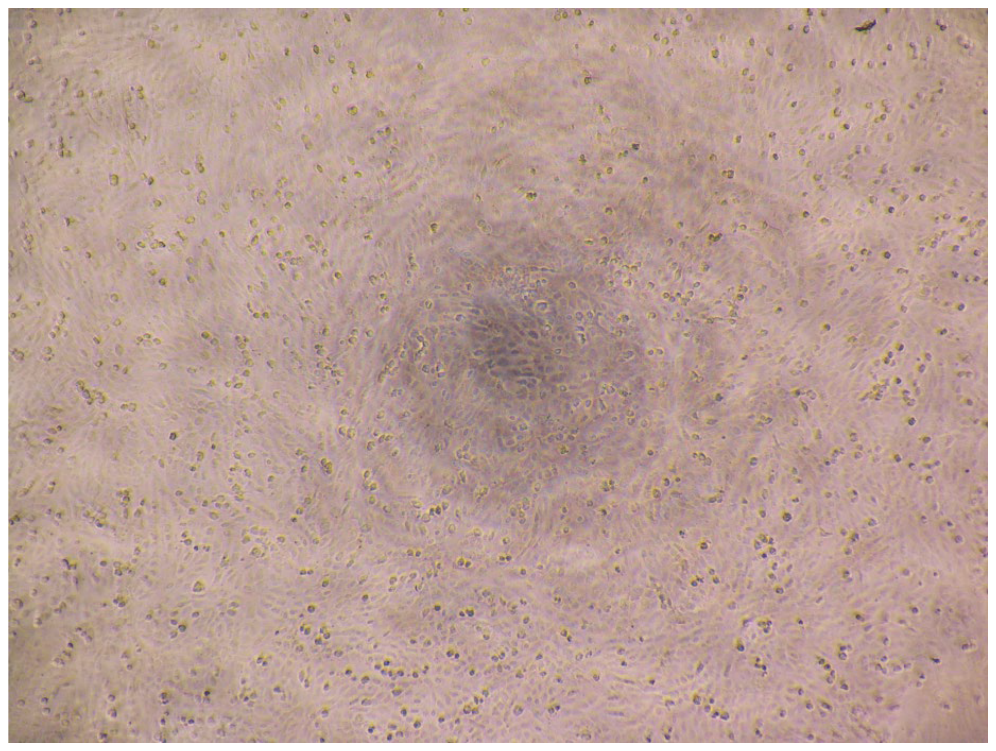 | Manidipine | 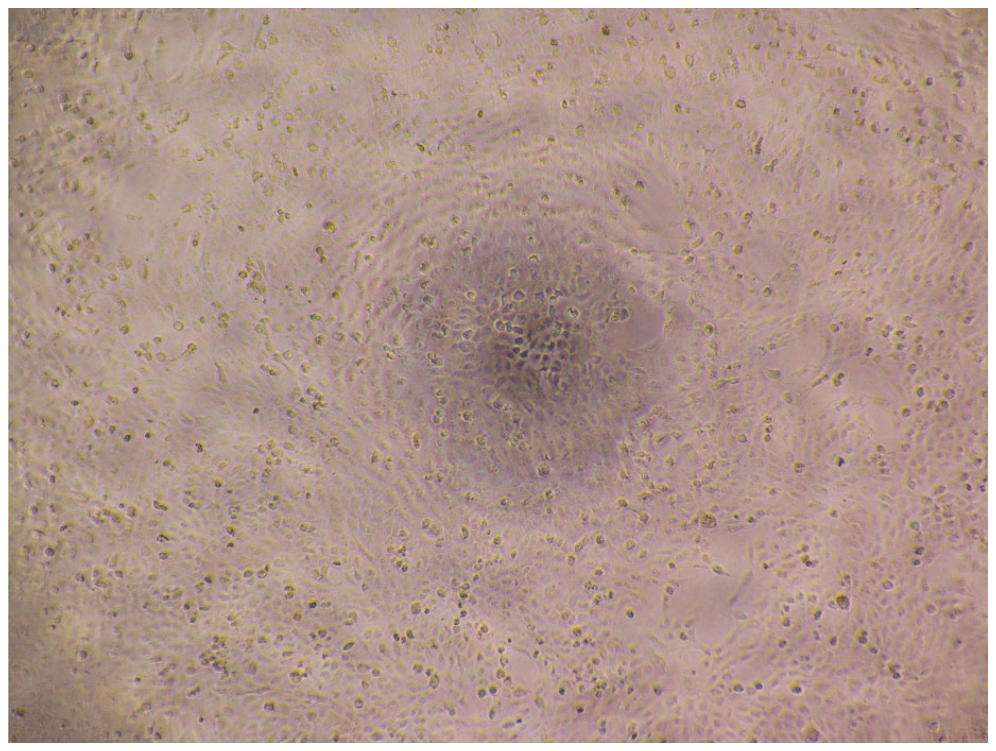 |
| Fulvestrant (Faslodex) | 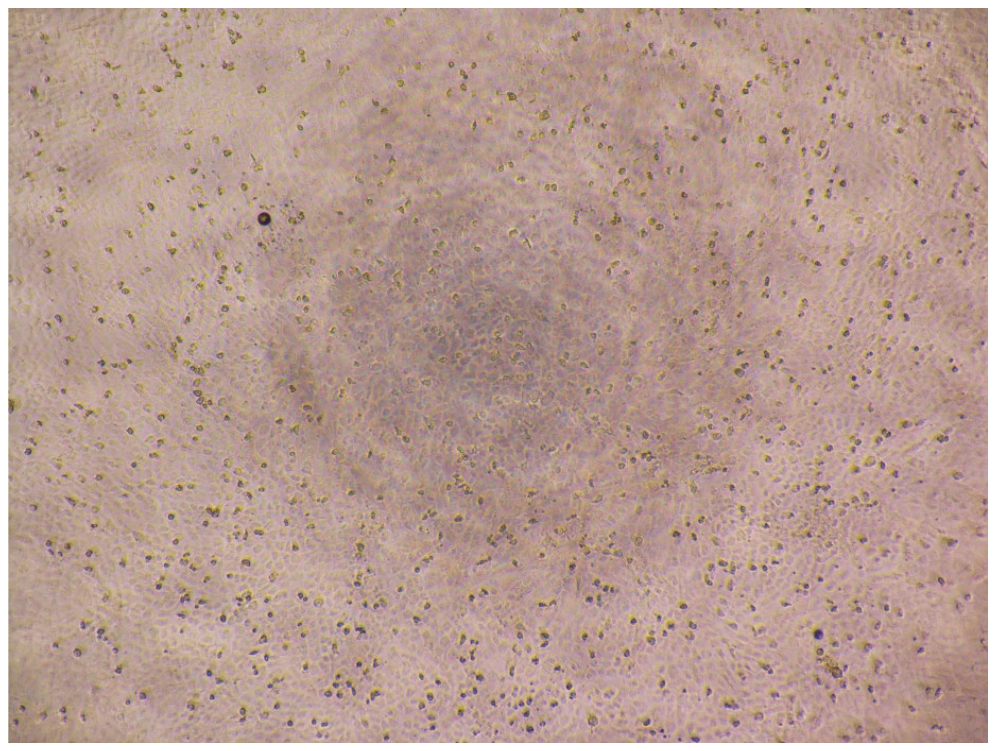 | Clomifene Citrate | 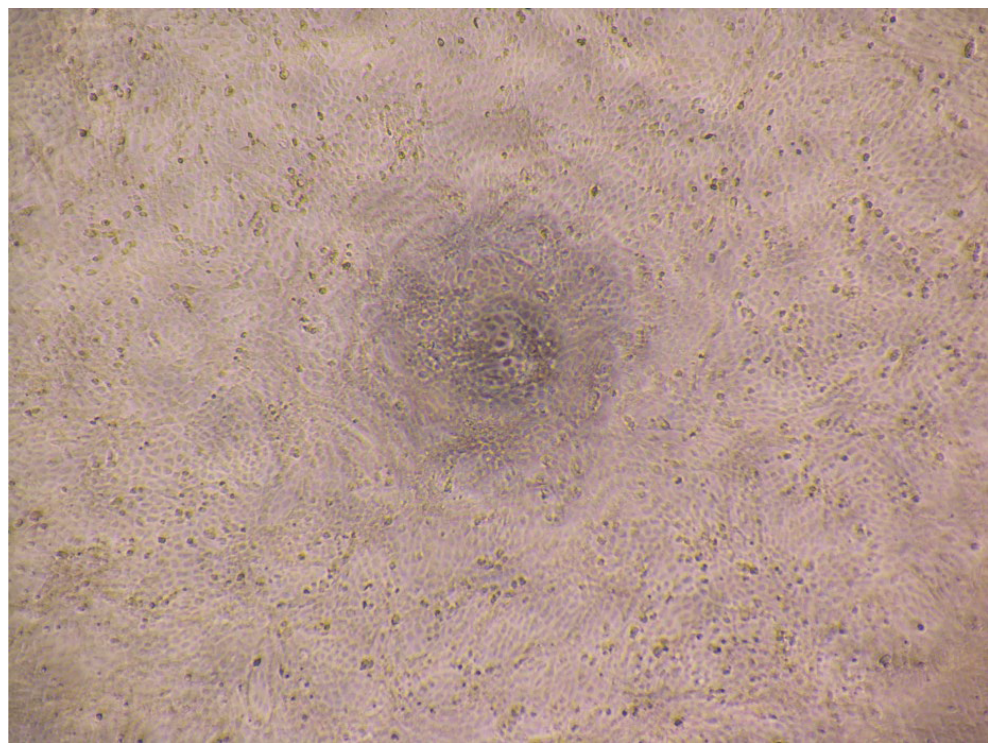 |
| Imatinib Mesylate | 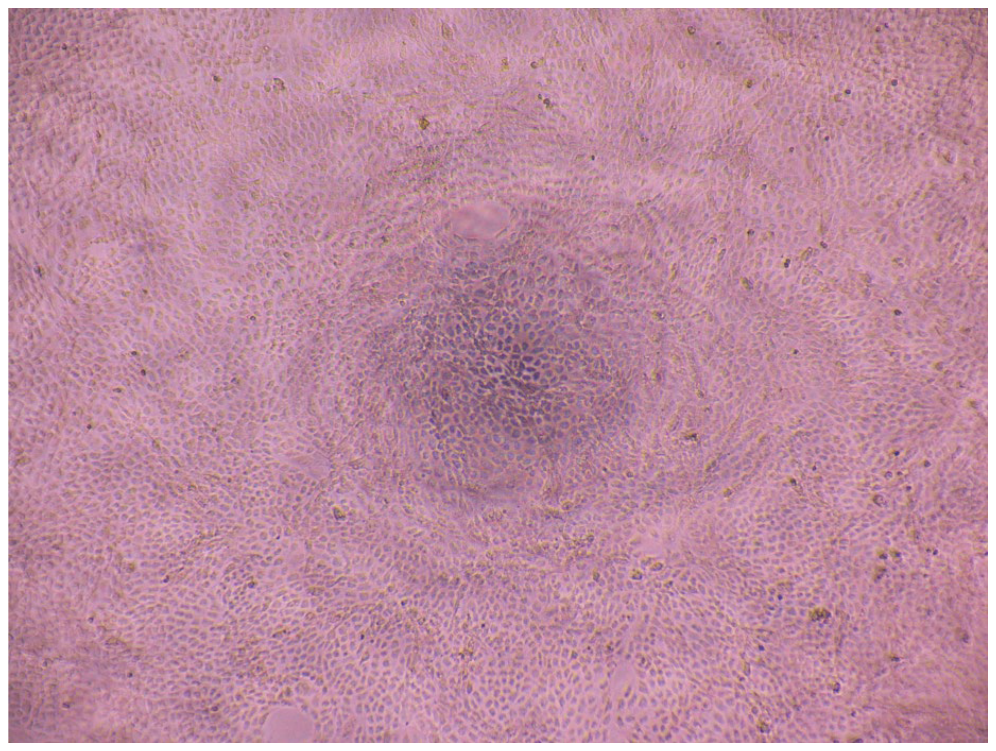 | Econazole Nitrate | 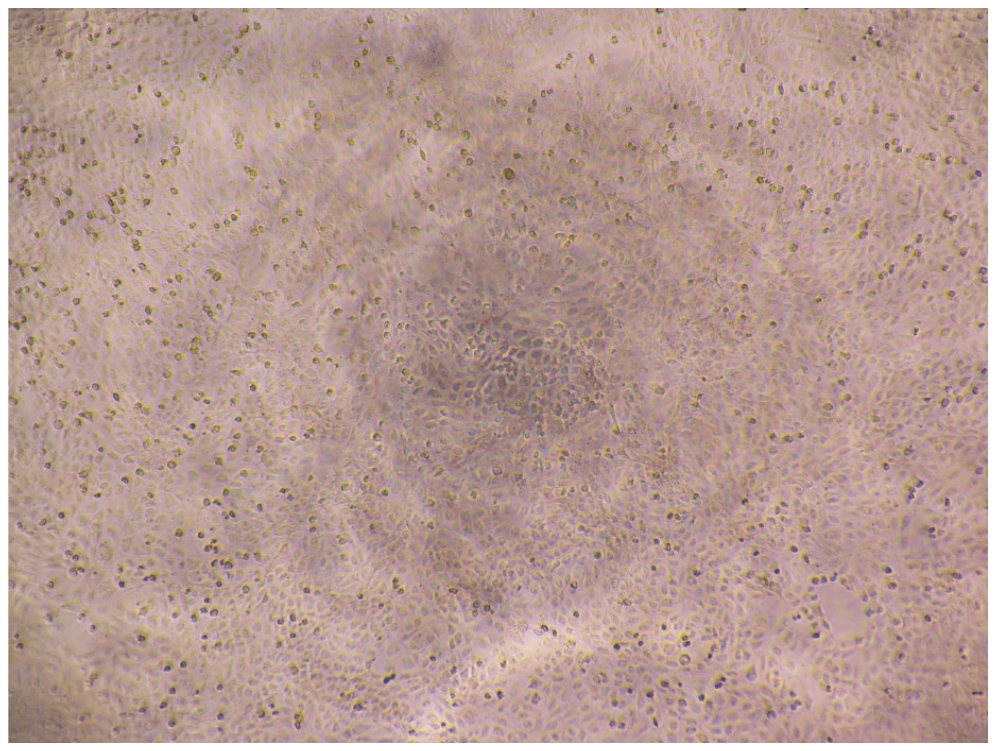 |
| Mianserin Hydrochloride | 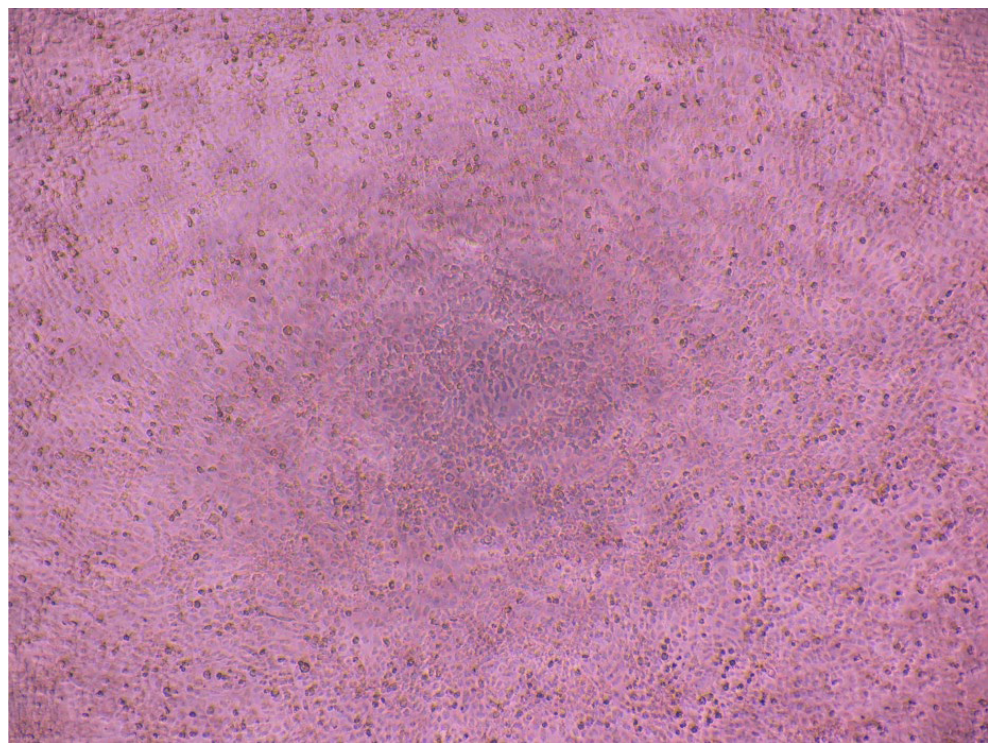 | Phenacetin | 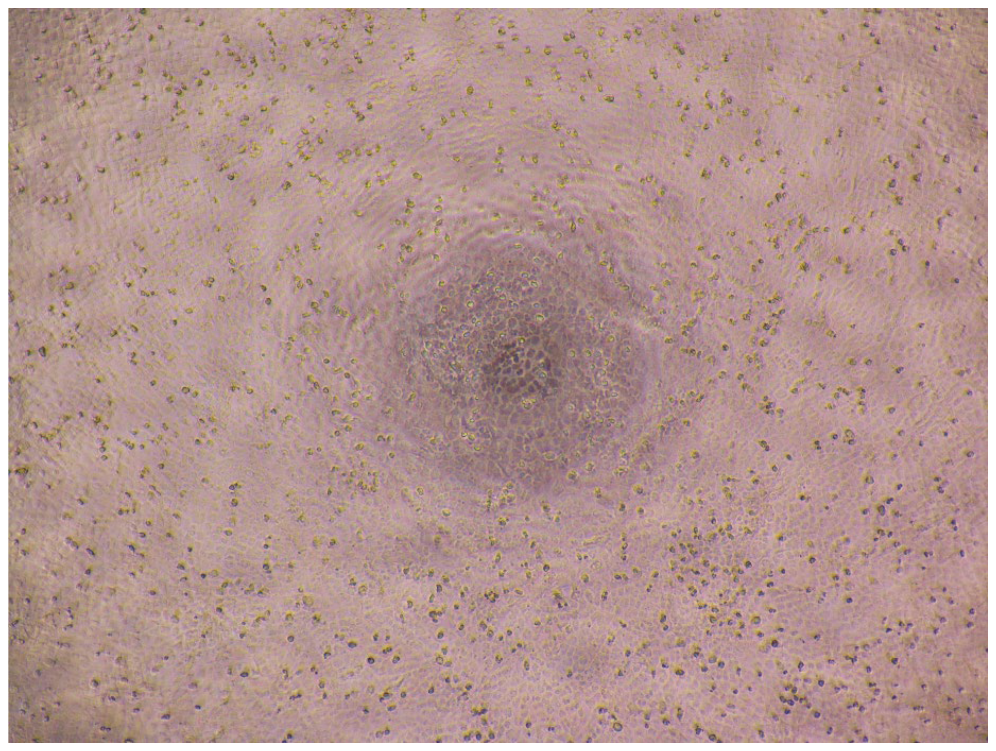 |
| Trilostane | 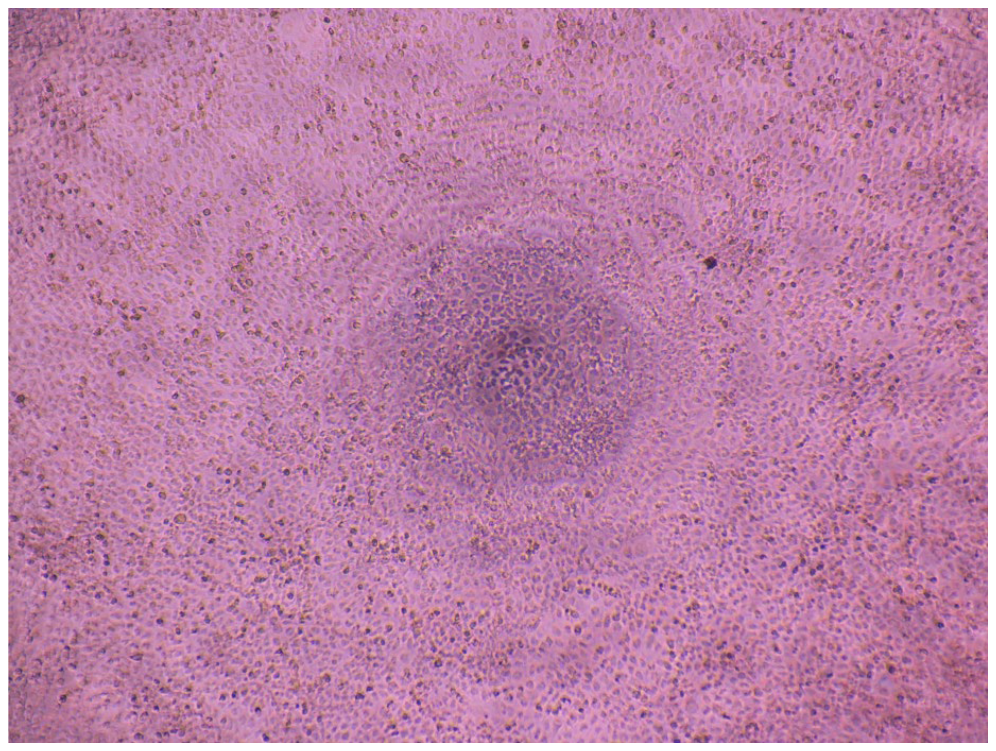 | Lonidamine | 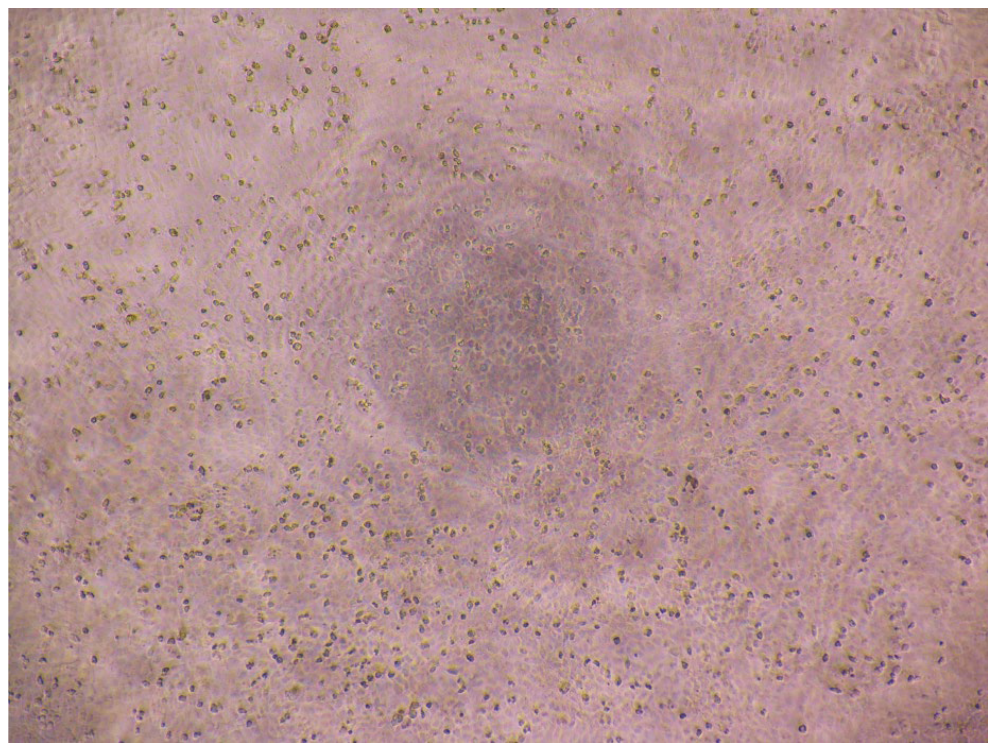 |
| Mercaptopurine | 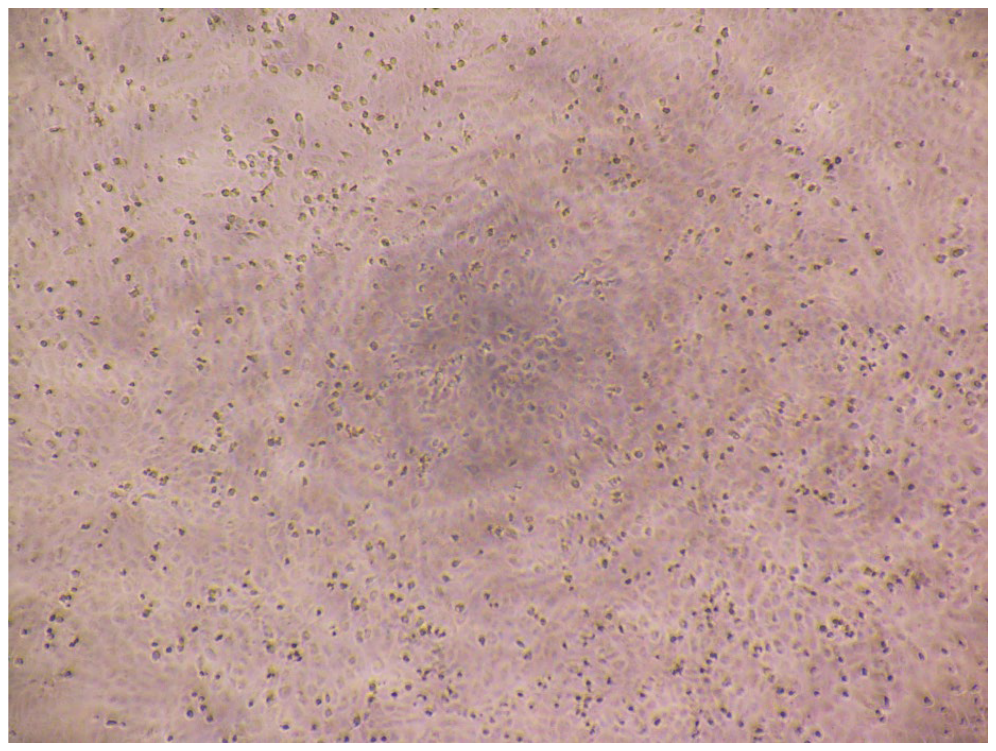 | Idebenone | 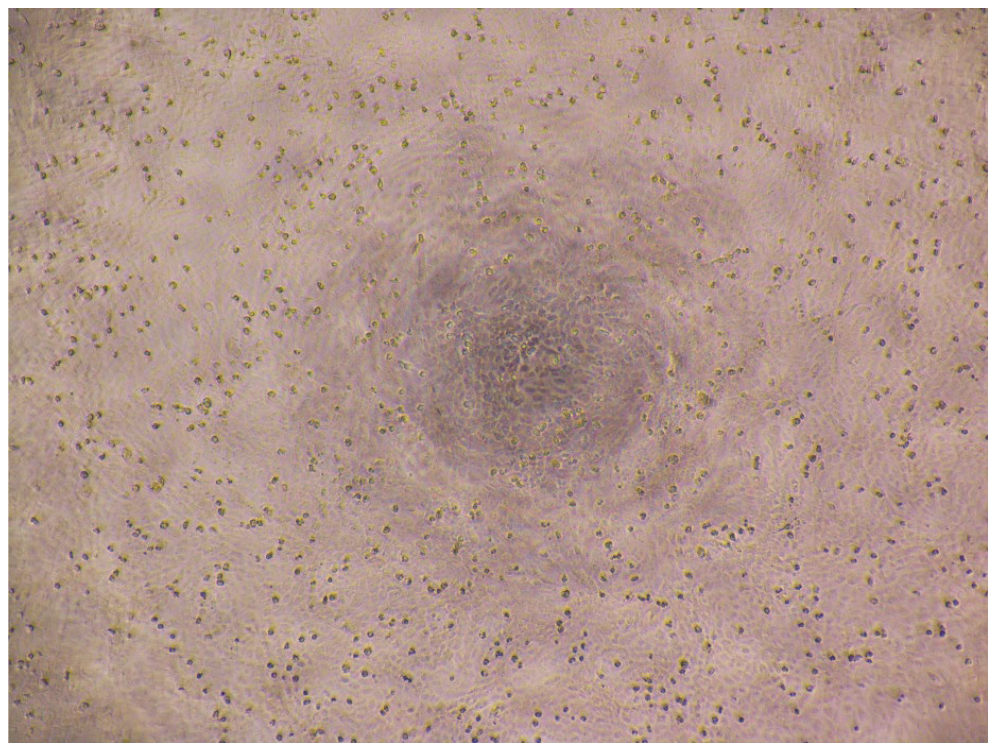 |
| Artemisinin | 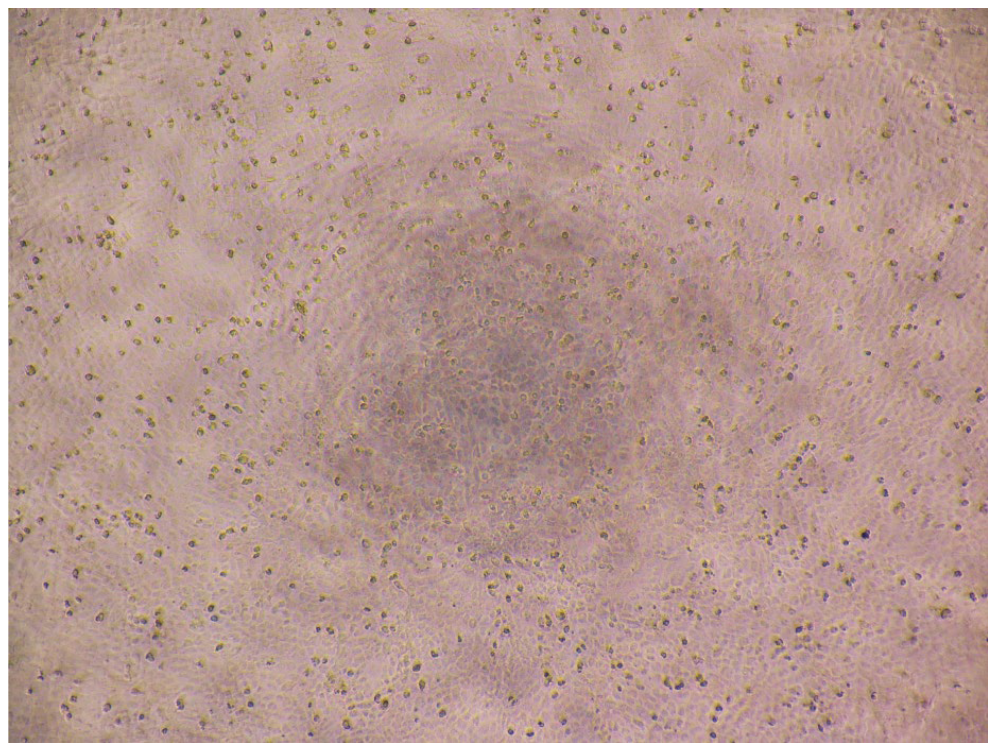 | Hygromycin B | 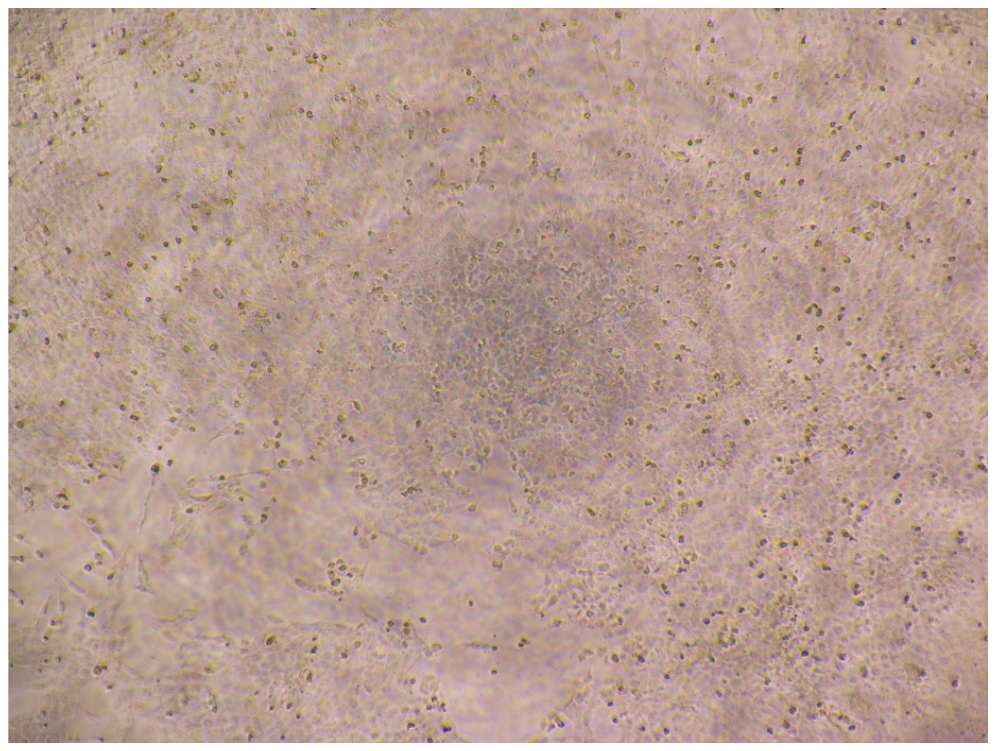 |
| Cetirizine Dihydrochloride | 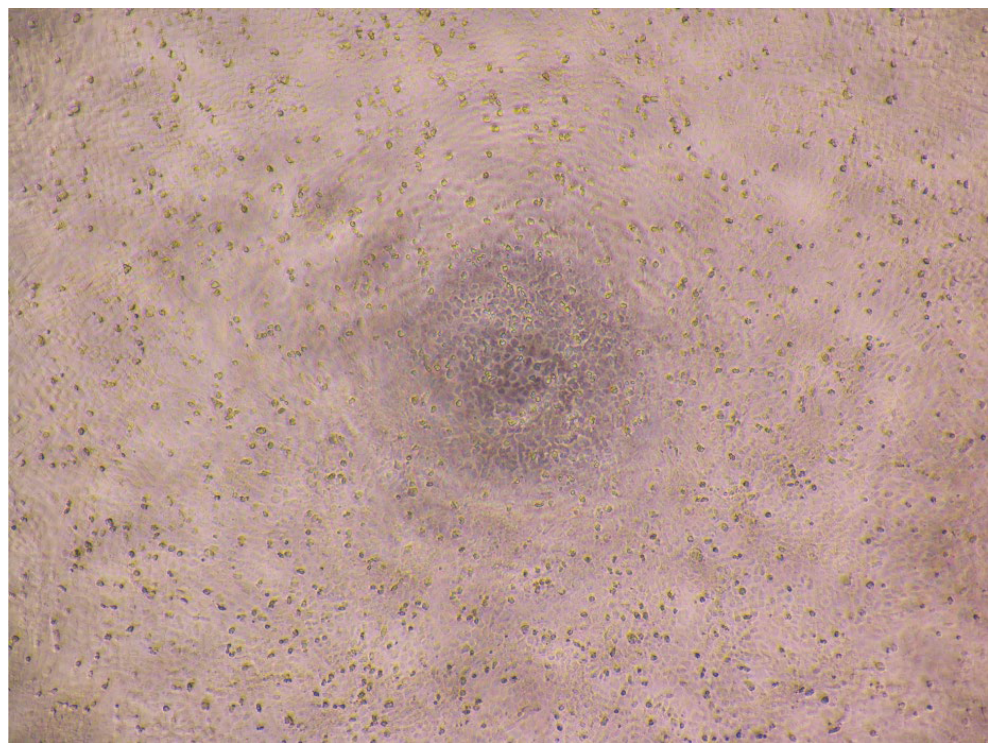 | Alverine Citrate | 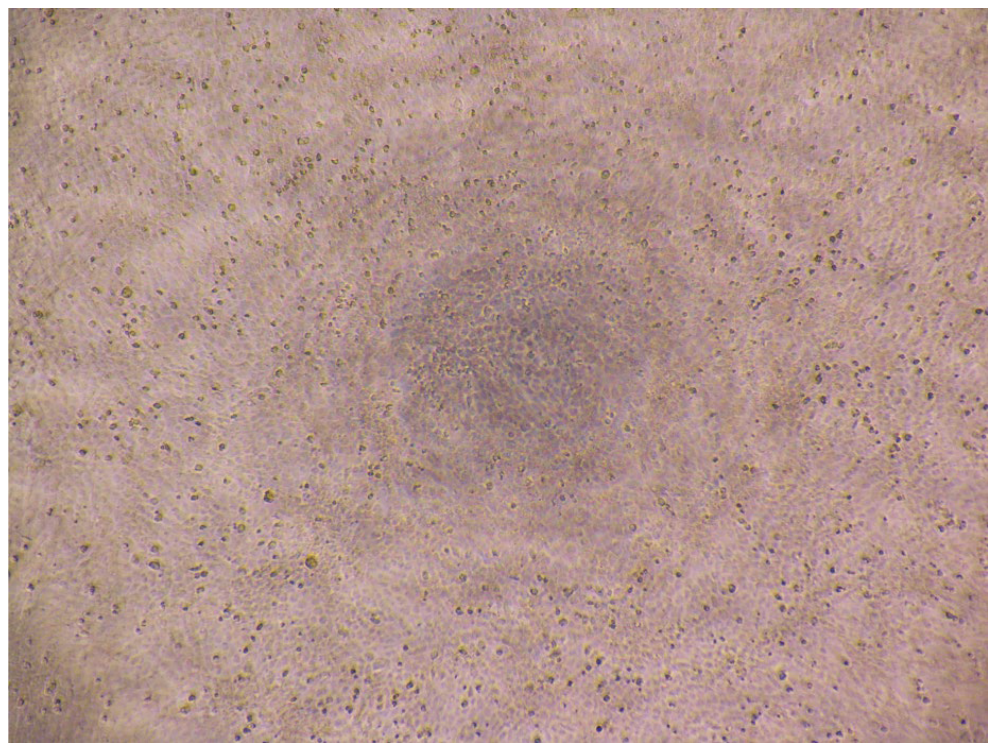 |
| Loratadine | 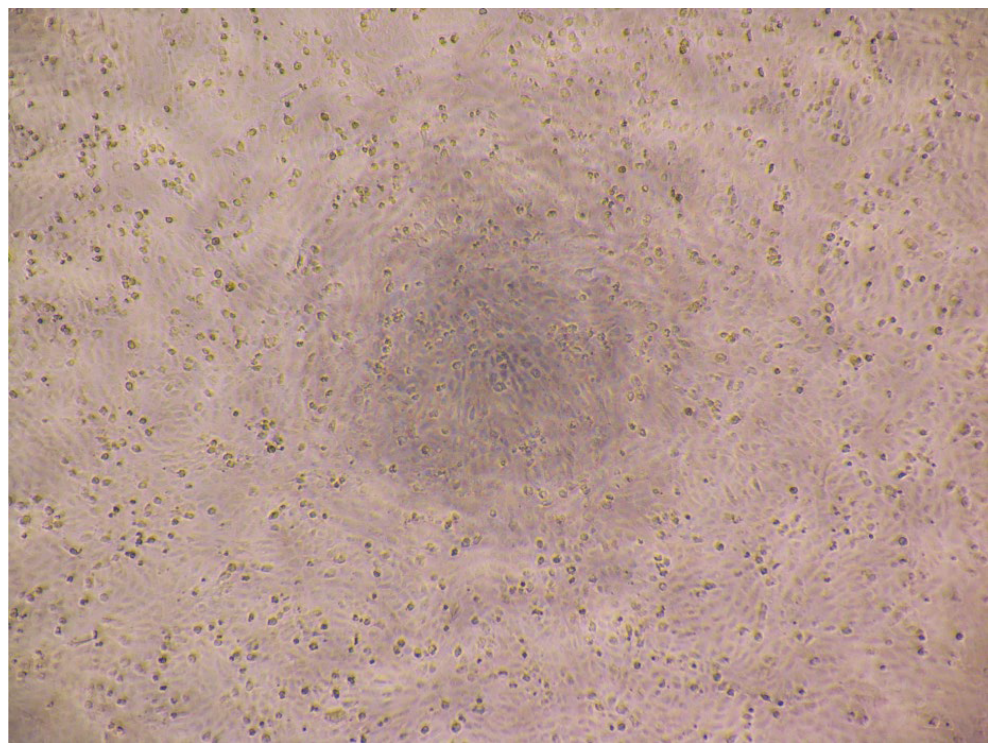 | Doxycycline Hydrochloride | 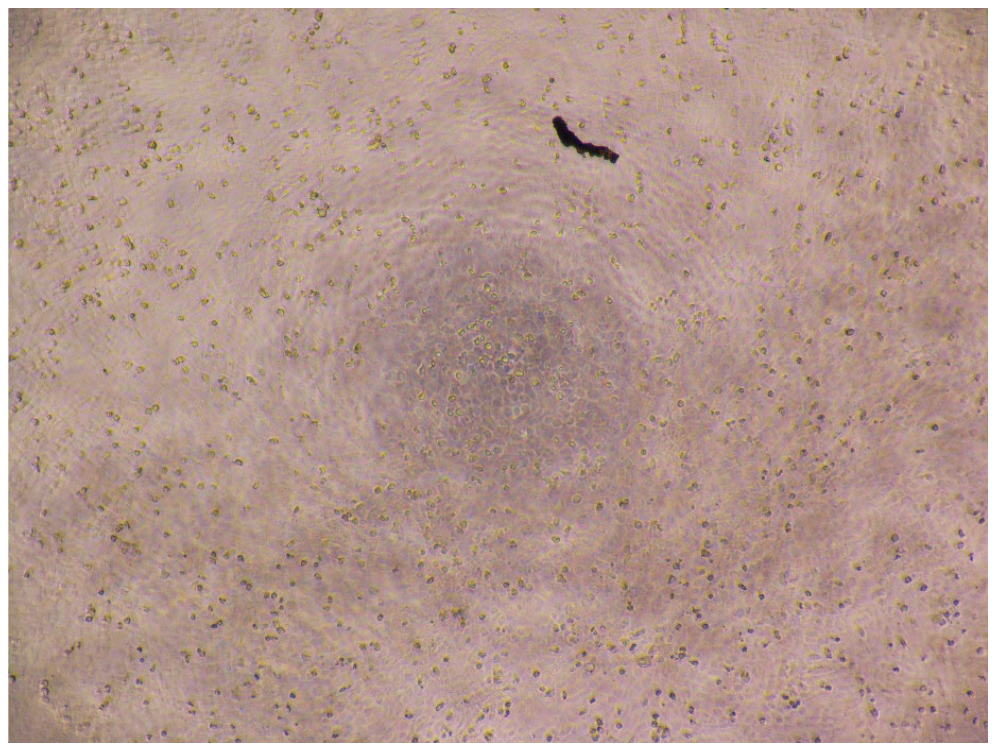 |
| Ketoconazole | 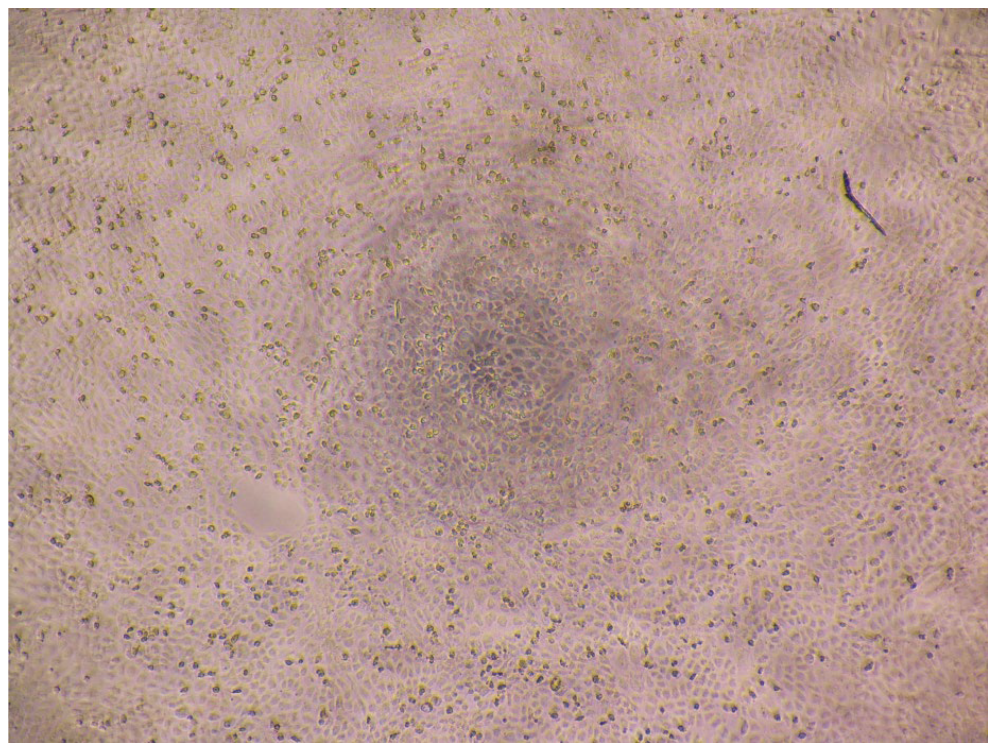 | Creatinine | 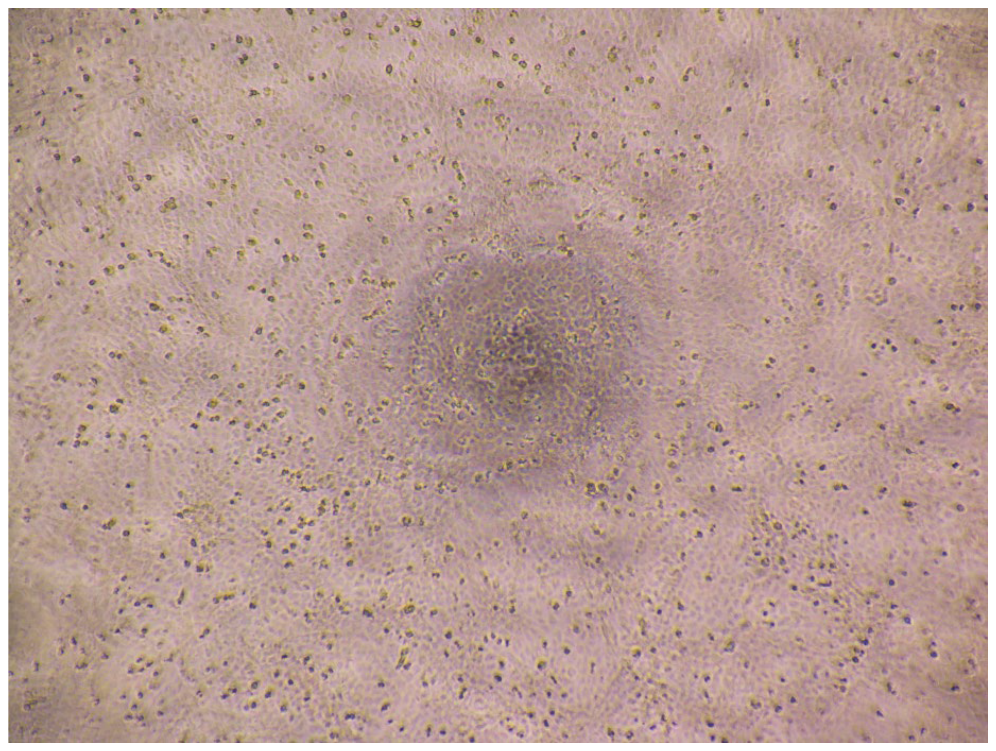 |
| Lopinavir | 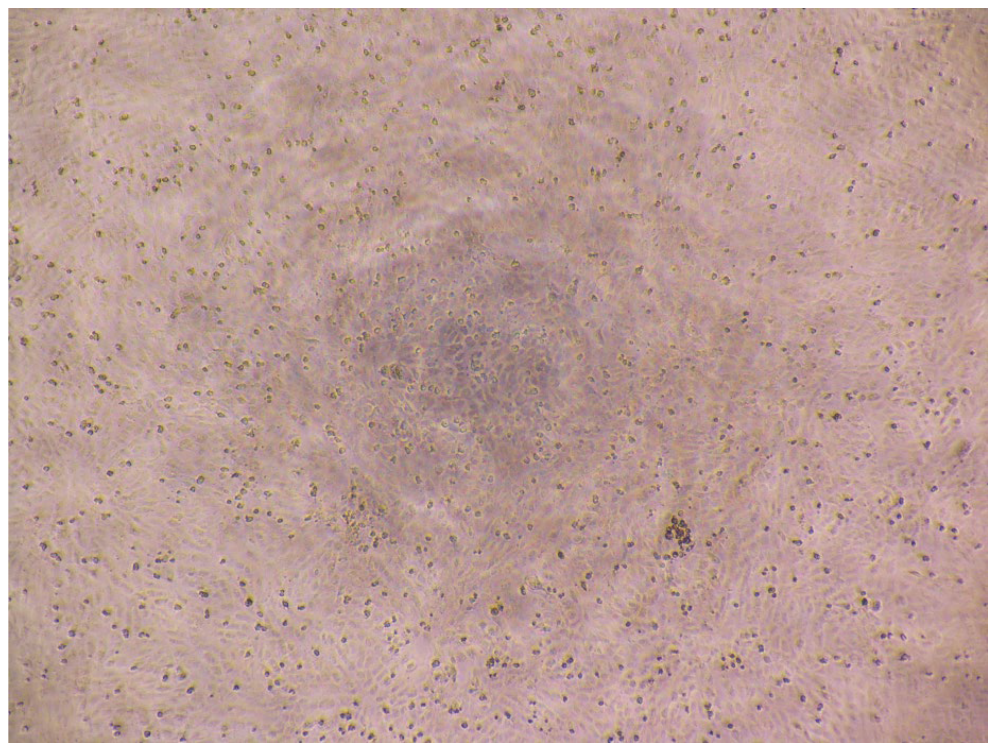 | Benztropine Mesylate | 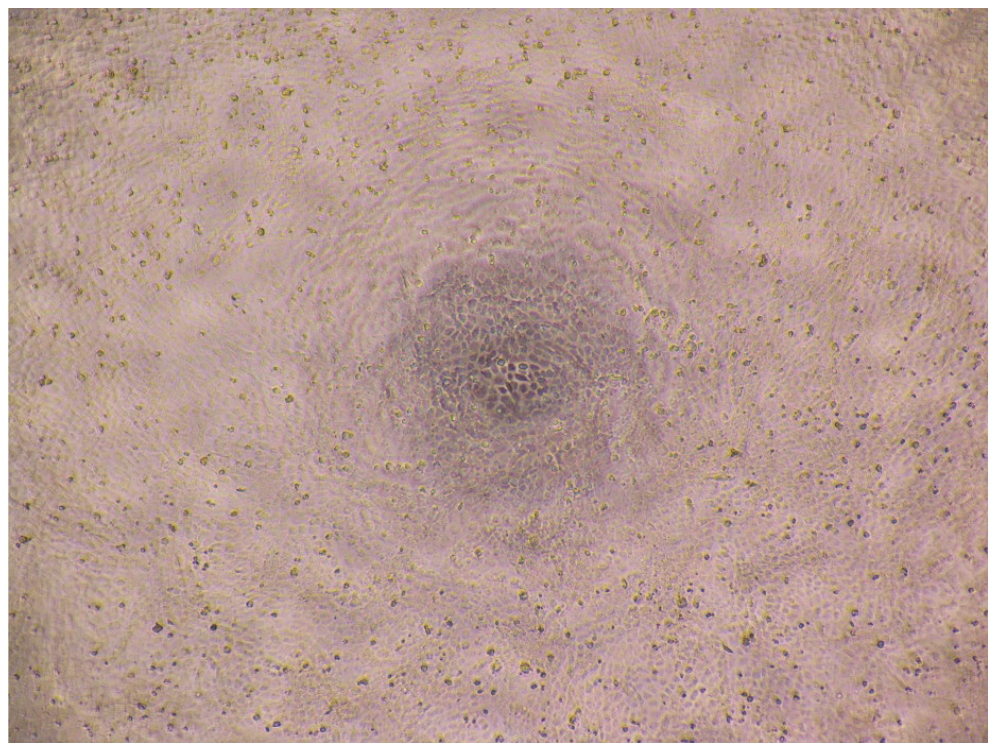 |
| Ranolazine Dihydrochloride | 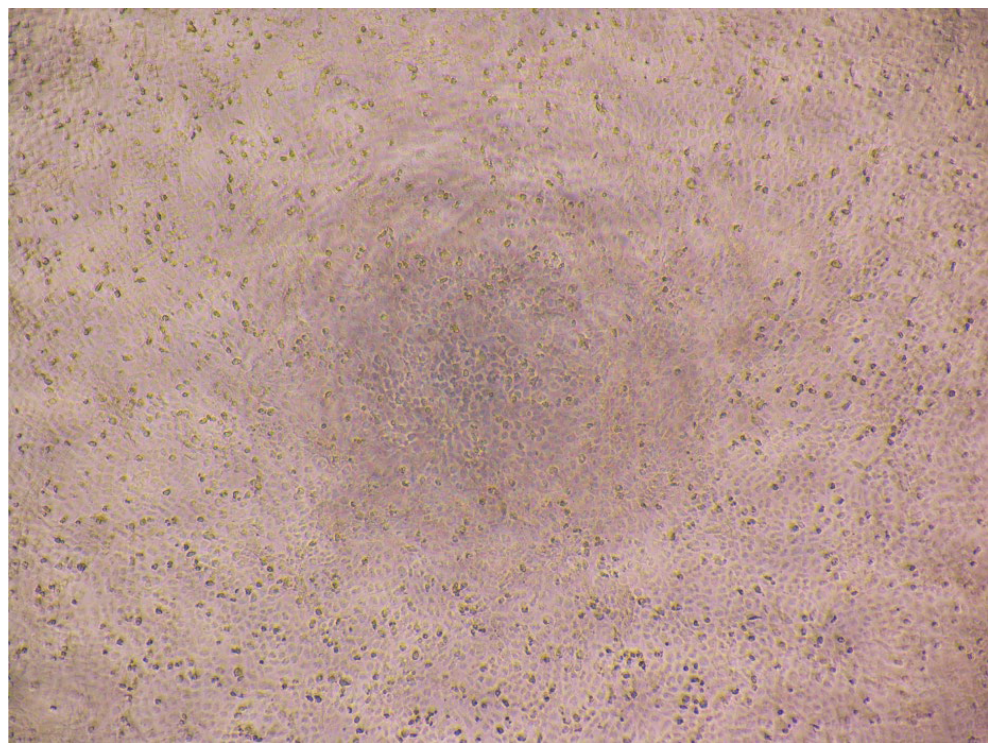 | Clinafloxacin | 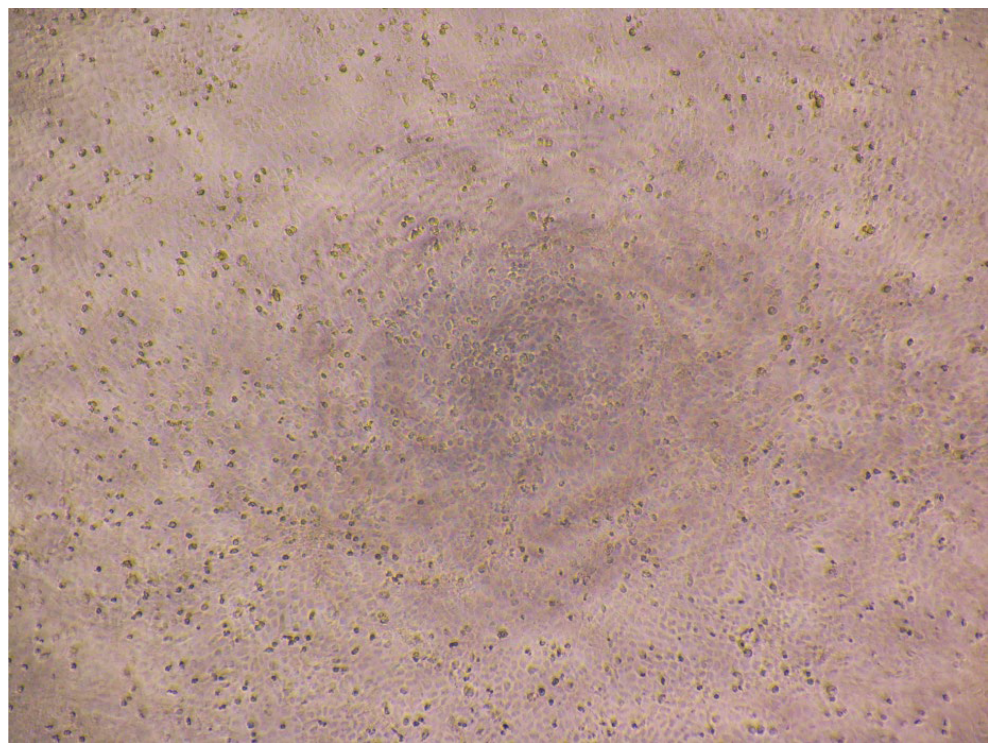 |
| Tranilast | 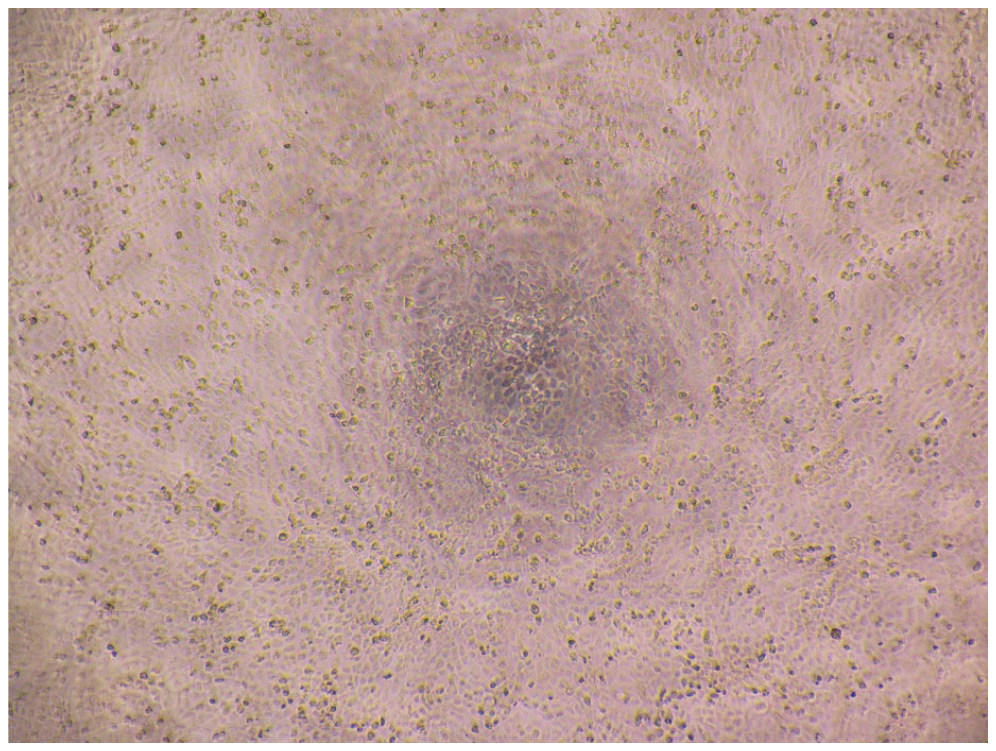 | Altrenogest | 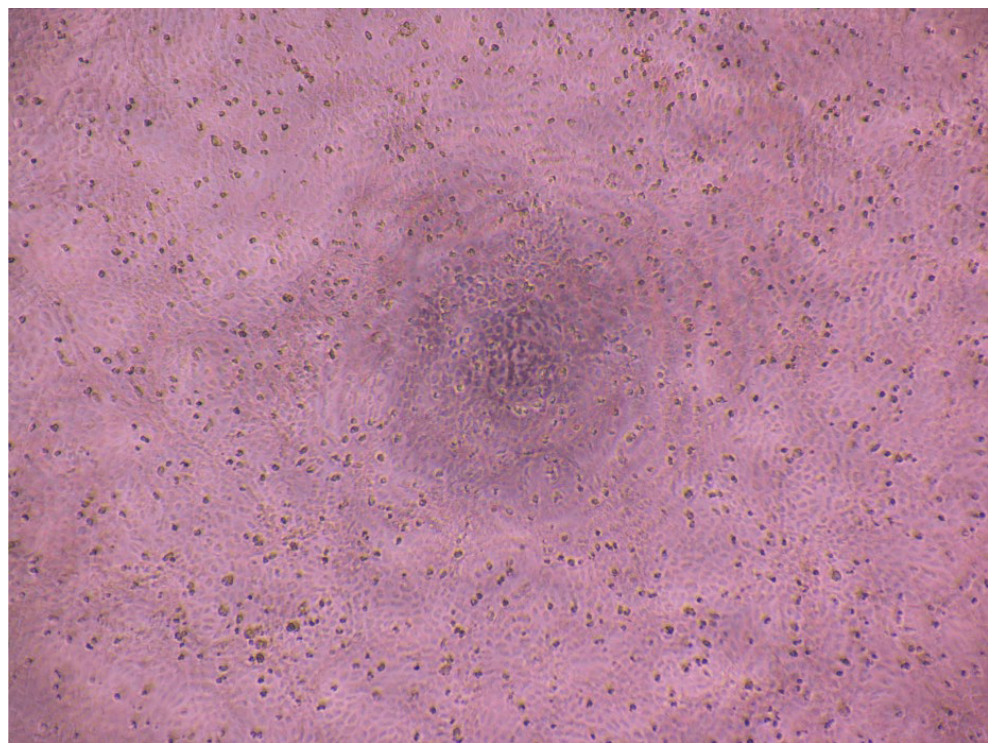 |
| Ofloxacin (Floxin) | 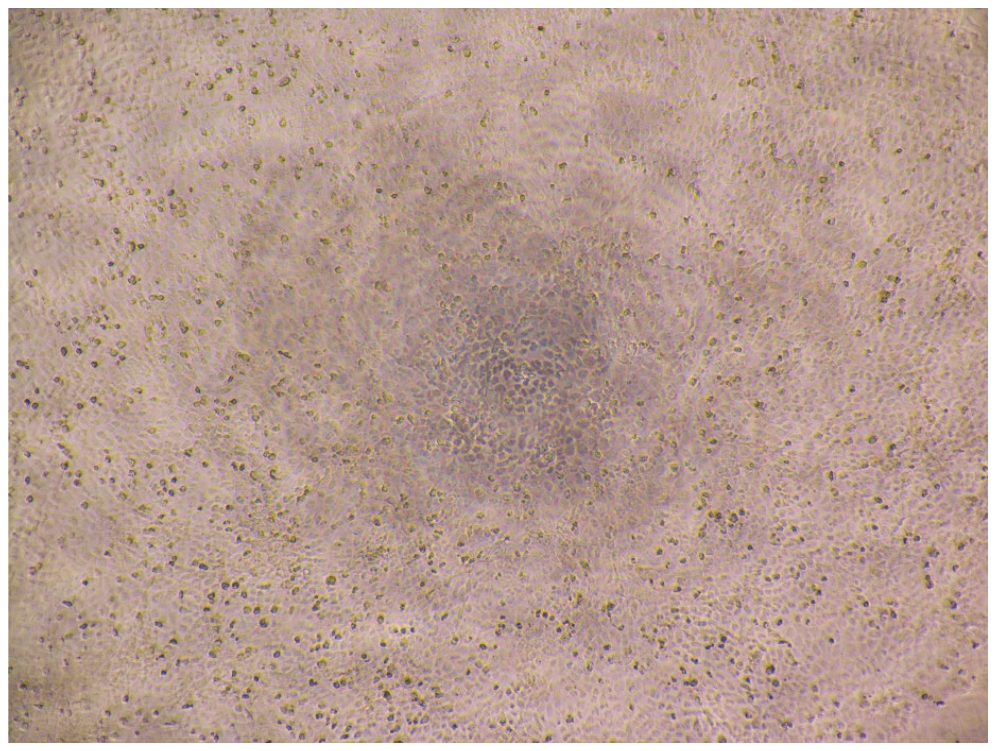 | Avanafil | 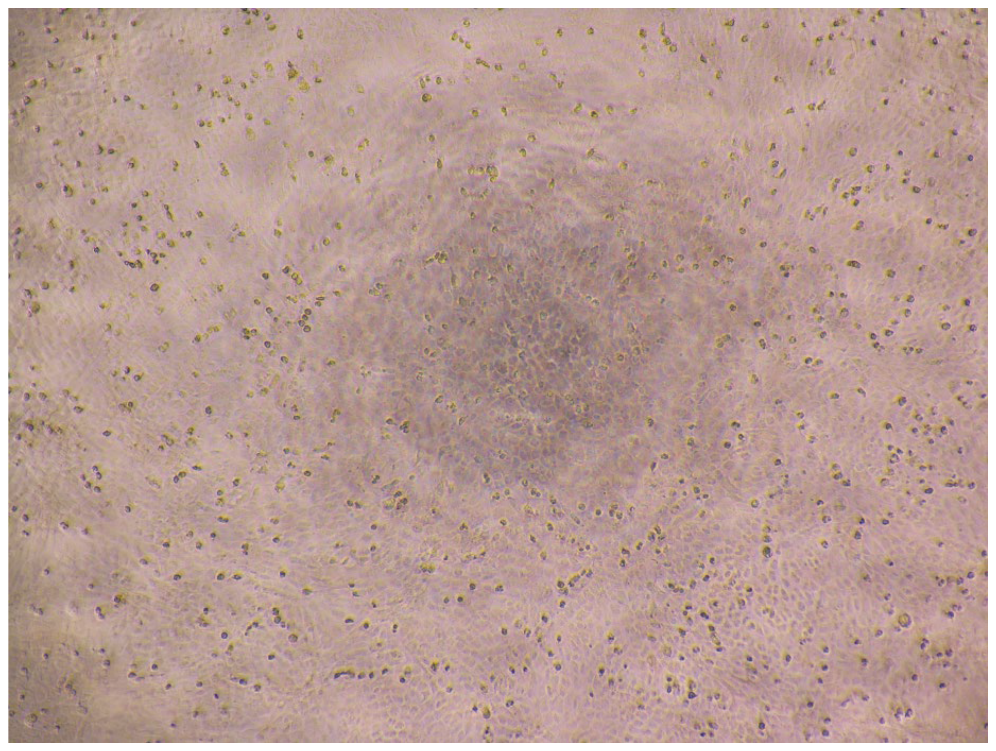 |
| Erythromycin | 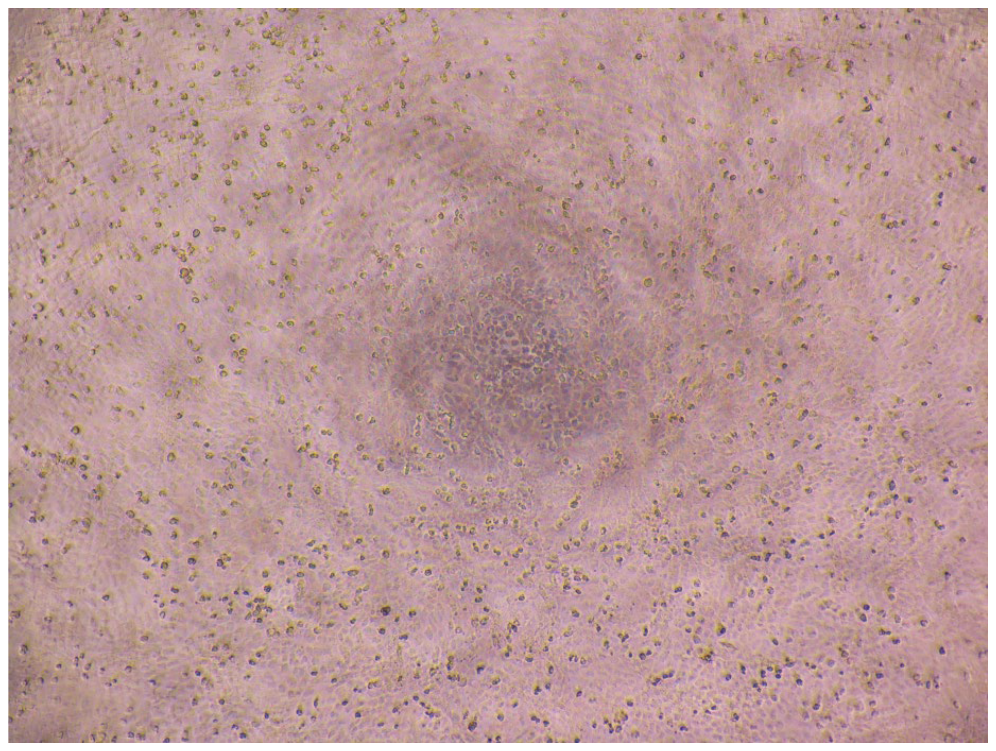 | Trifluoperazine Dihydrochloride | 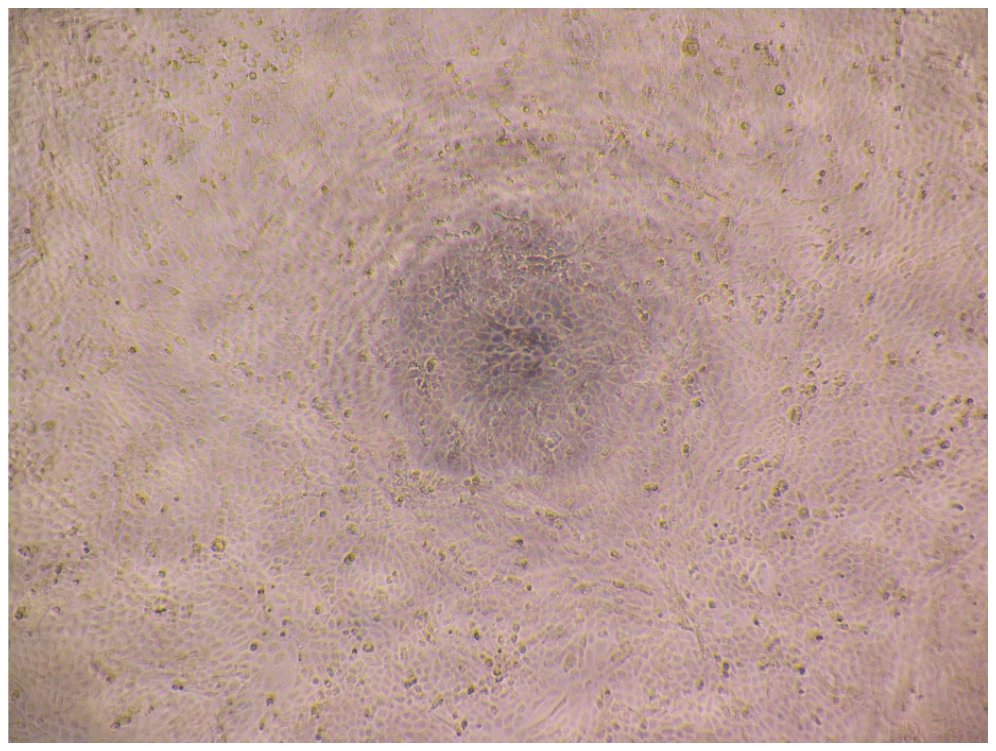 |
| Moxifloxacin Hydrochloride | 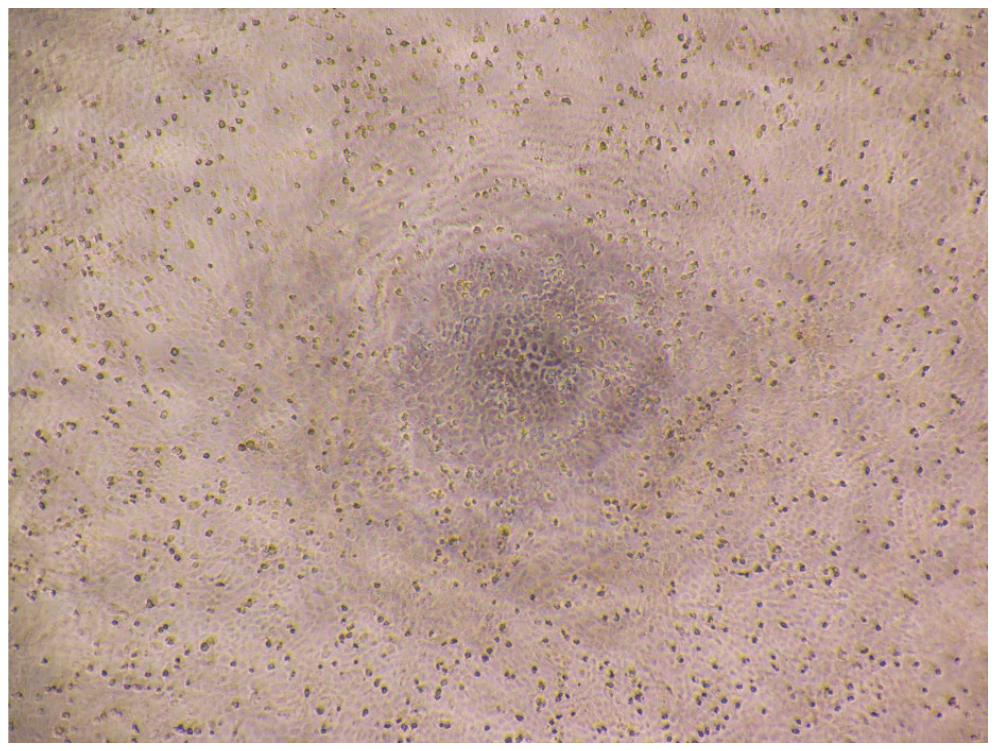 | Sertraline Hydrochloride | 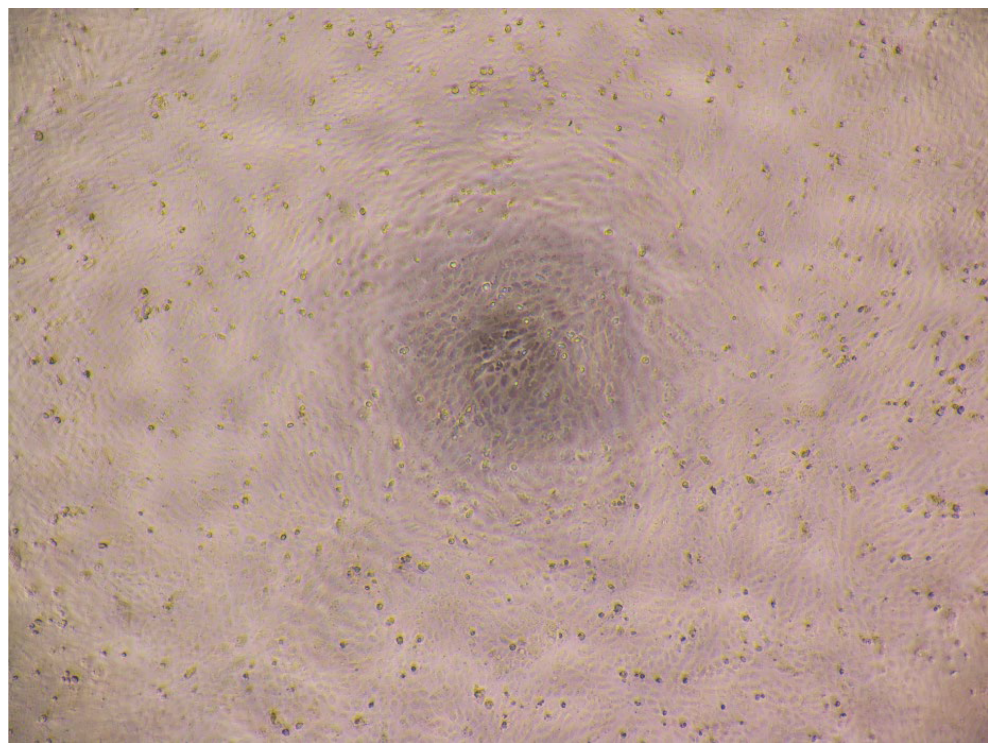 |
| Calcitriol | 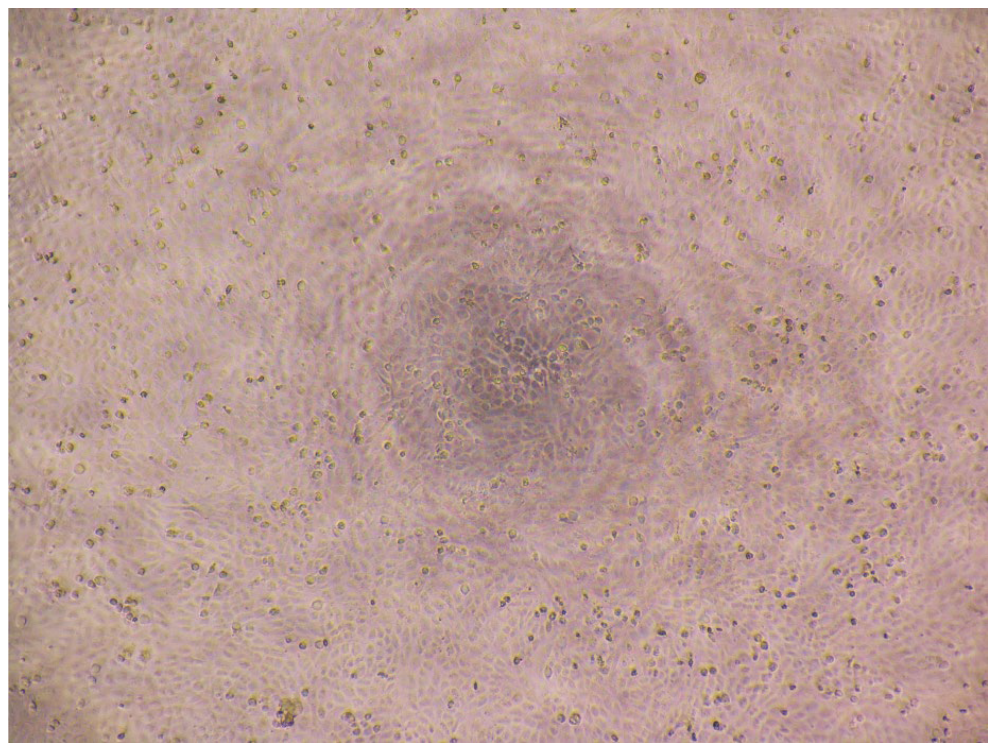 | Dexlansoprazole | 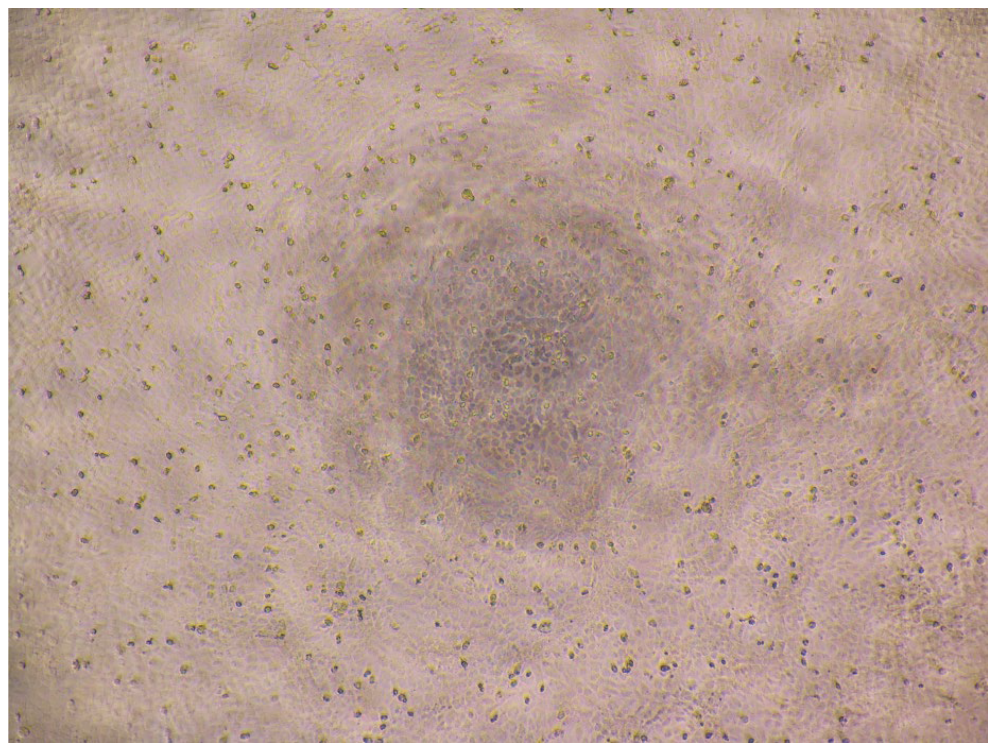 |
| Telaprevir | 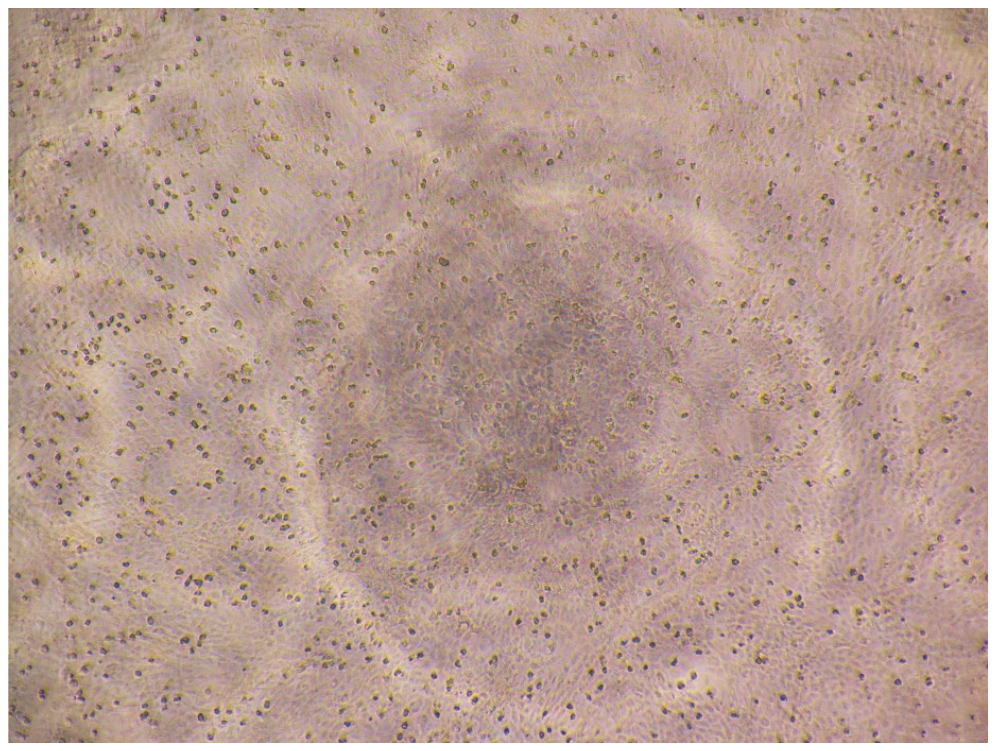 | Closantel Sodium | 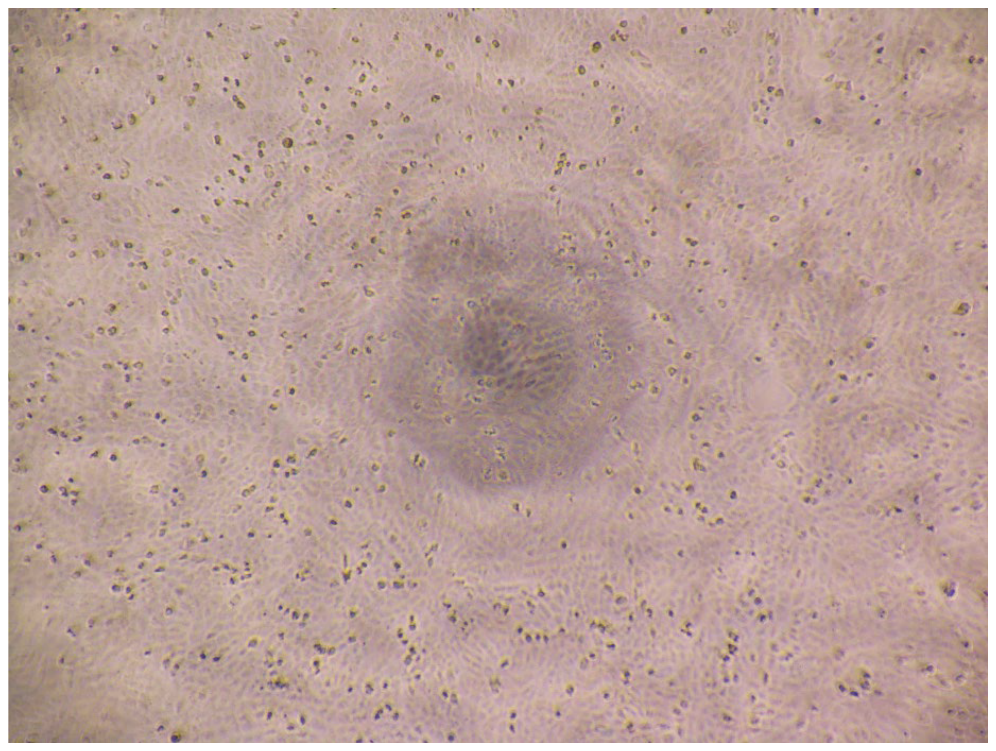 |
| Calcifediol | 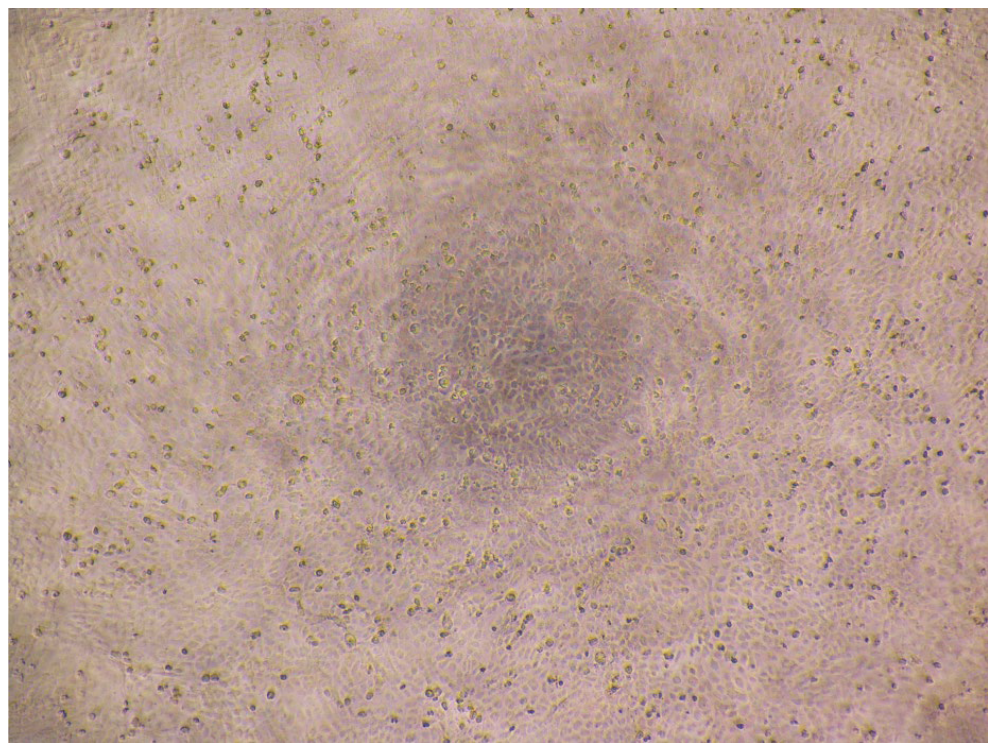 | Famprofazone | 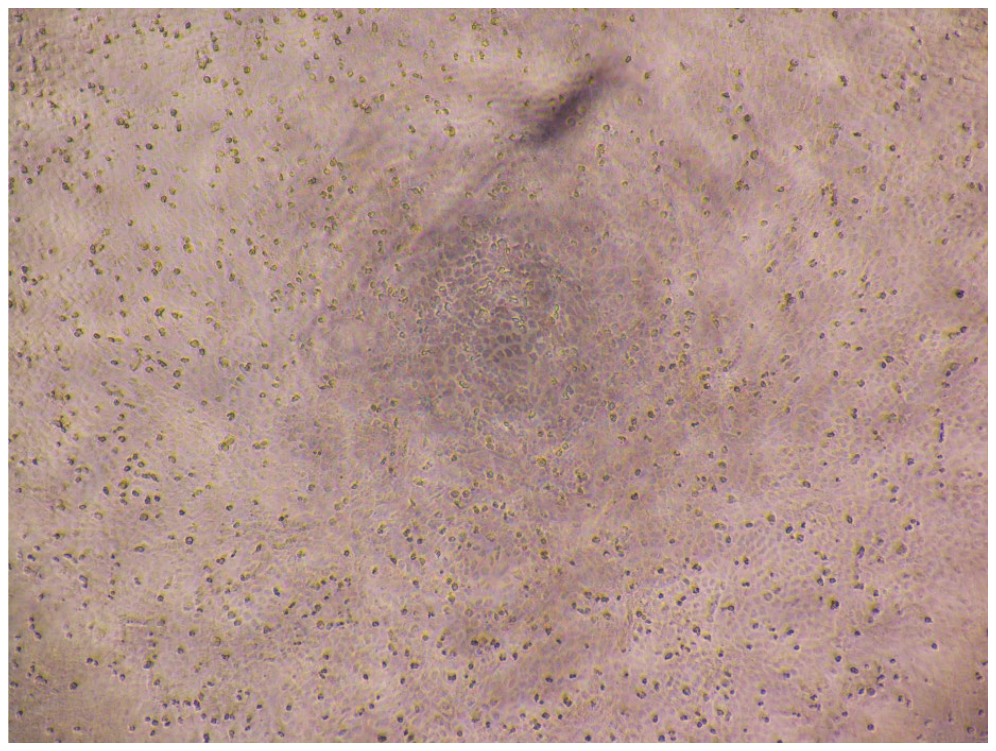 |
| Cyclosporine | 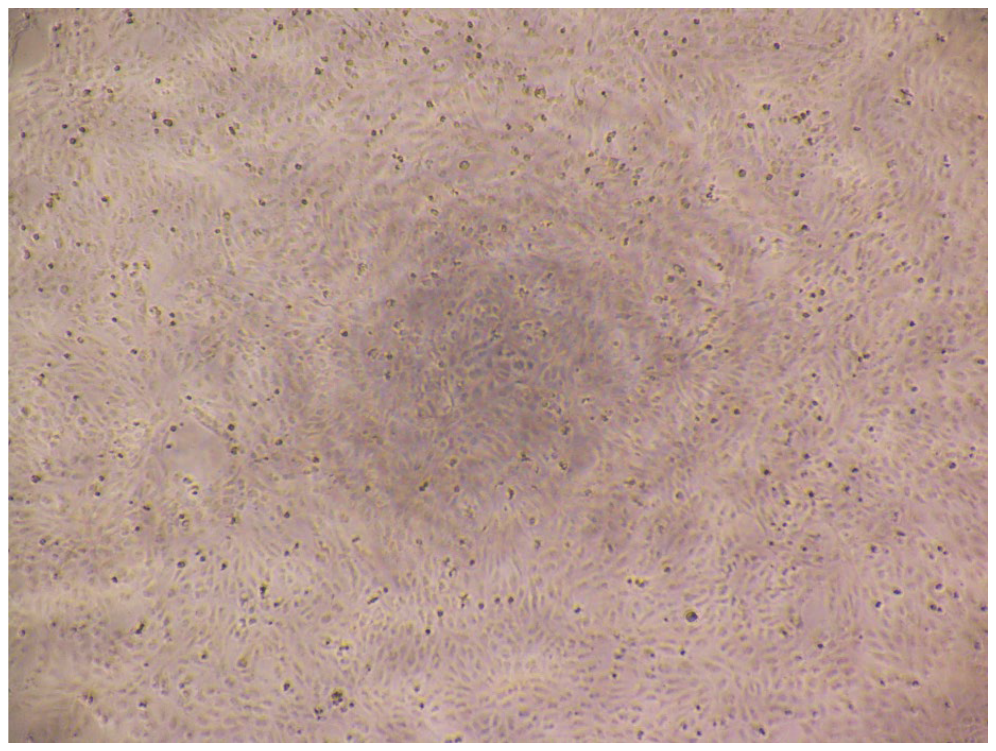 | Metaproterenol Sulfate (Orciprenaline) | 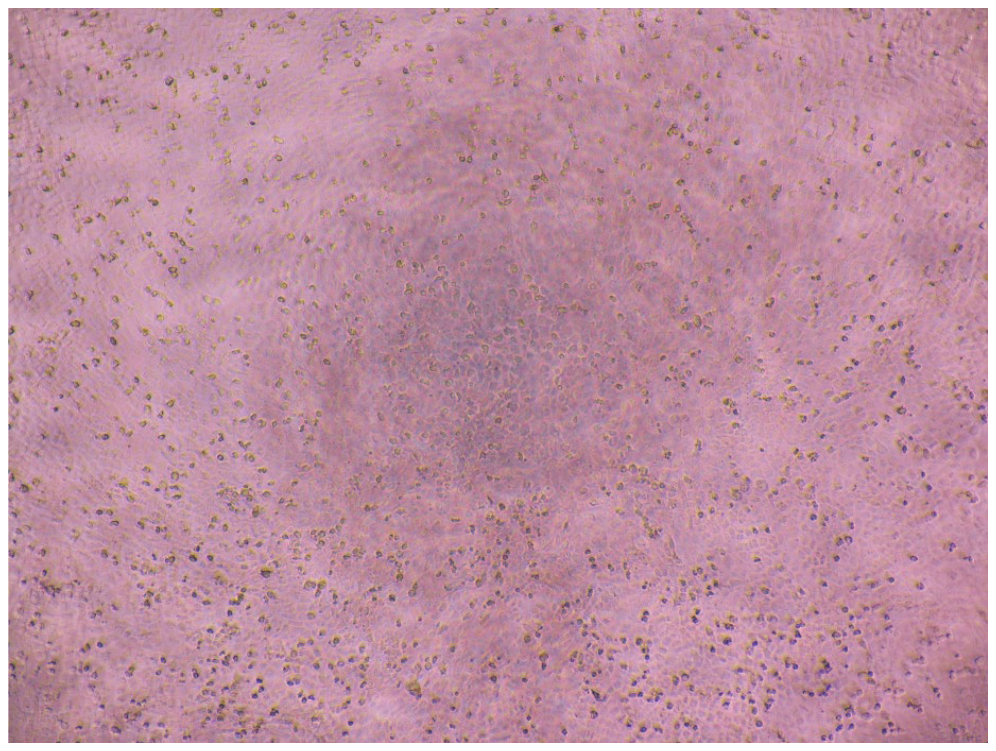 |
| Telbivudine | 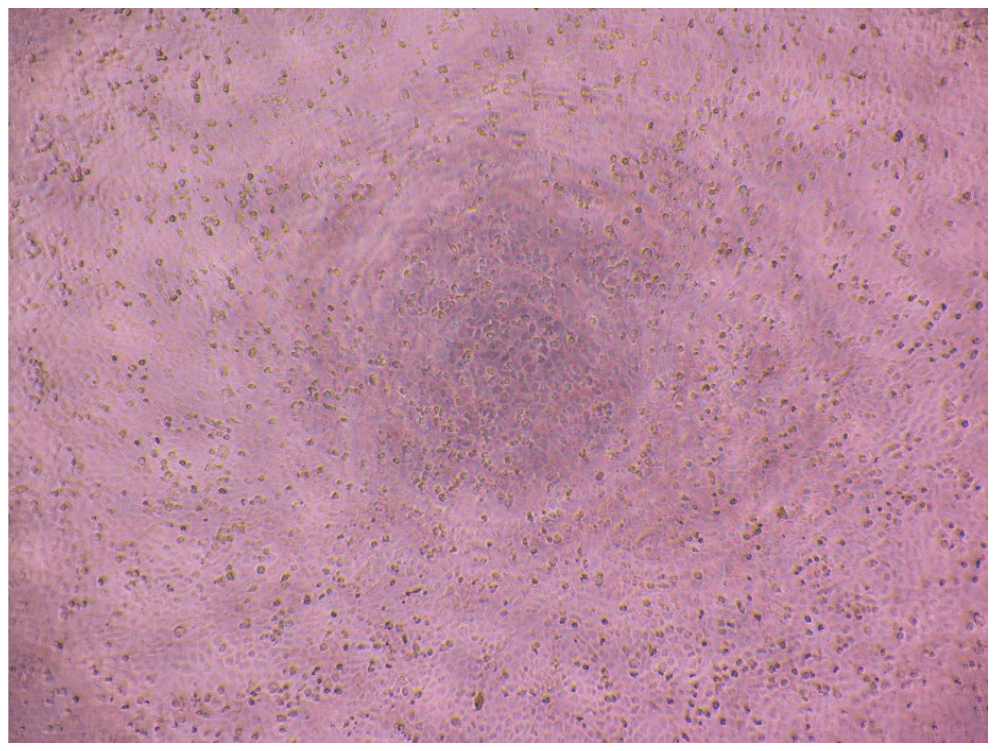 | Ractopamine Hydrochloride | 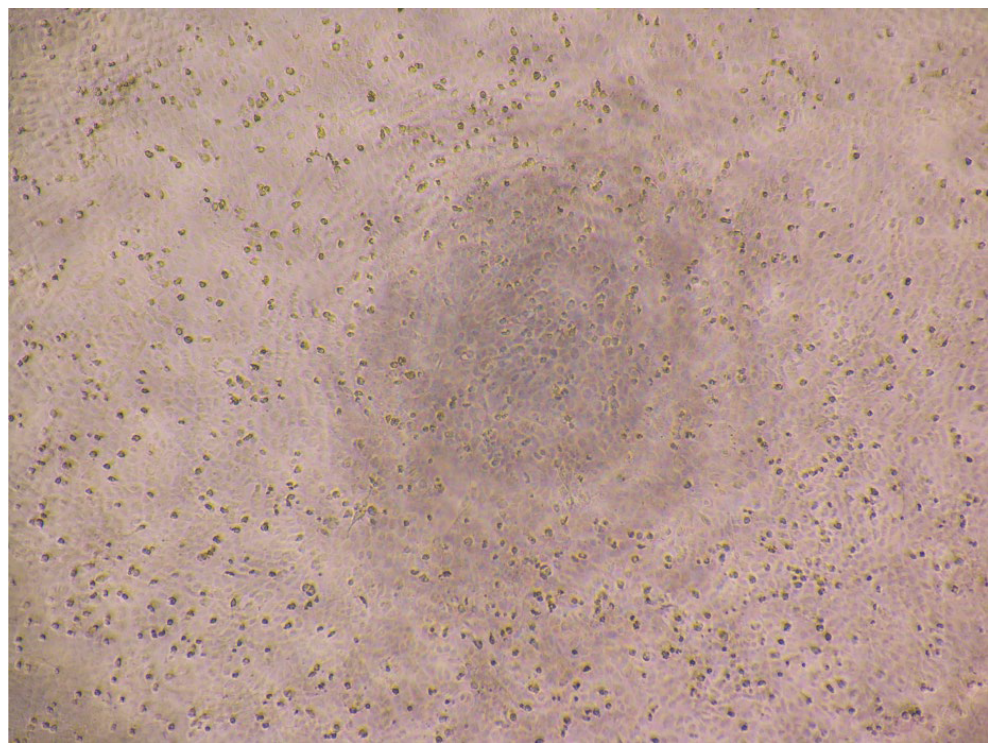 |
| Nevirapine | 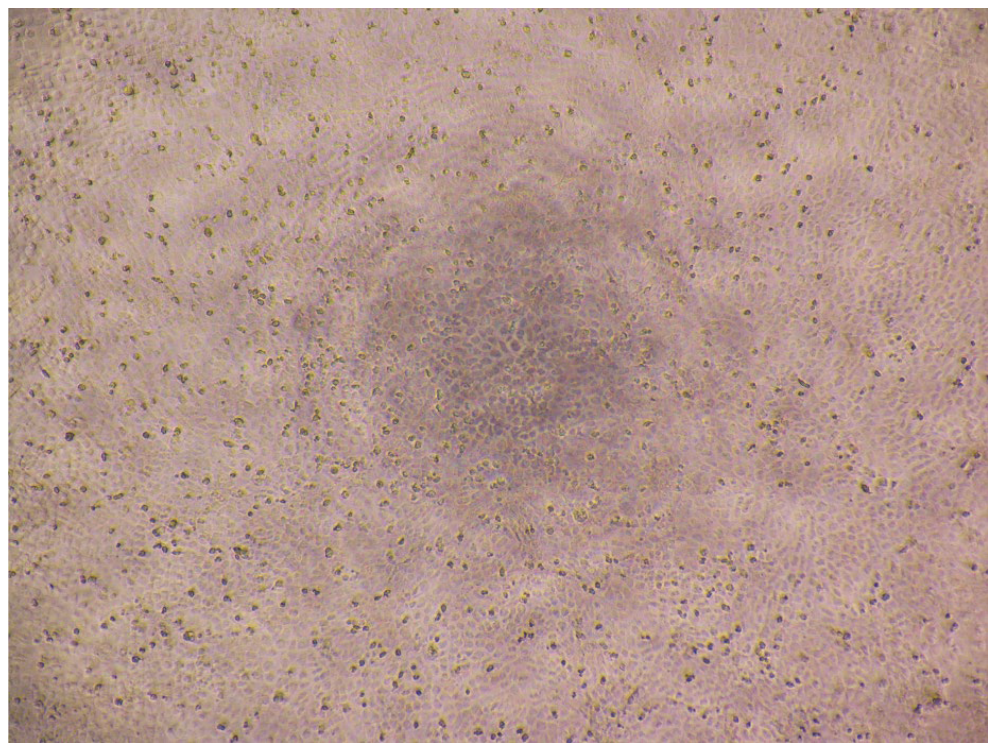 | Terfenadine | 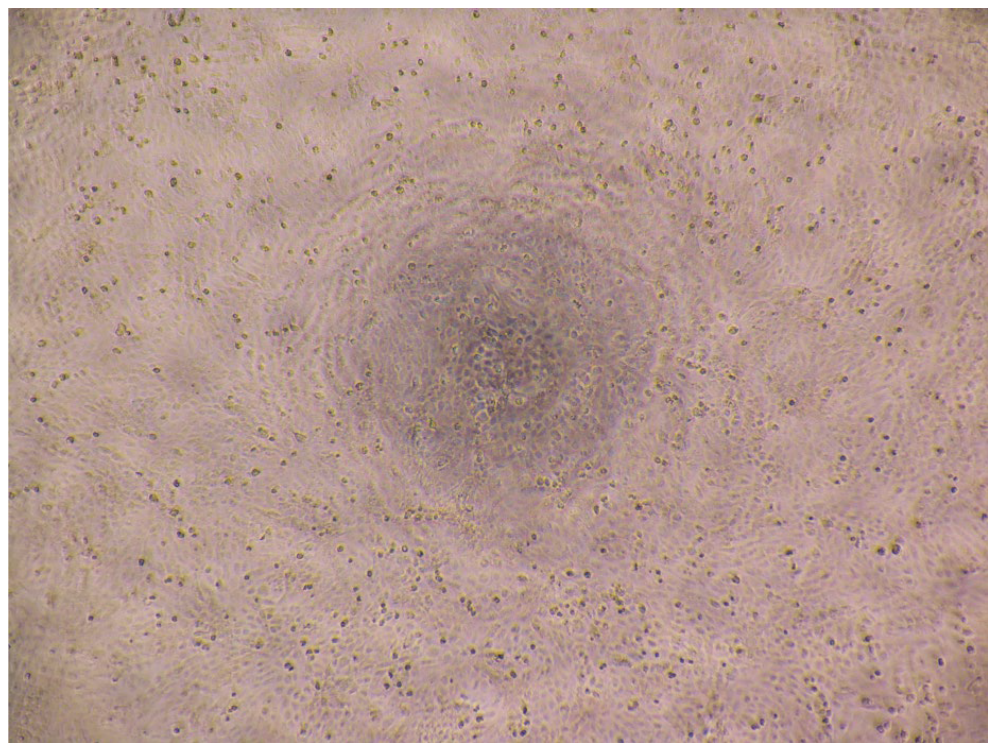 |
| Esomeprazole | 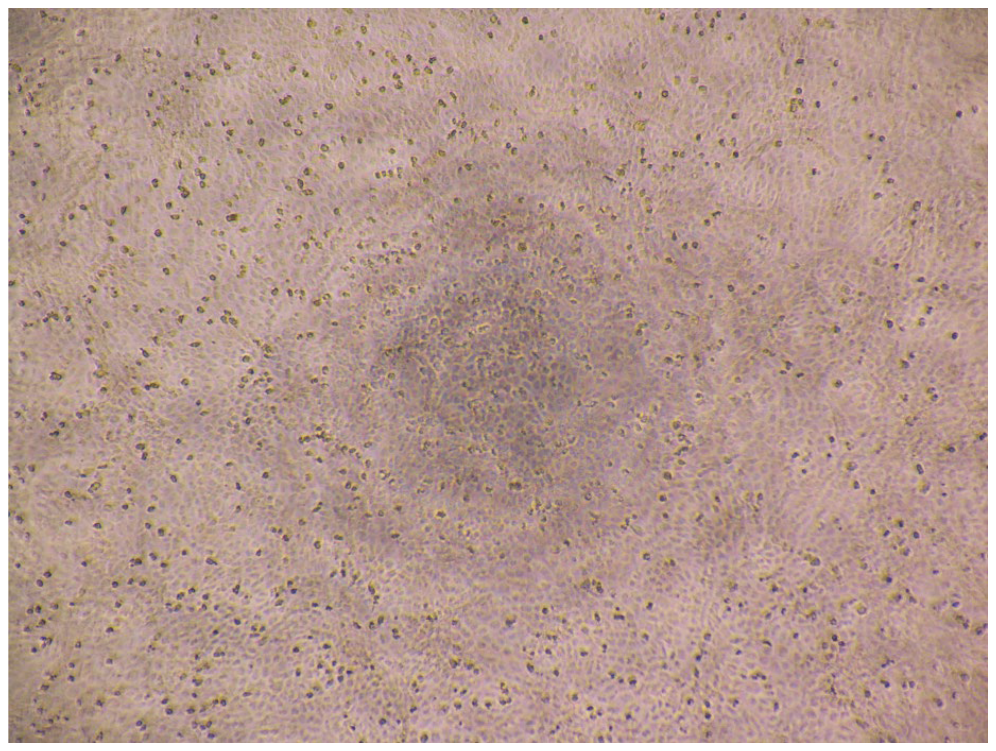 | Thiostrepton | 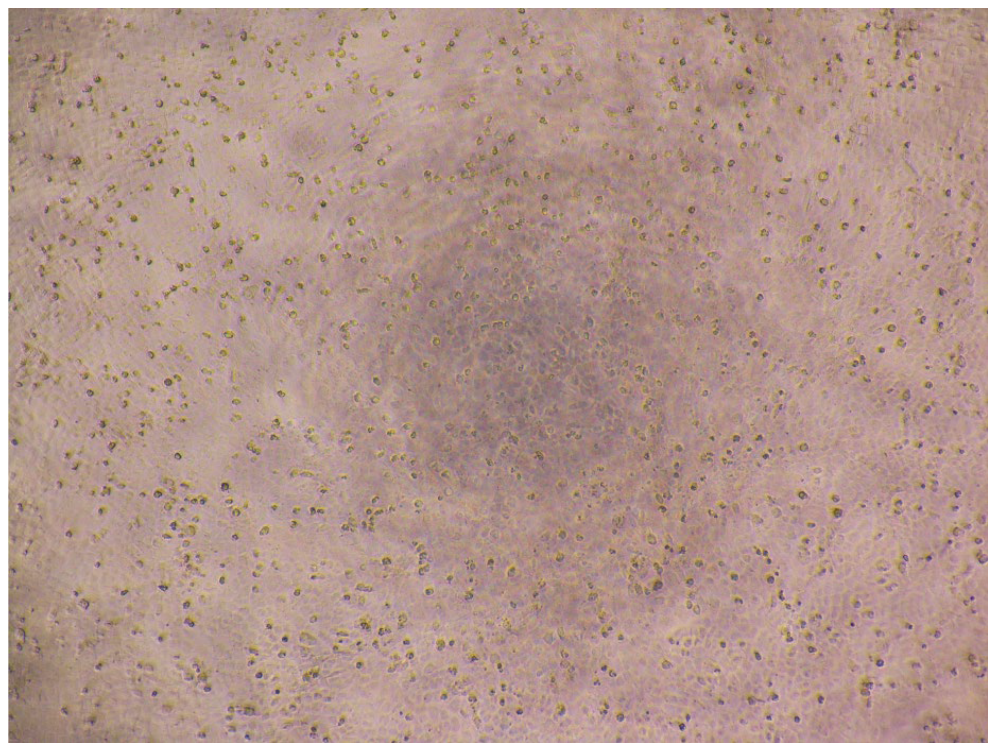 |
| Progesterone | 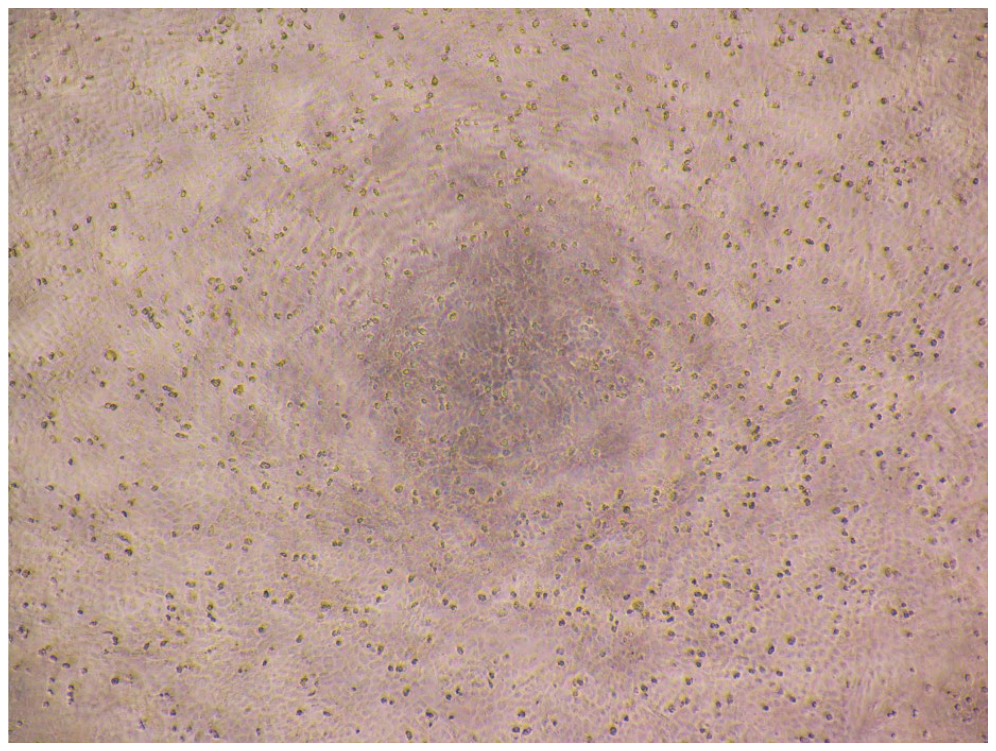 | Glafenine Hydrochloride | 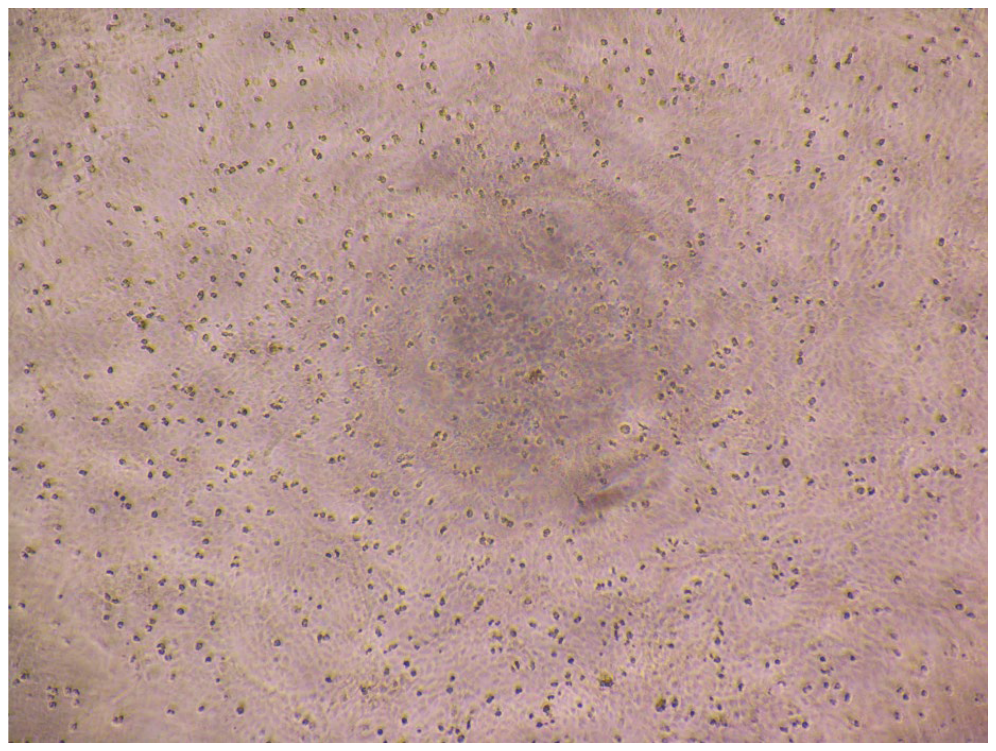 |
| Nimodipine | 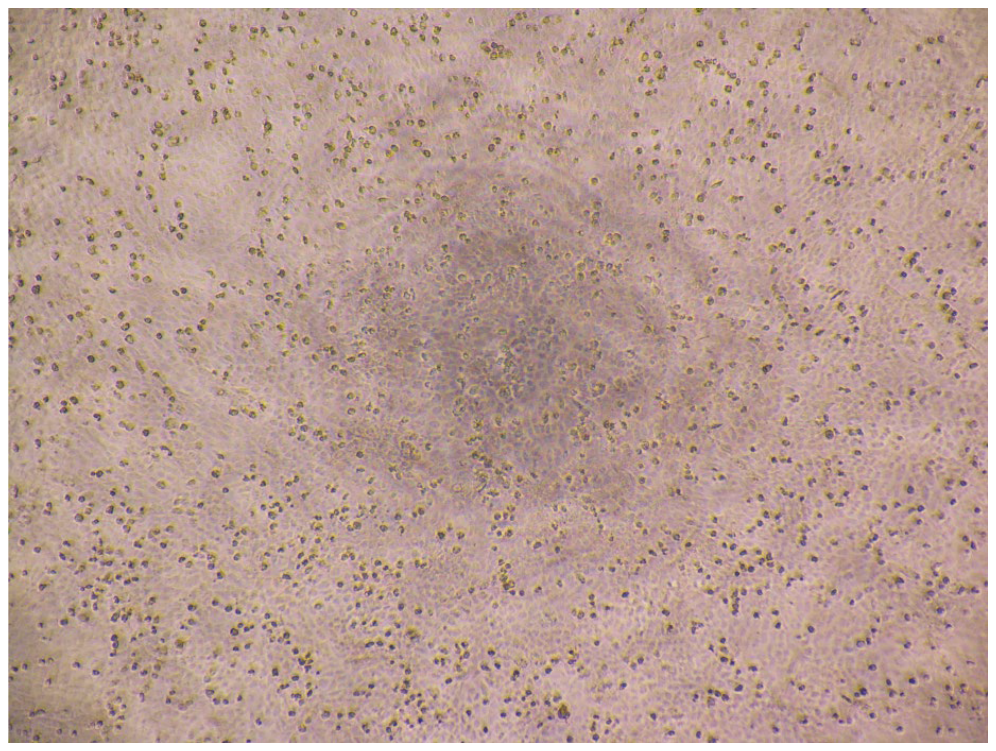 | Meclocycline Sulfosalicylate | 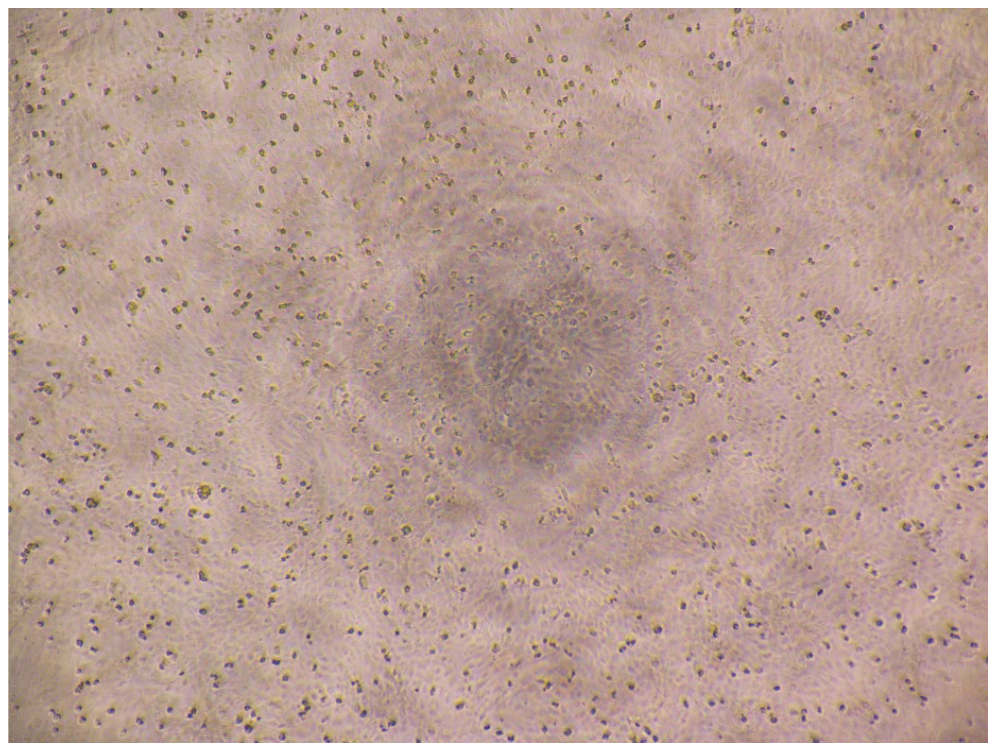 |

**Table S2**: List of primary hits identified from the primary screen performed using an ACE2 inhibitors library.

| **Compound/Treatment** | **Image** | **Compound/Treatment** | **Image** |
| --- | --- | --- | --- |
| Uninfected | 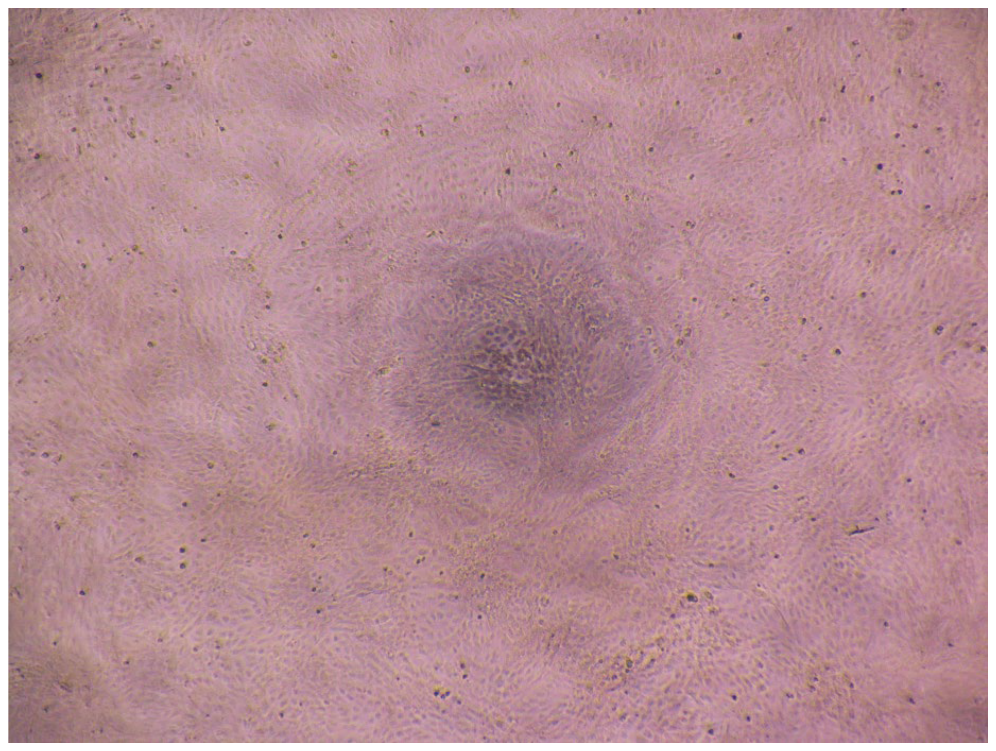 | 0.1% DMSO | 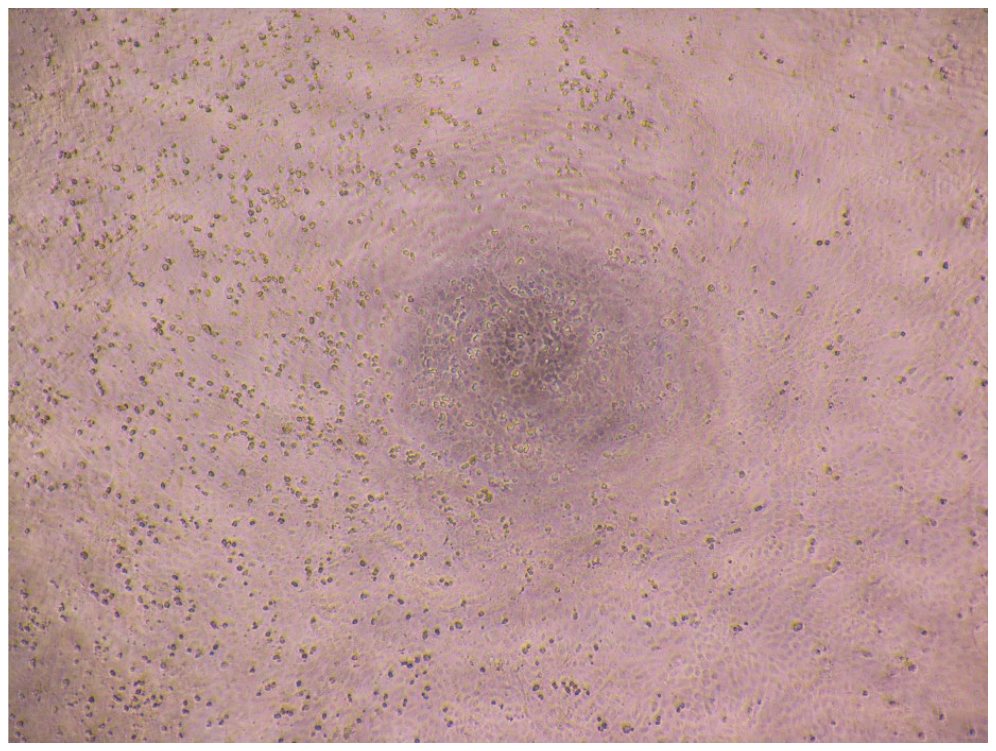 |
| Remdesivir | 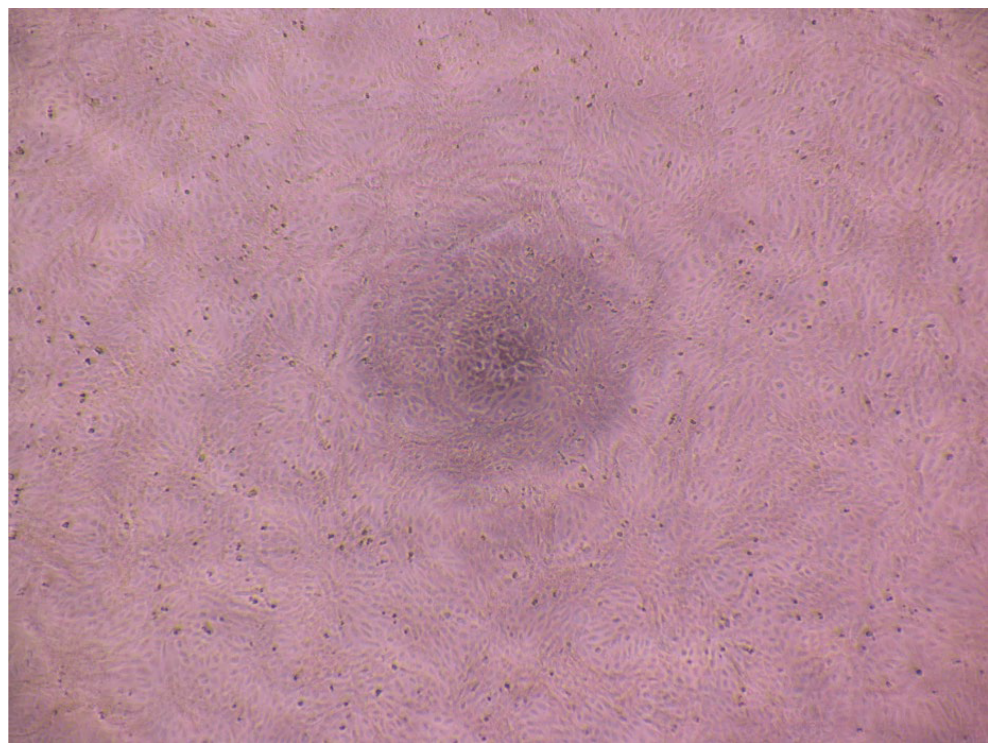 |  |  |
| Citicoline | 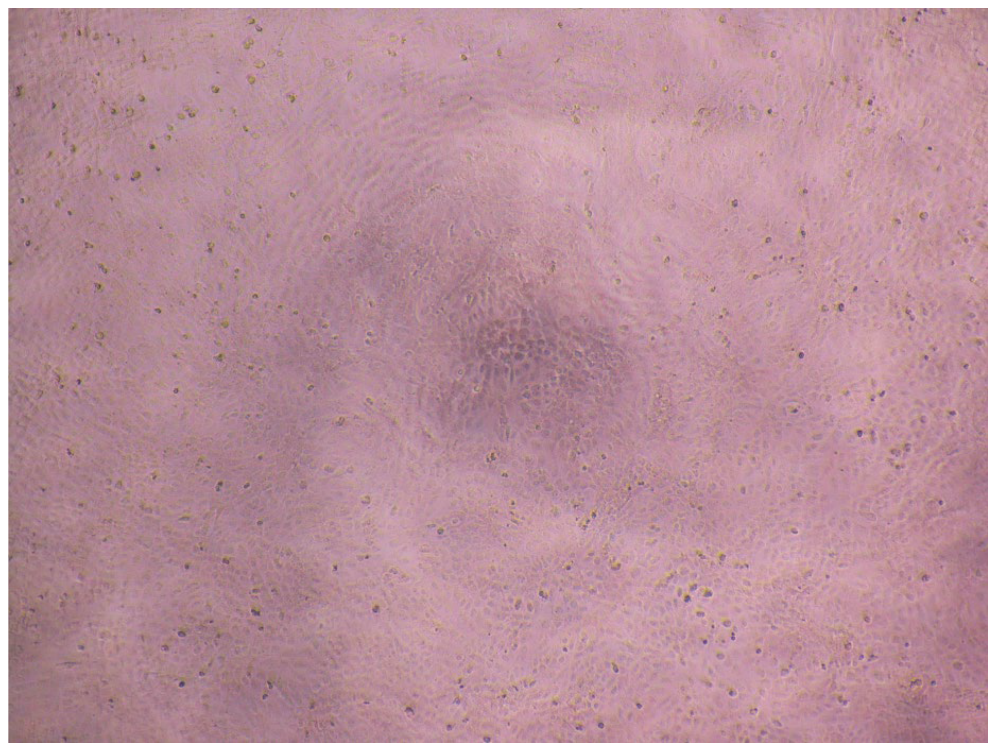 | Nuezhenide | 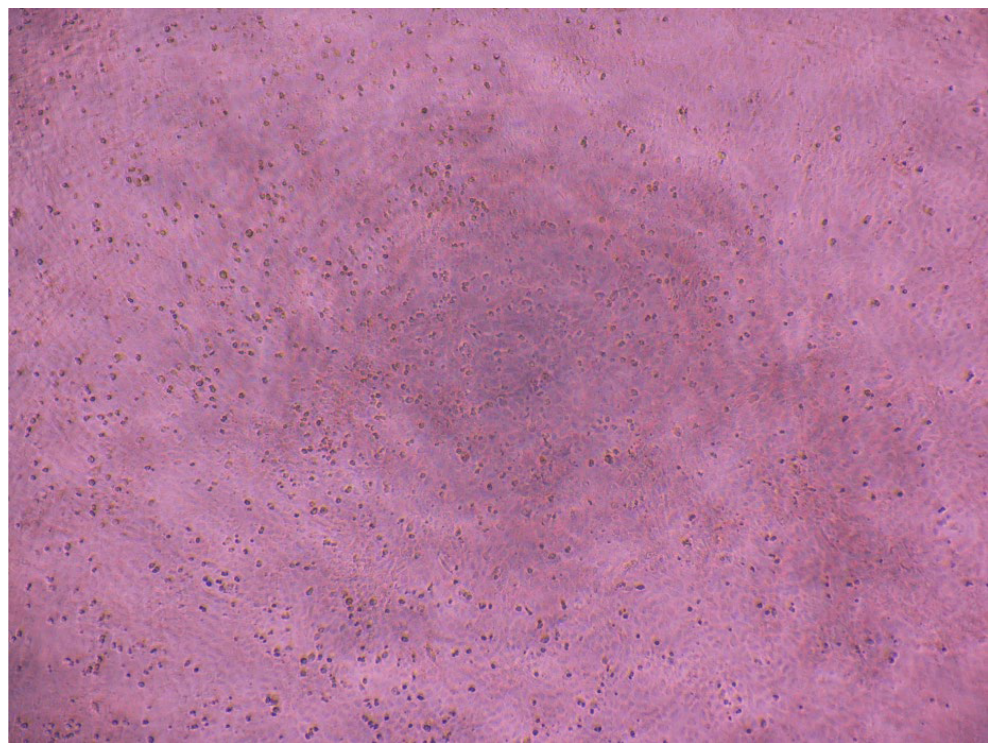 |
| Cefpiramide Acid | 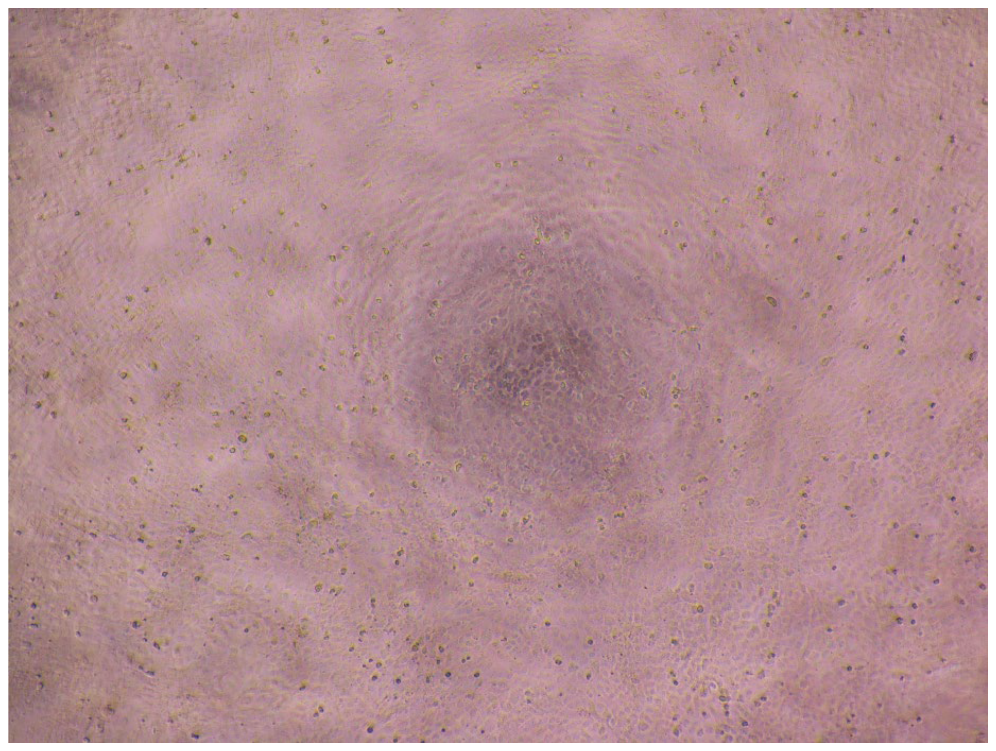 | Tepotinib | 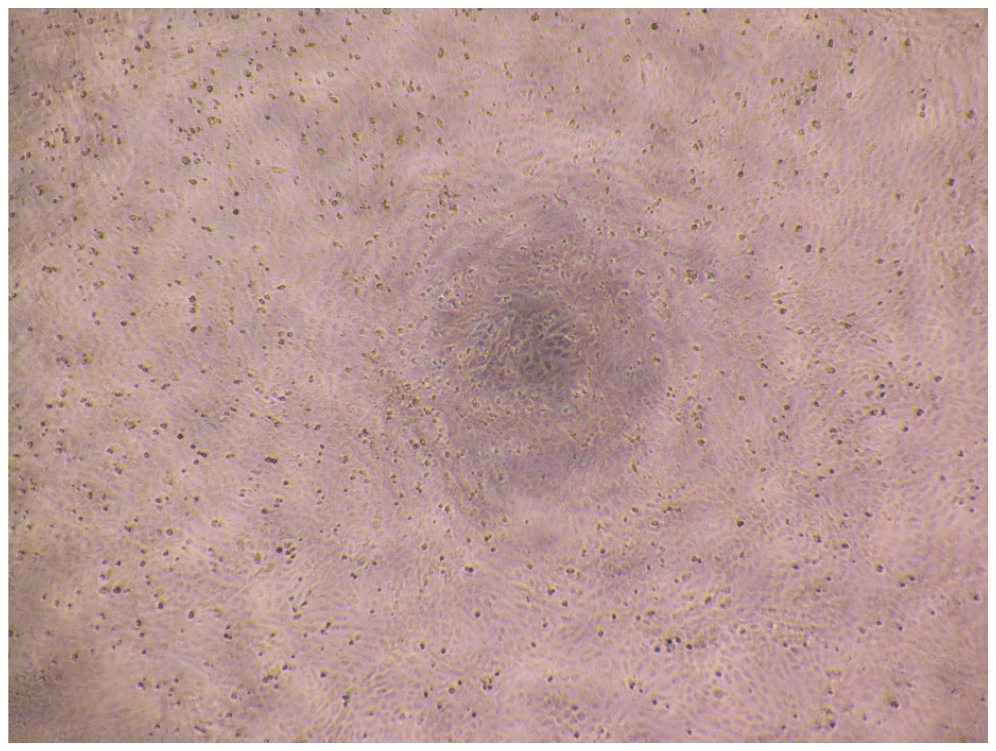 |
| Calcium Follinate | 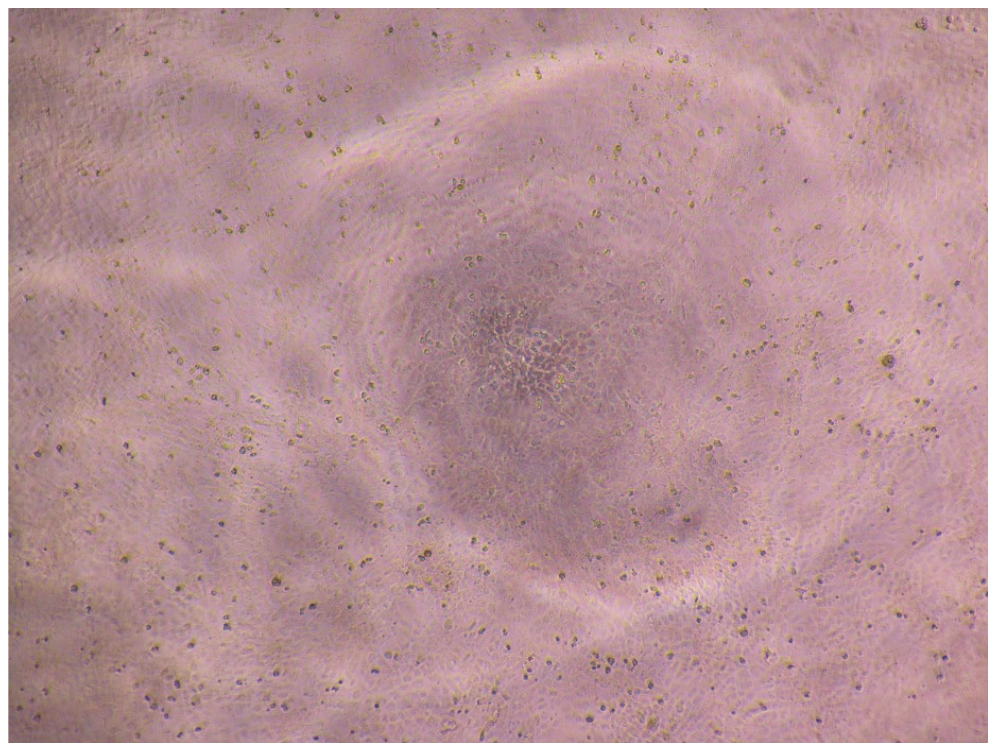 | Mulberroside A | 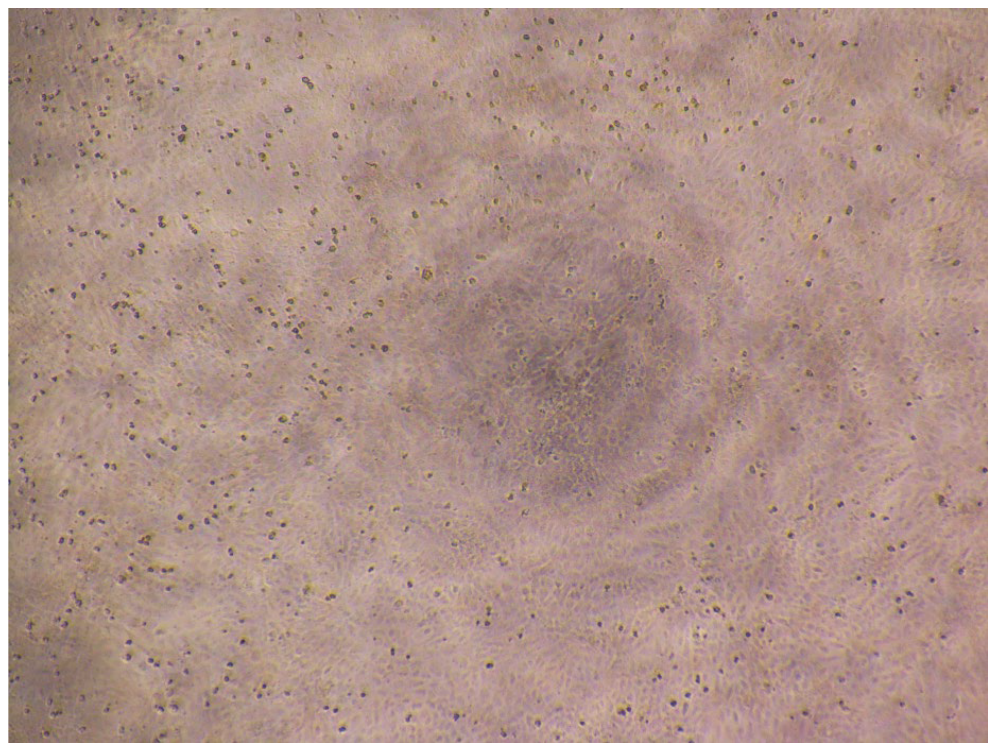 |
| Dronedarone Hydrochloride | 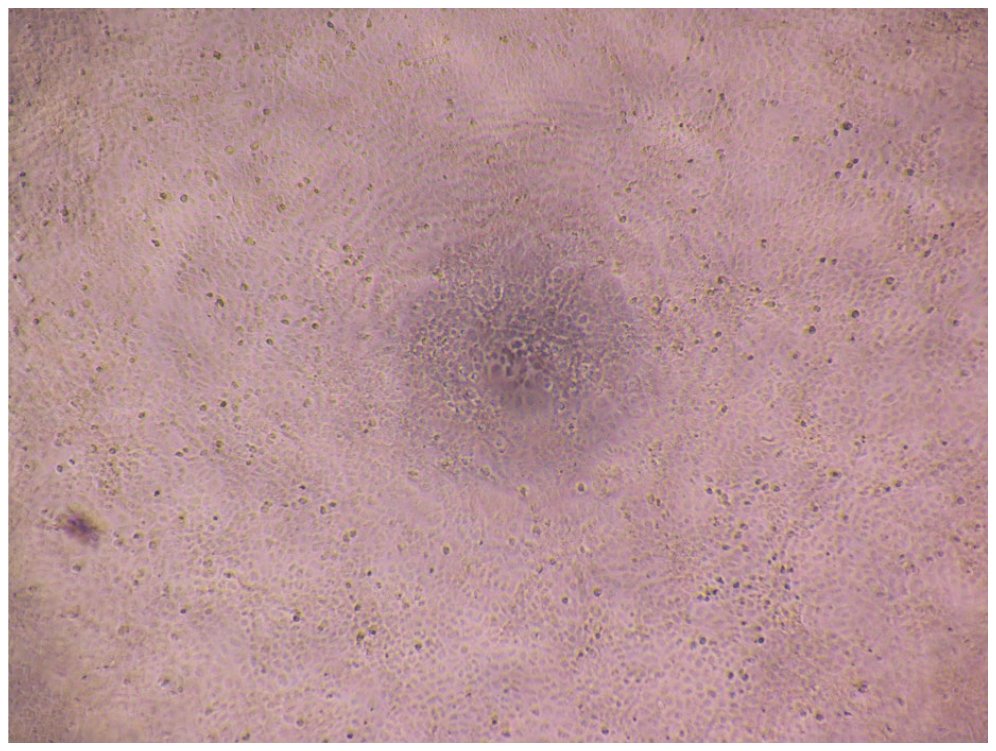 | 4,5-Dicaffeoylquinic acid | 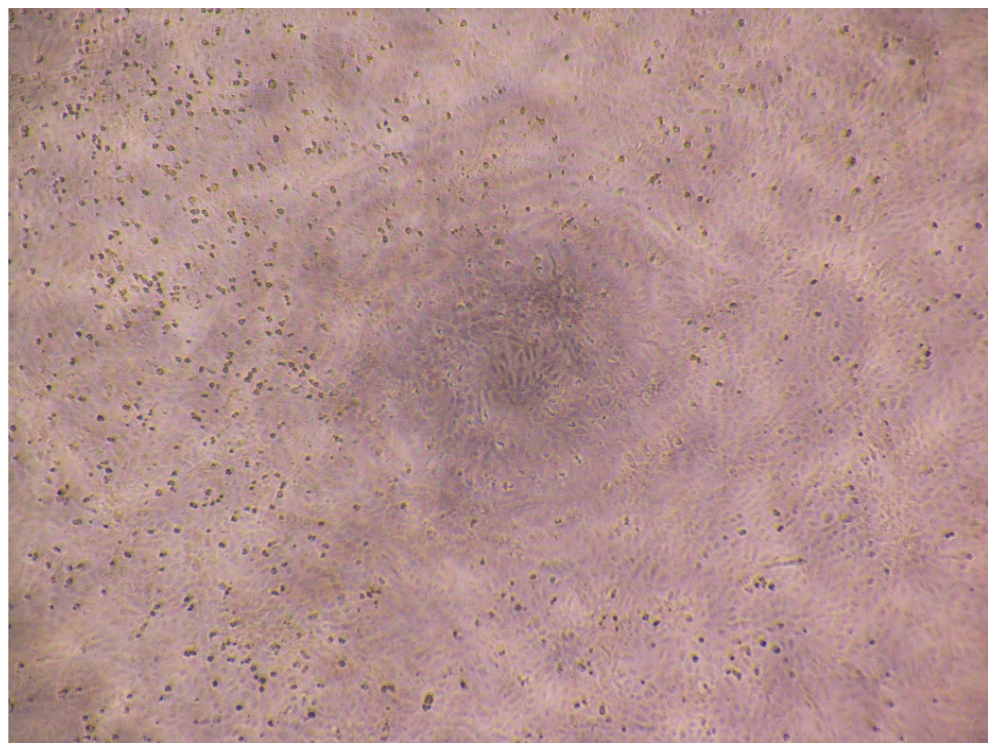 |
| Pravastatin Sodium | 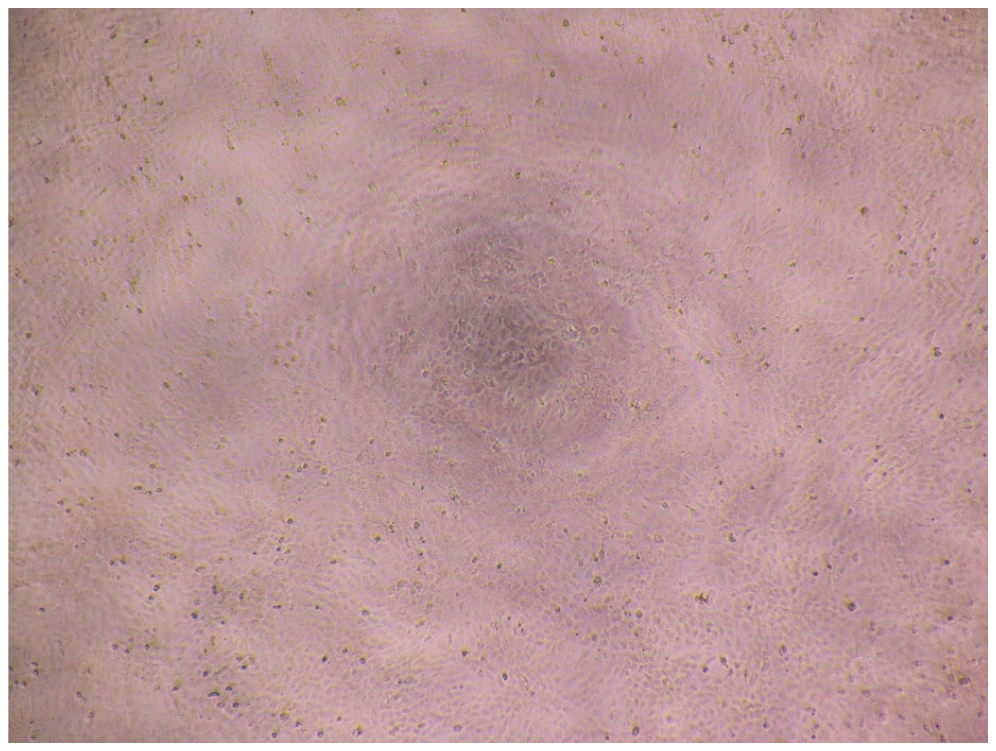 | Crocin I | 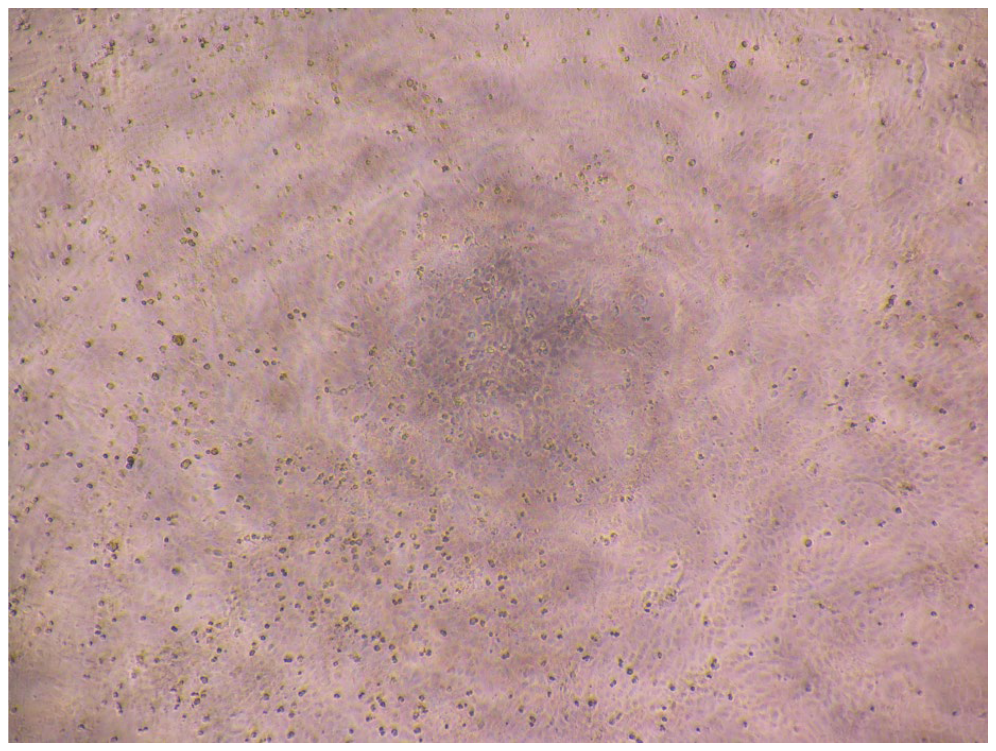 |
| Tenofovir Alafenamide | 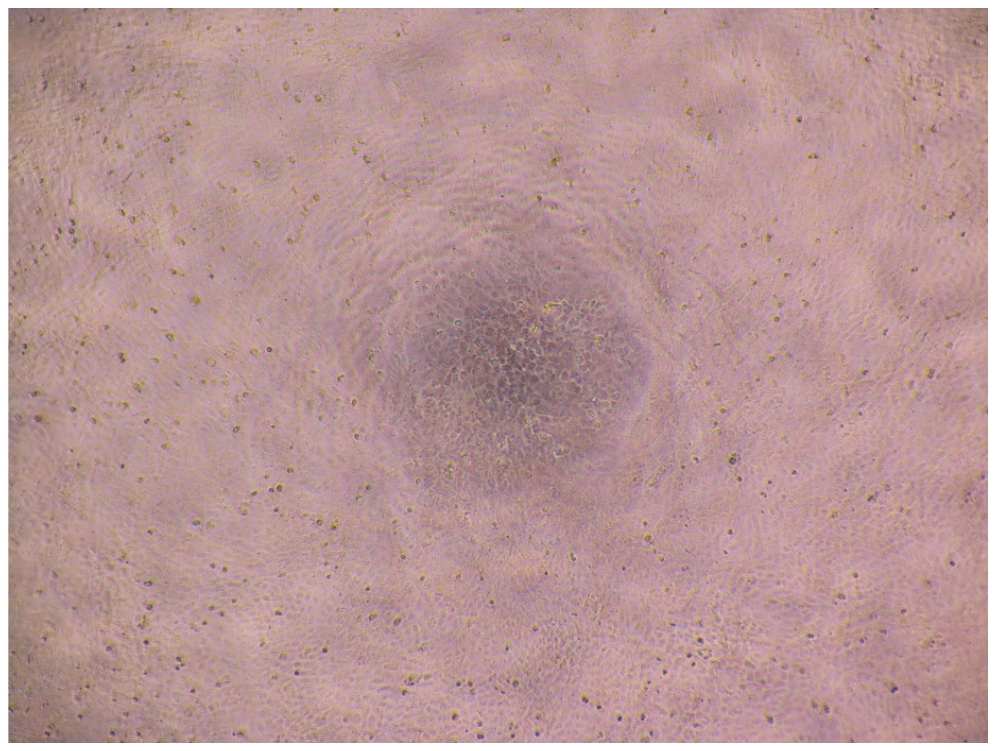 | PLX8394 | 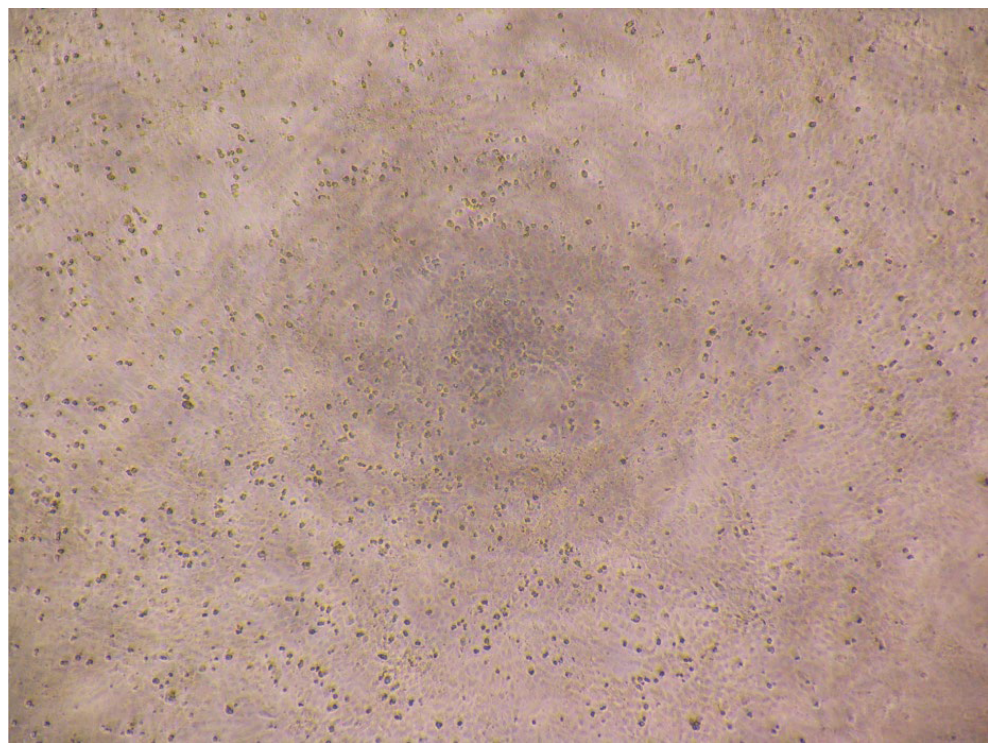 |
| Navitoclax | 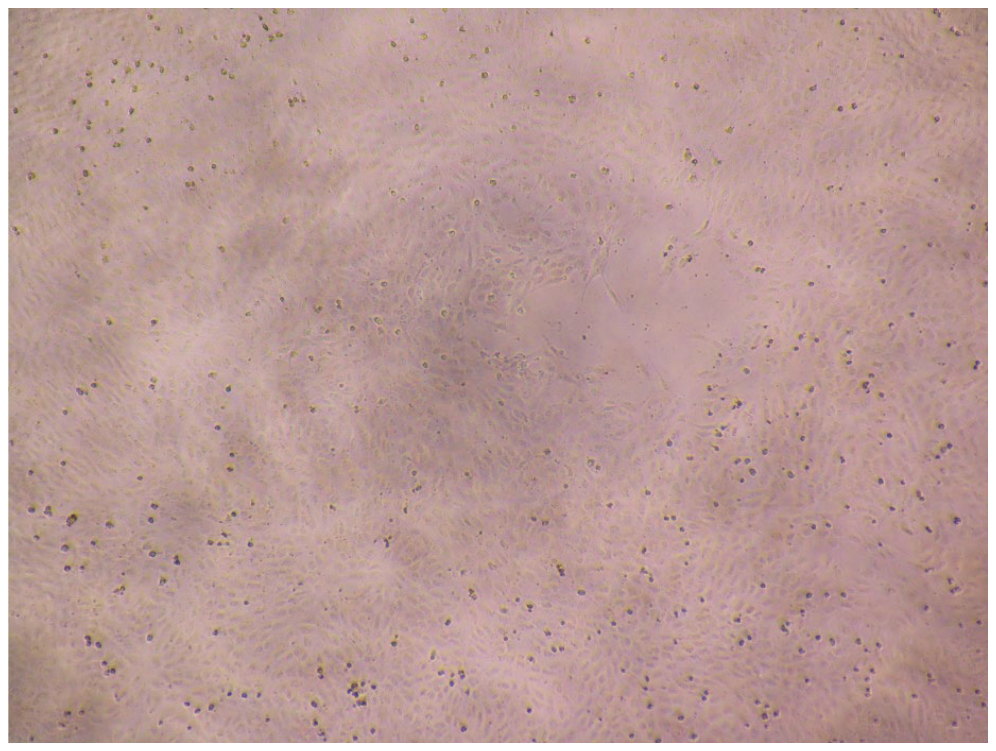 | Complanatoside | 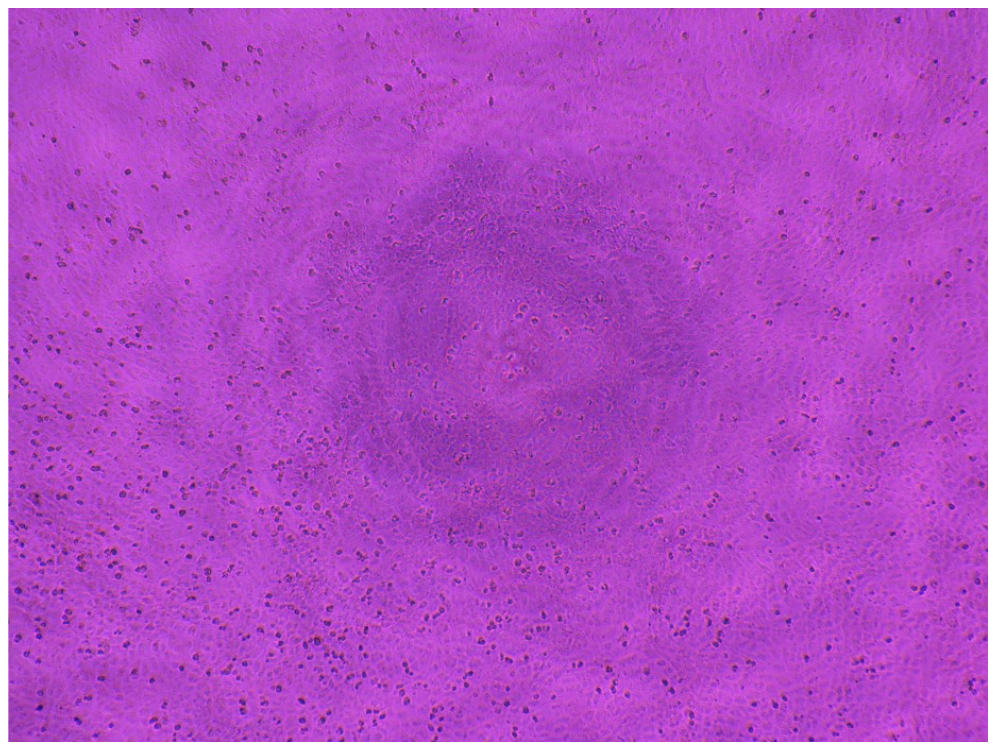 |
| Bimatoprost | 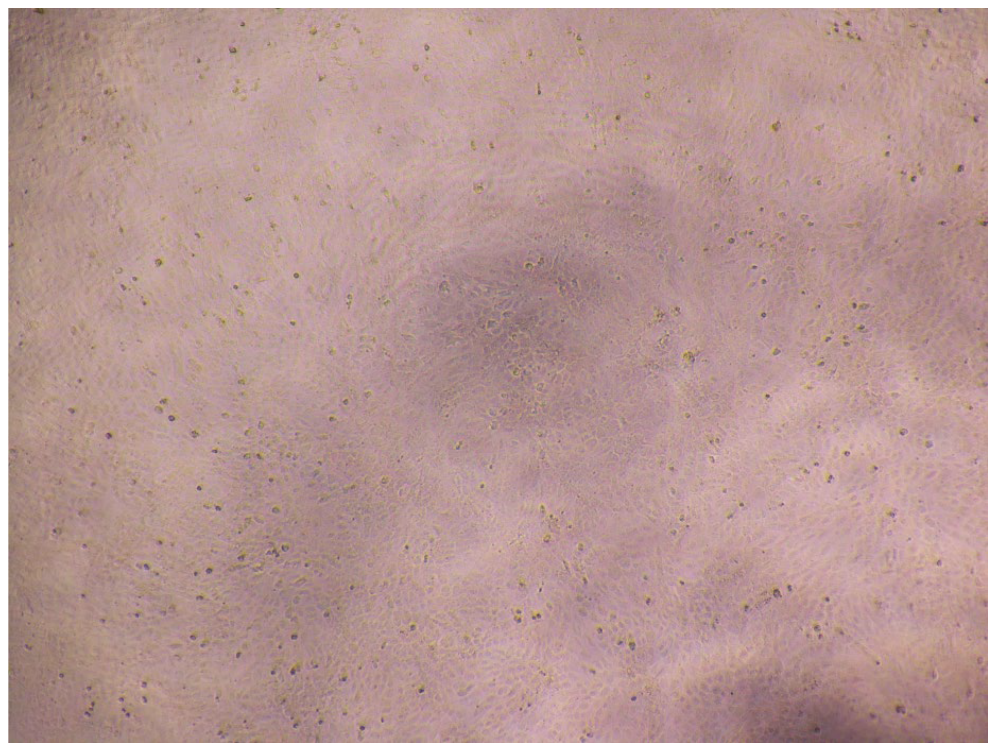 | 1,2-Dipalmitoyl-sn-glycerol | 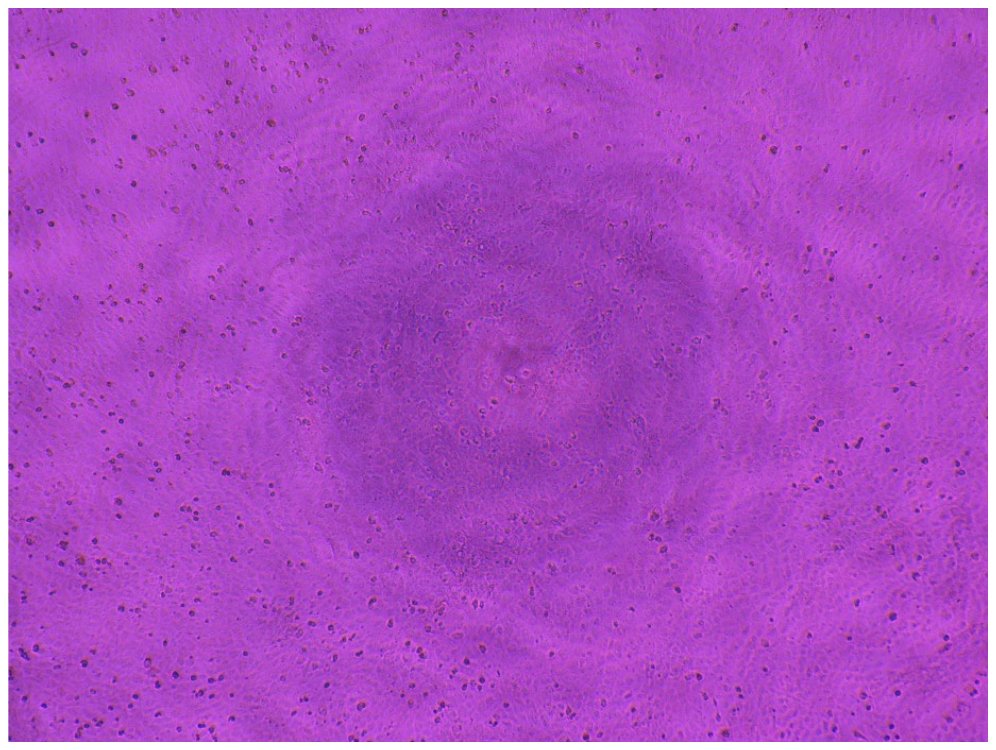 |
| Echinacoside | 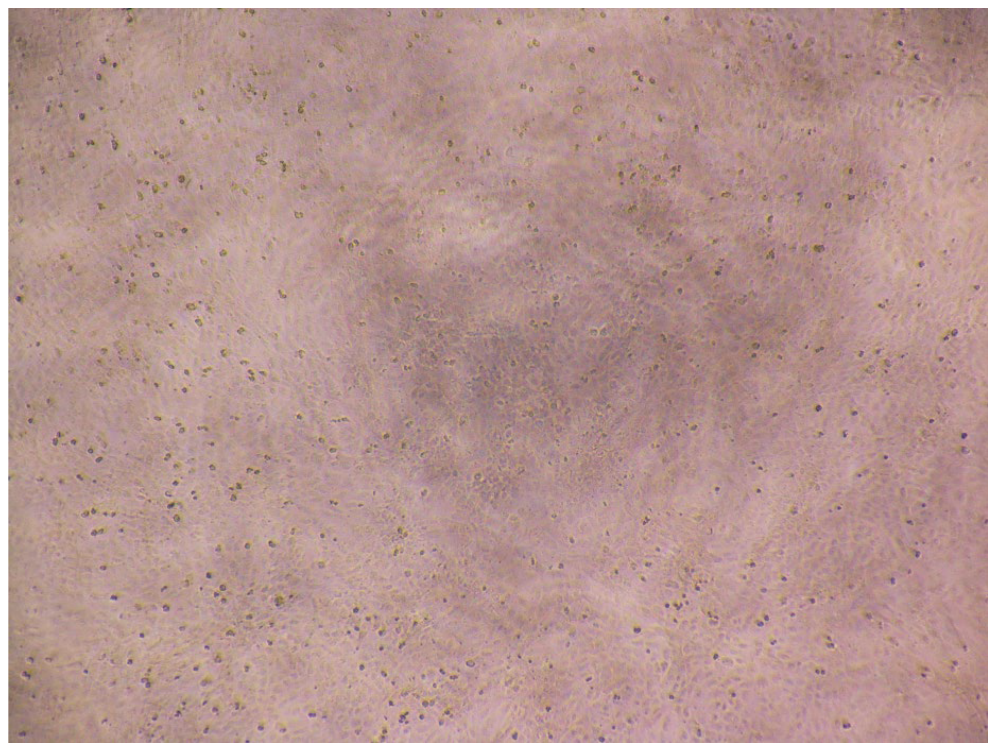 | rac-Glycerol 1-myristate | 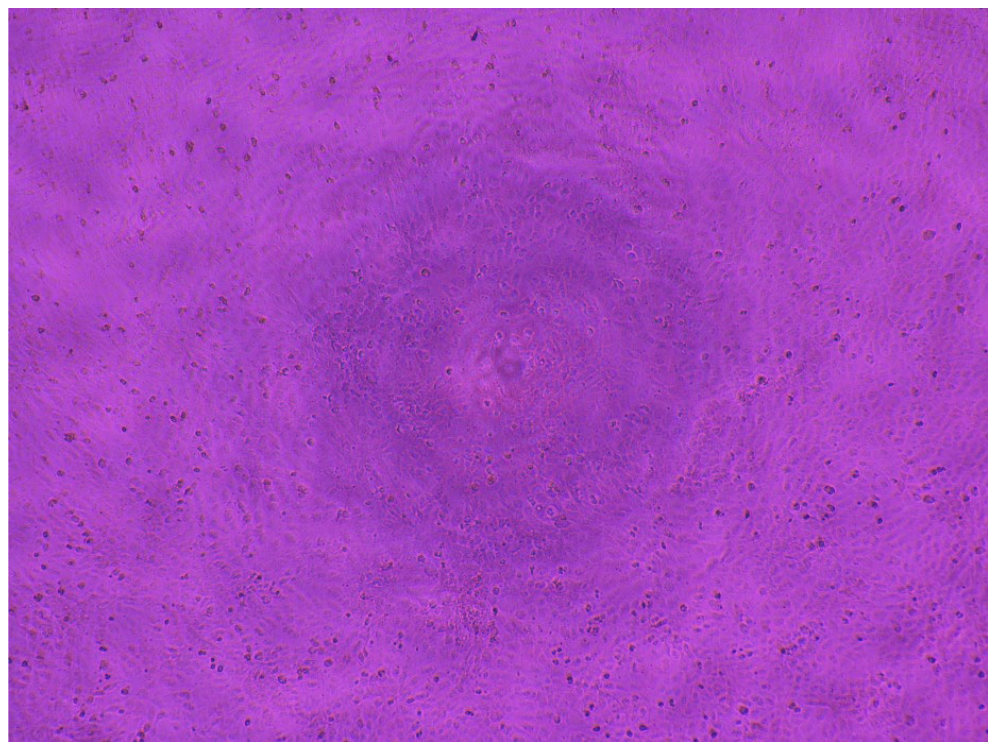 |
| CUDC101 | 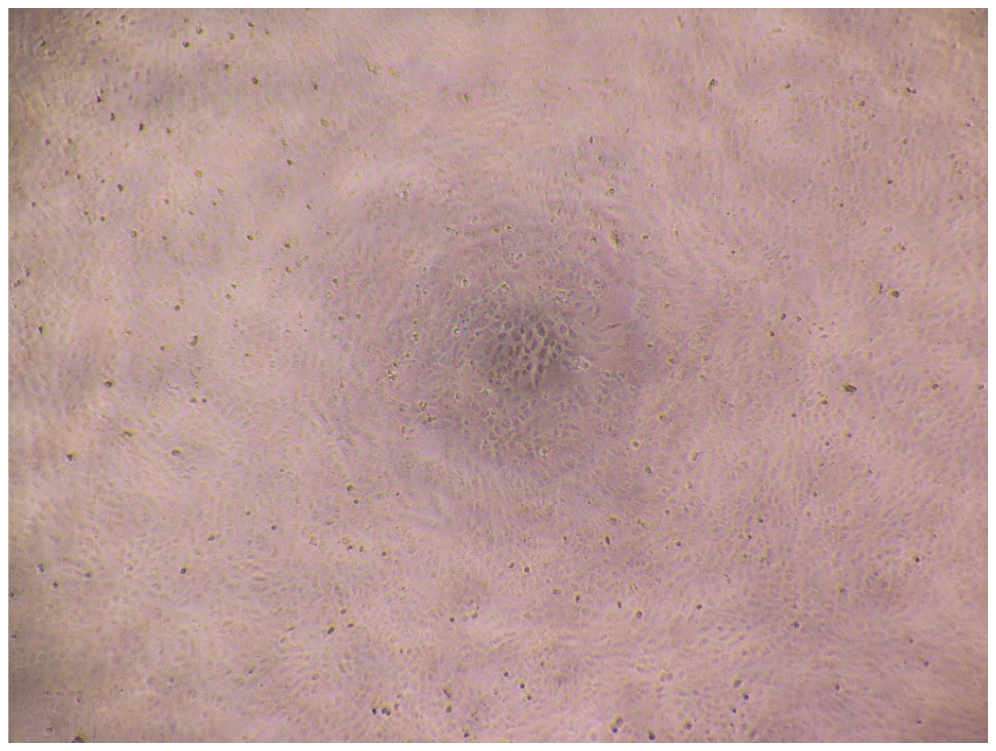 | Cefotiam Hexetil Hydrochloride | 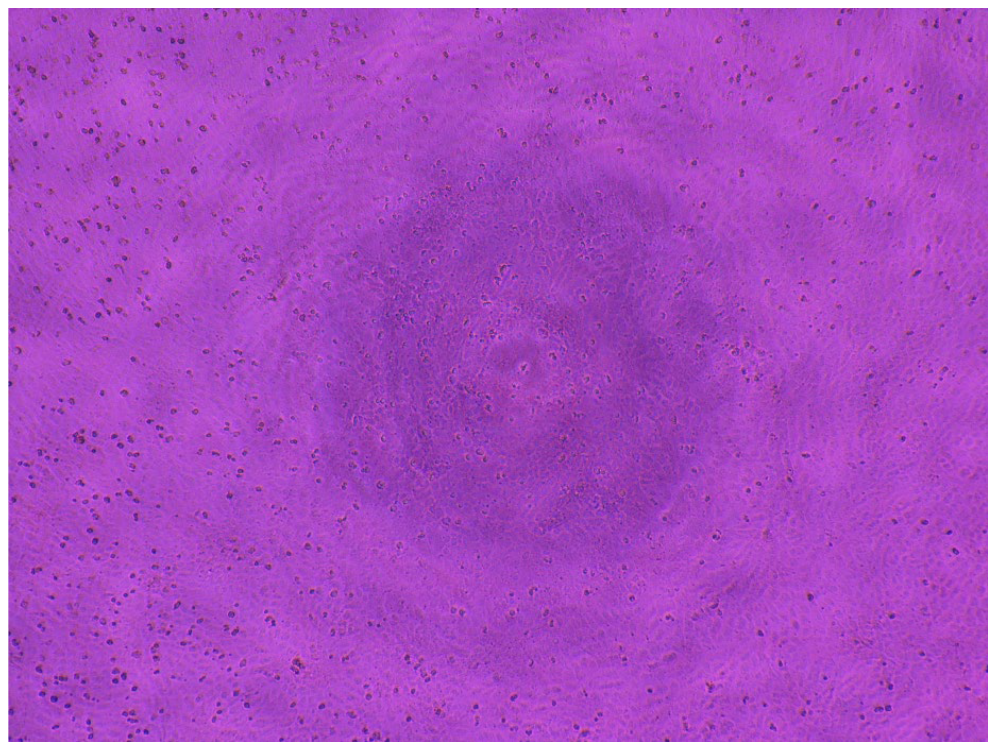 |
| PHT427 | 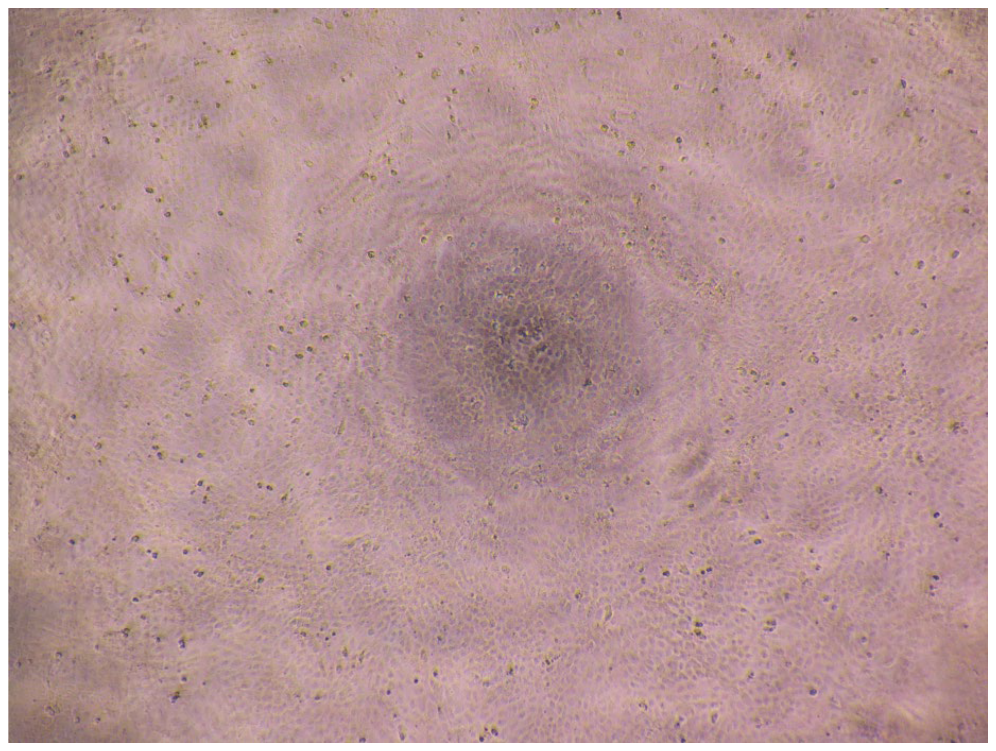 | Dabigatran Etexilate Mesylate | 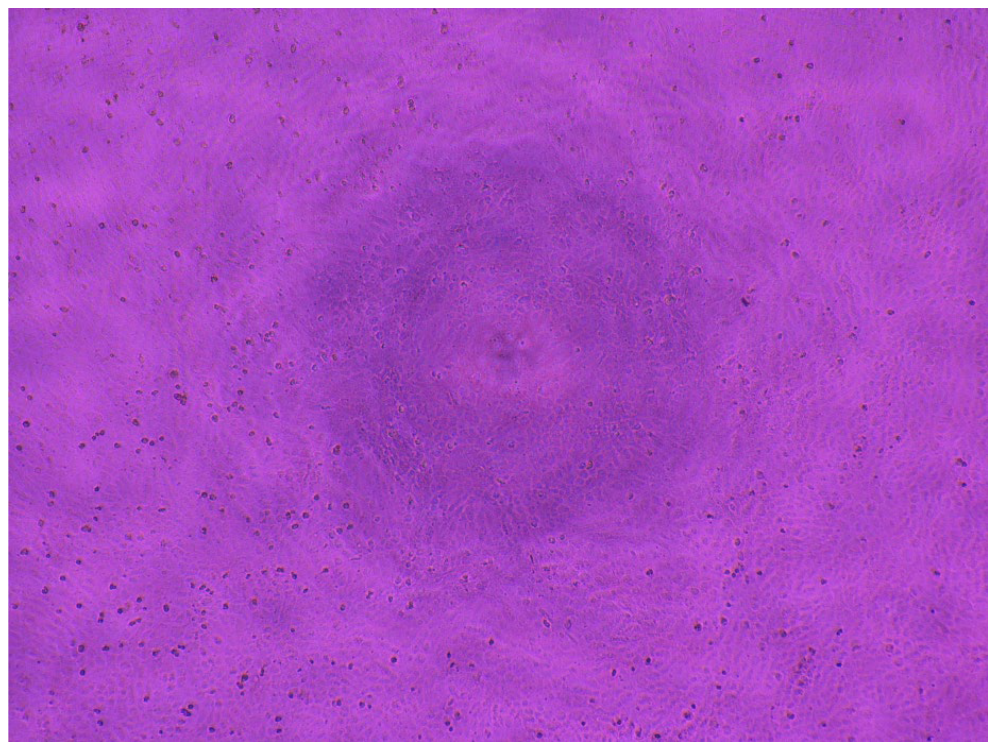 |
| Puromycin Dihydrochloride | 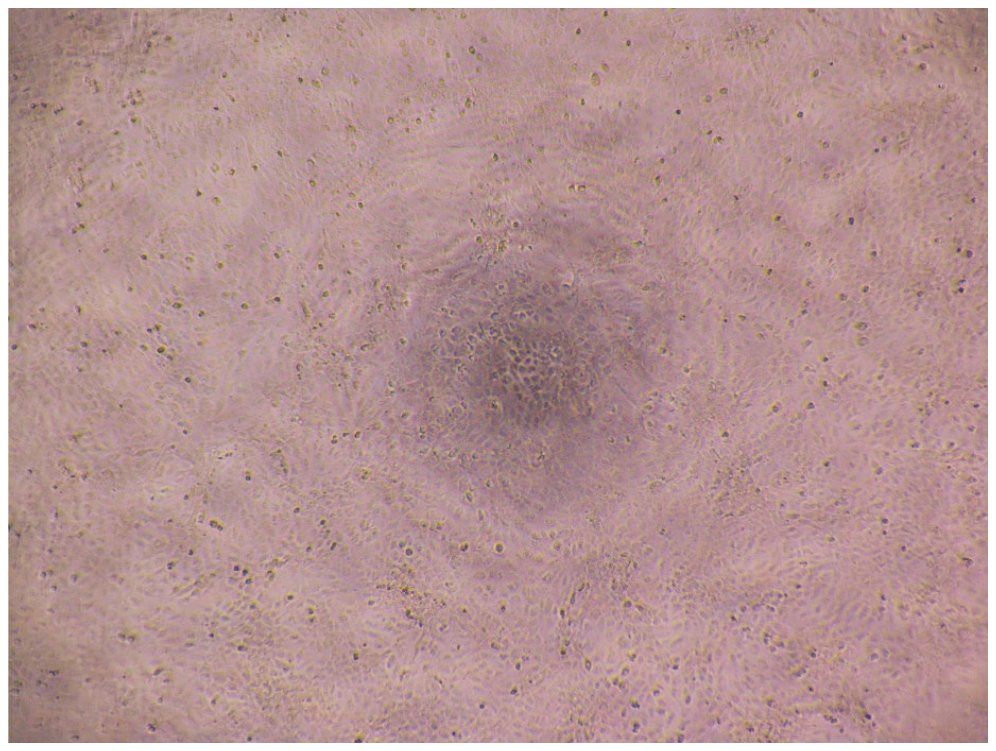 | Glycerol Tridecanoate | 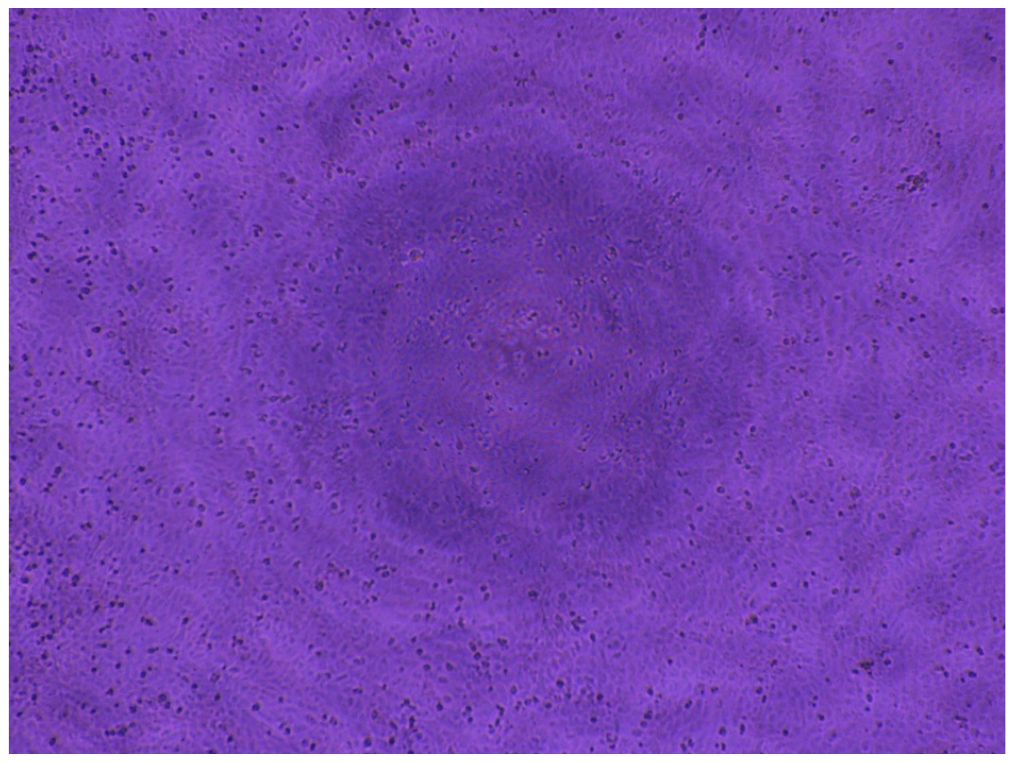 |
| UNC0638 | 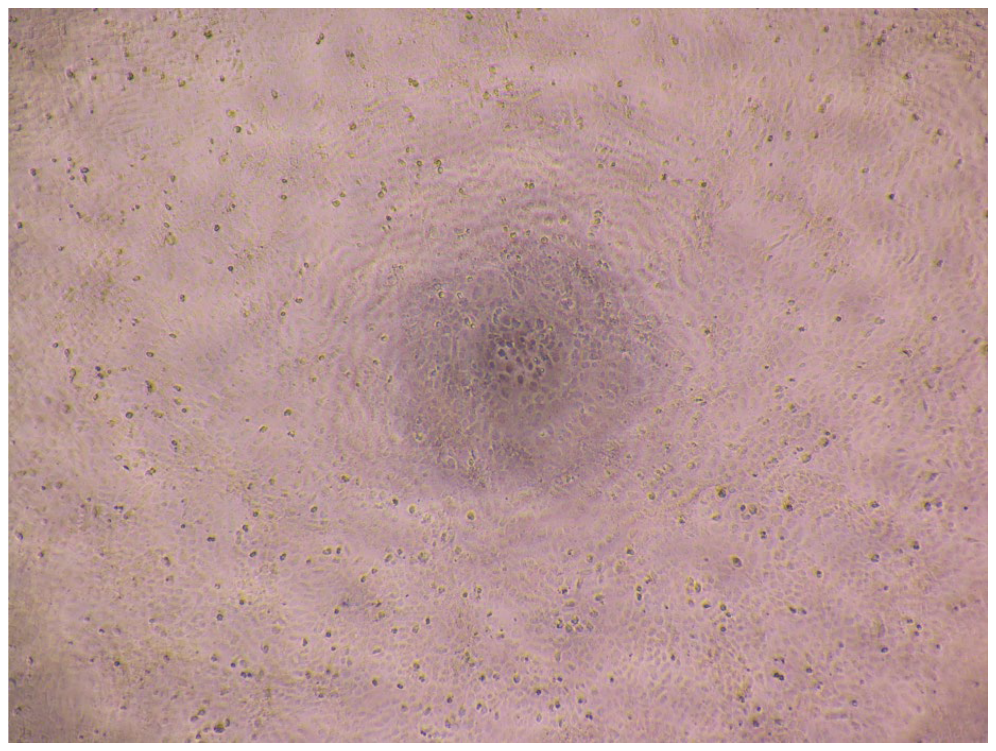 | ARV-825 | 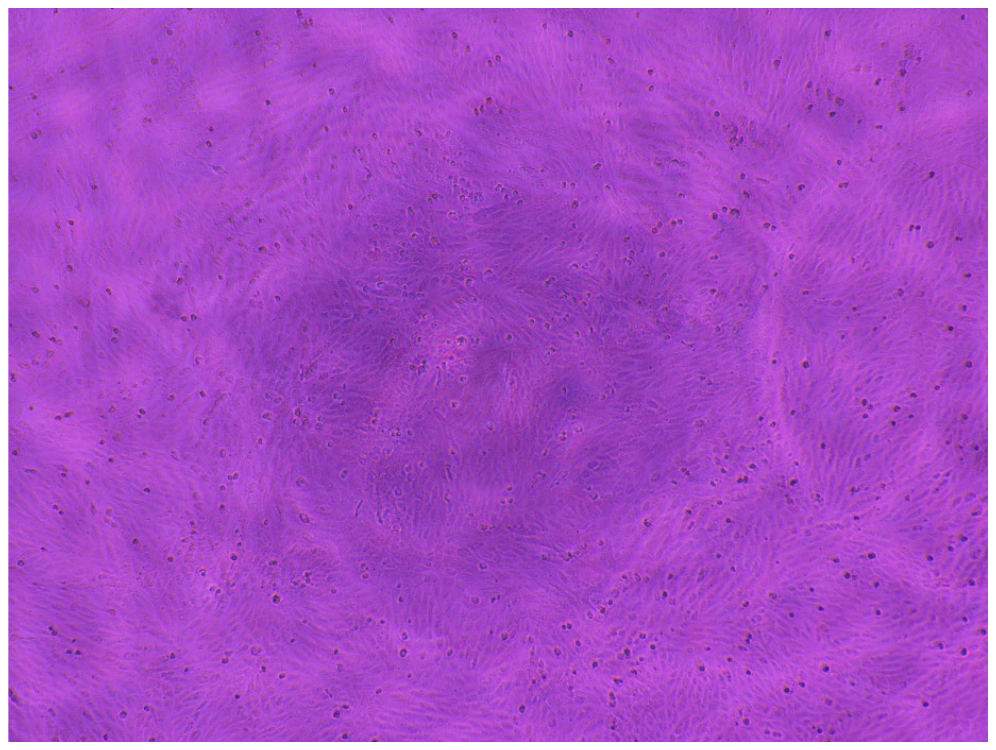 |
| Homoharringtonine | 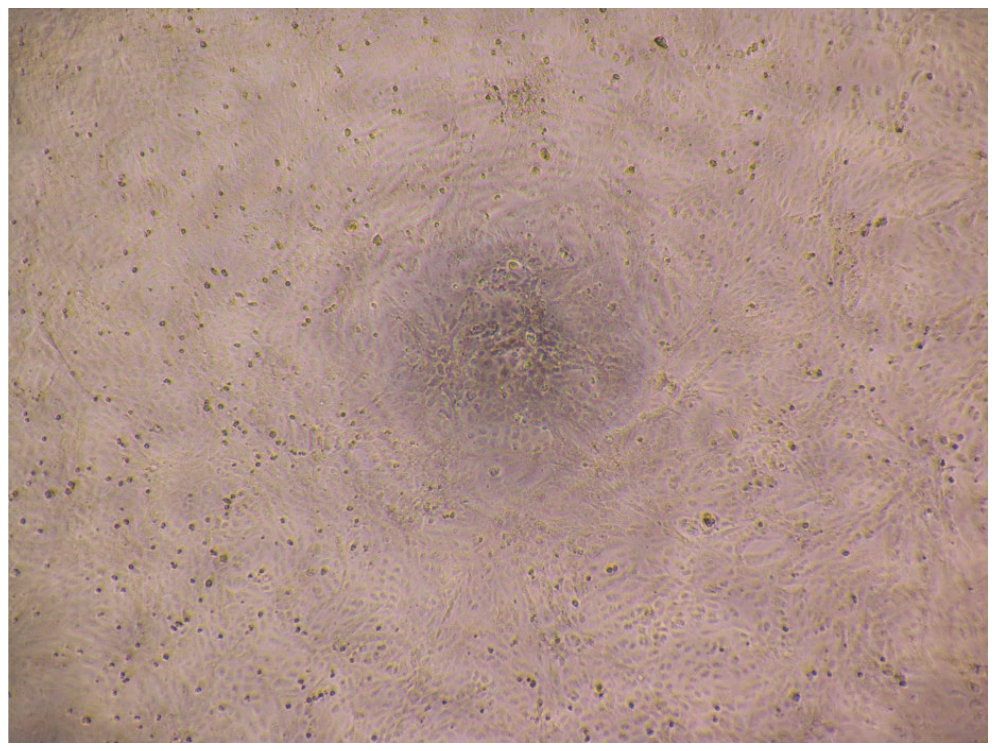 | Levobetaxolol Hydrochloride | 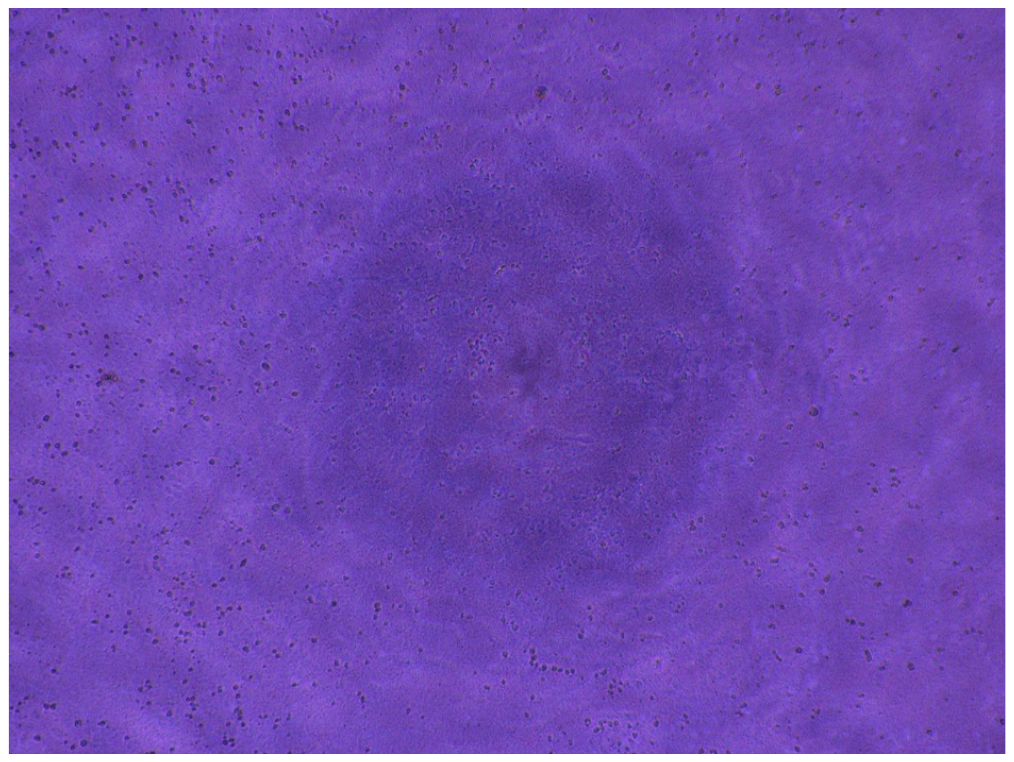 |
| Calpeptin | 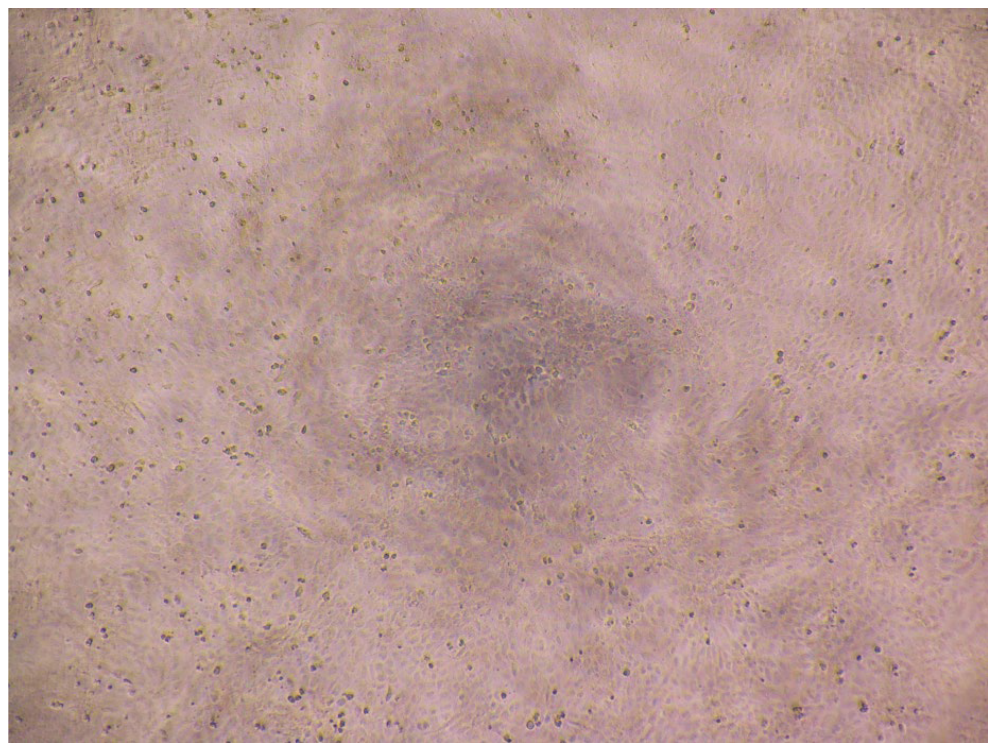 | Travoprost | 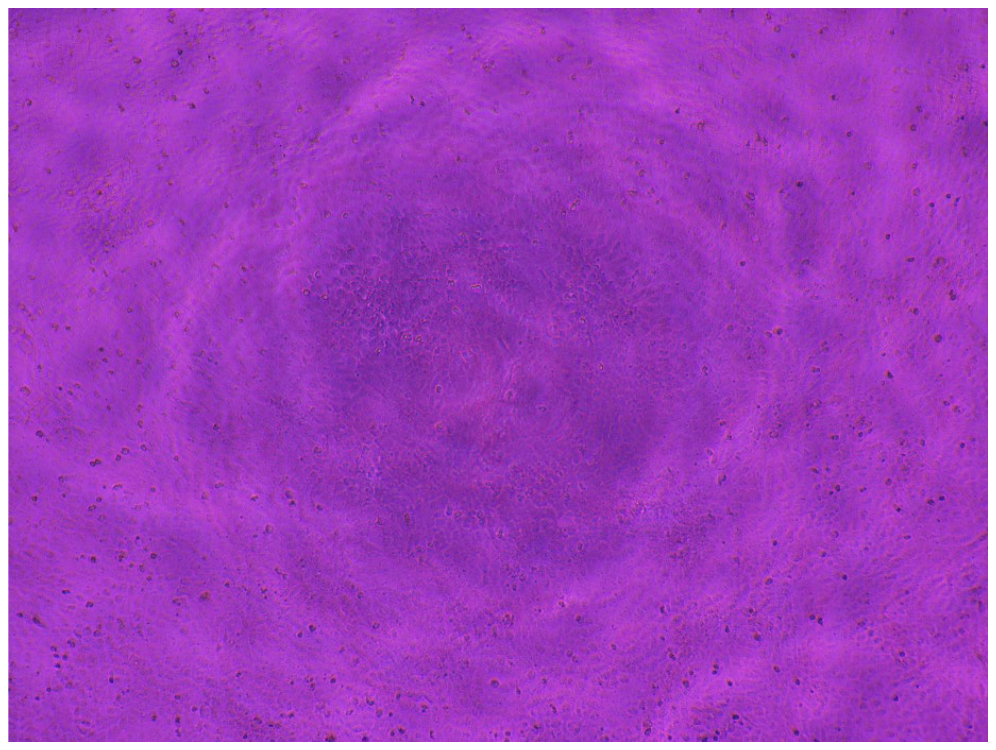 |
| Crocin II | 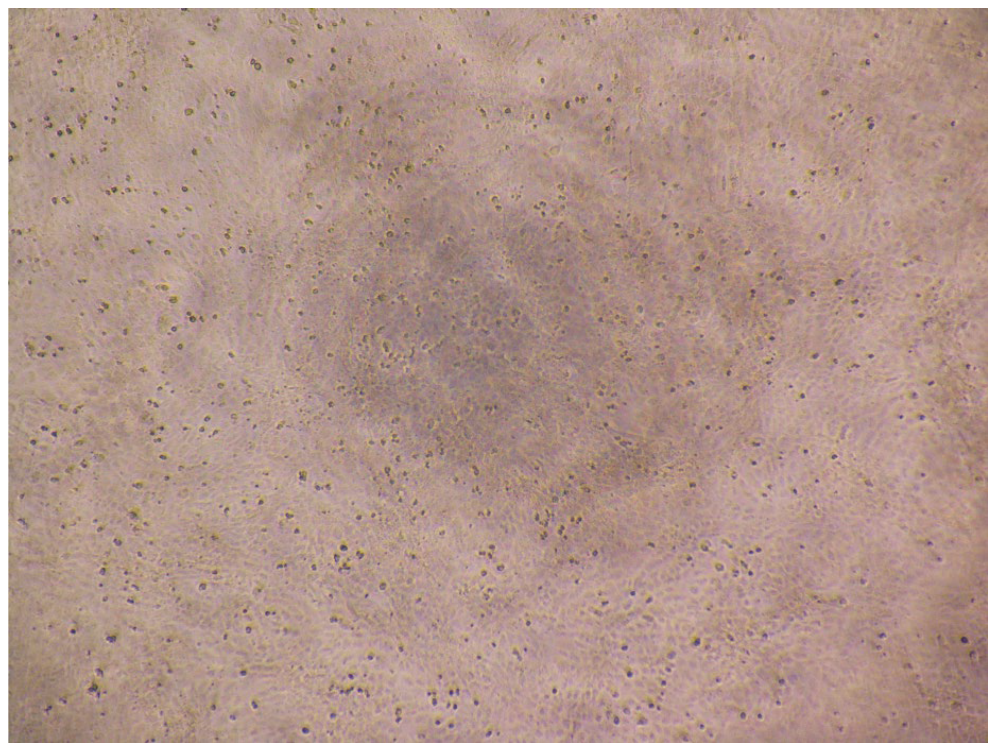 |  |  |

**Table S3**: List of primary hits identified from the primary screen performed using a flavonoid library.

| **Compound/Treatment** | **Image** | **Compound/Treatment** | **Image** |
| --- | --- | --- | --- |
| Uninfected | 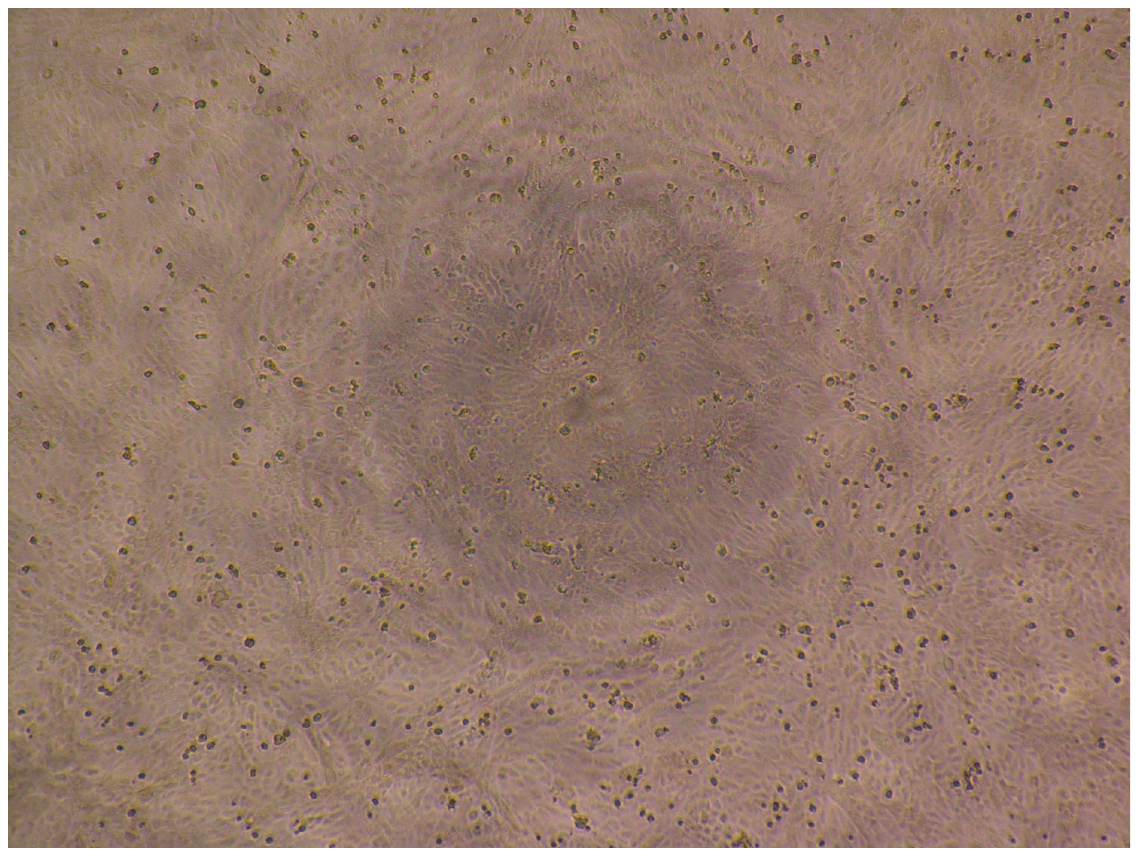 | 0.1% DMSO | 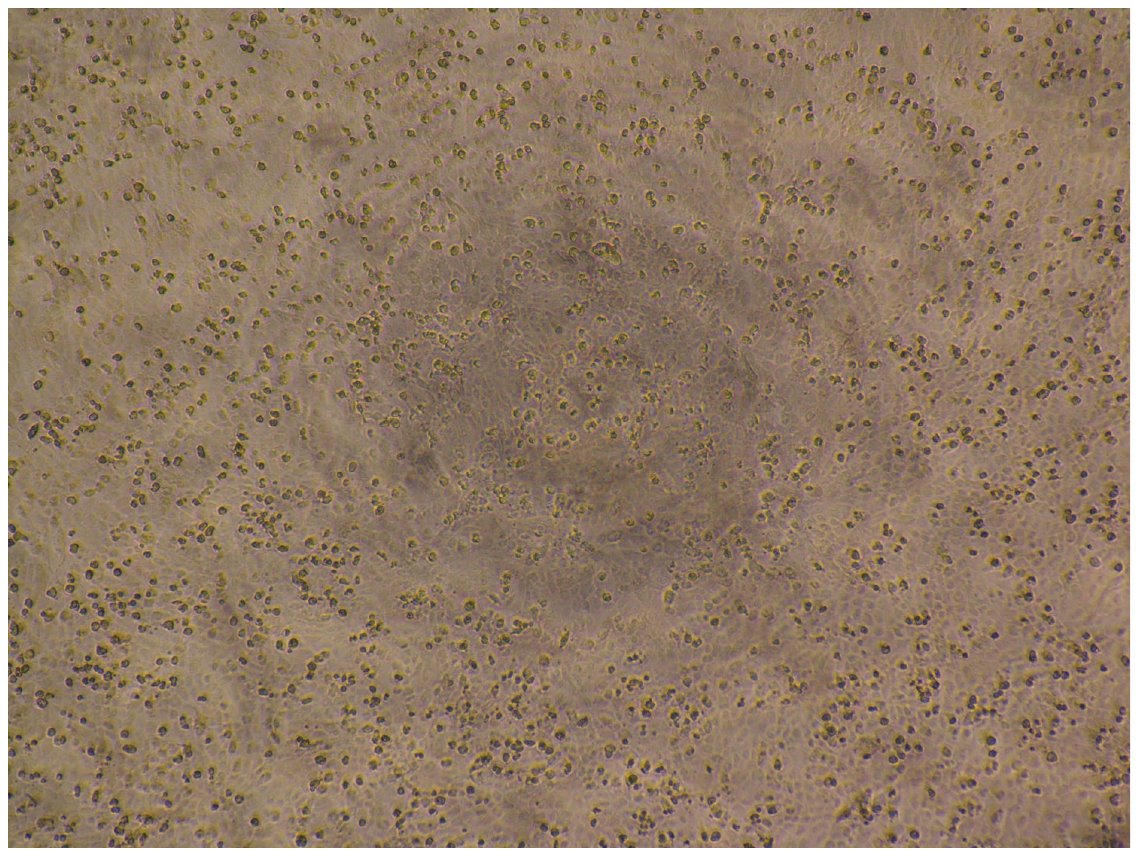 |
| Remdesivir | 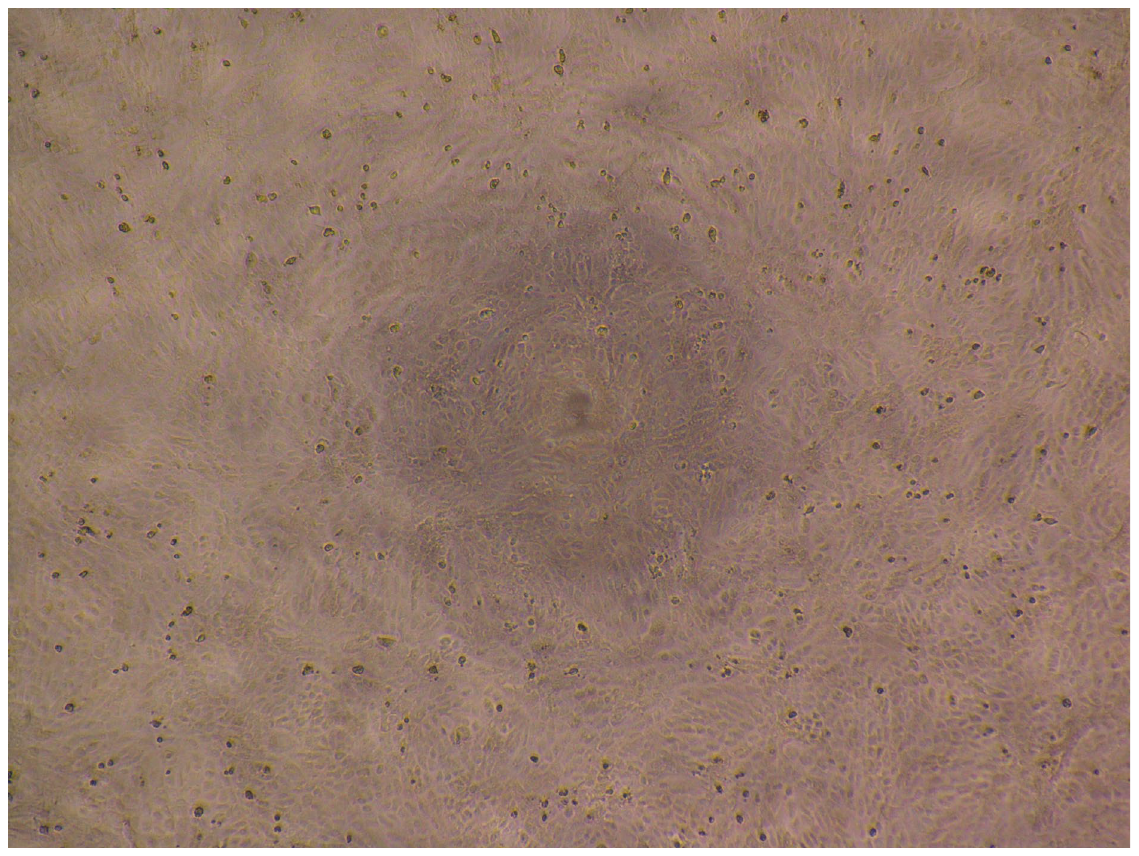 |  |  |
| ST002047 | 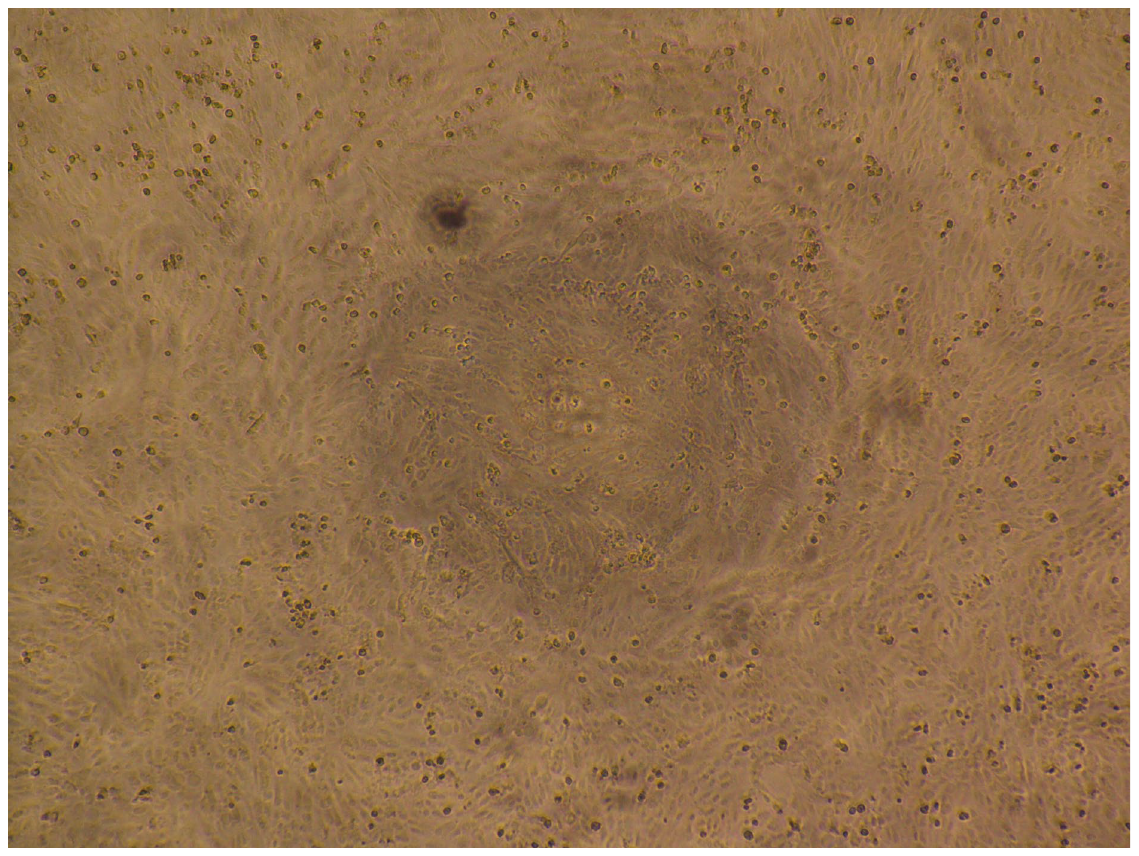 | ST079547 | 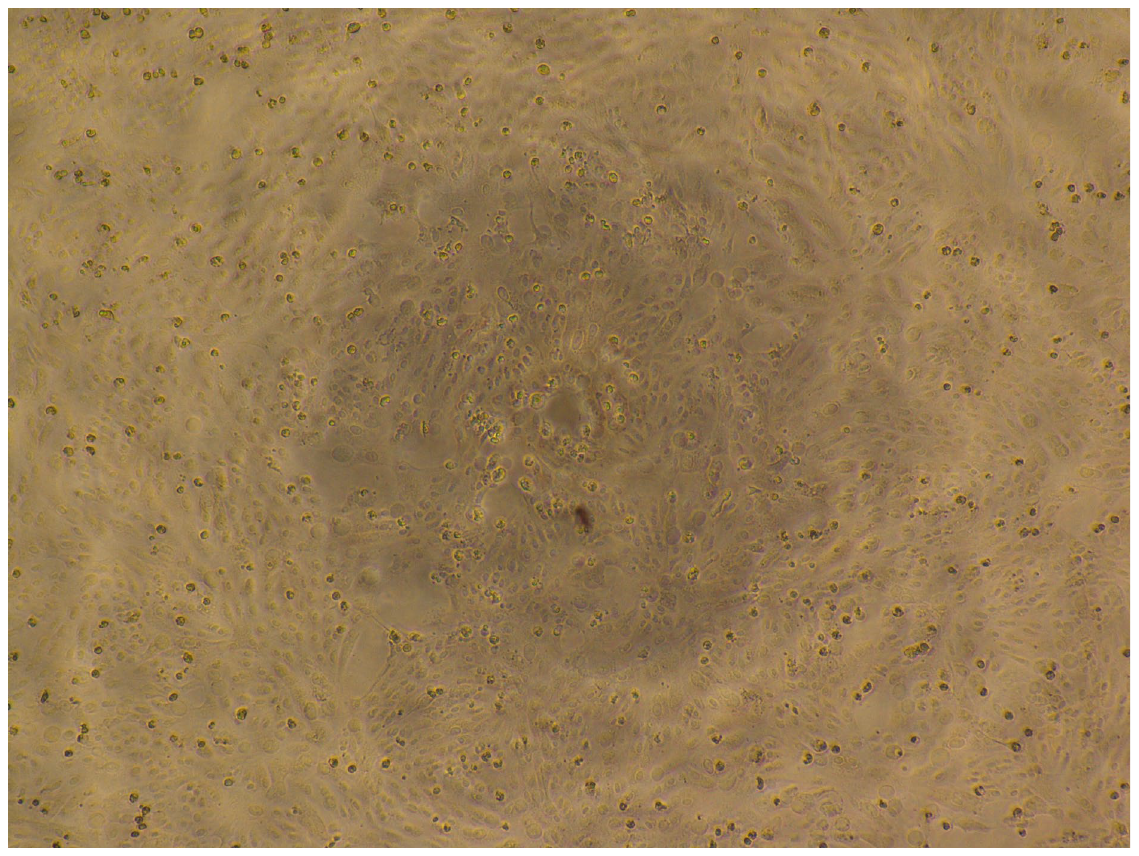 |
| ST014848 | 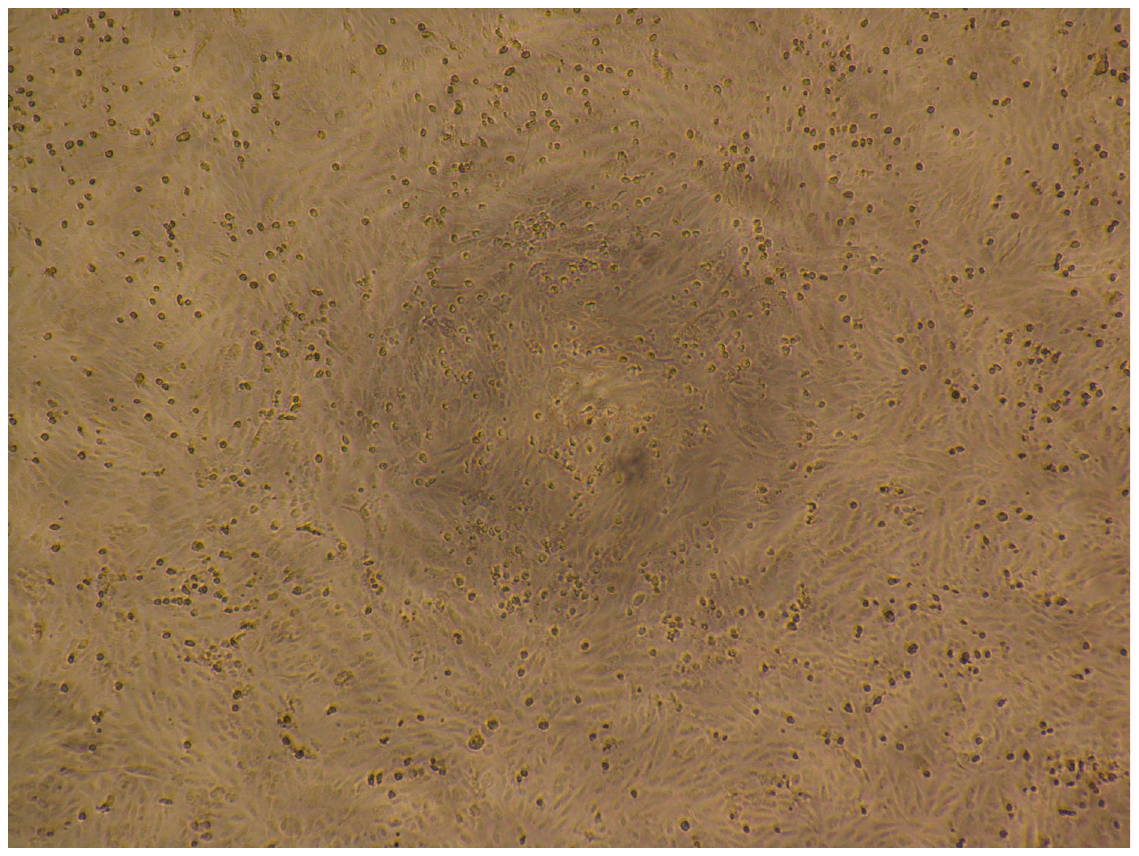 | ST085666 | 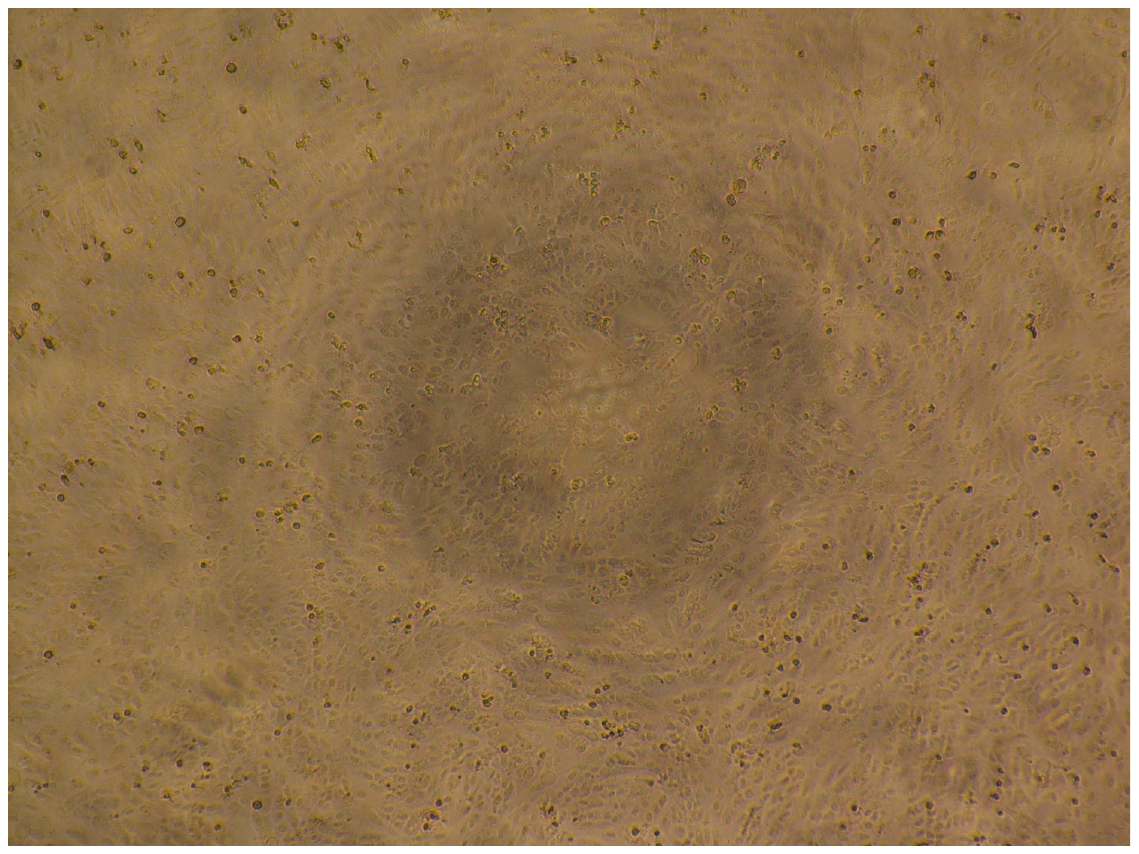 |
| ST055995 |  | ST076522 |  |
| ST056004 |  | ST083654 |  |
| ST056249 |  | ST081389 |  |
| ST056248 |  | ST076533 |  |
| ST057711 |  | ST095414 |  |
| ST058442 |  | ST097097 |  |
| ST057646 |  | ST094884 |  |
| ST059590 |  | ST095197 |  |
| ST069324 |  | ST095849 | 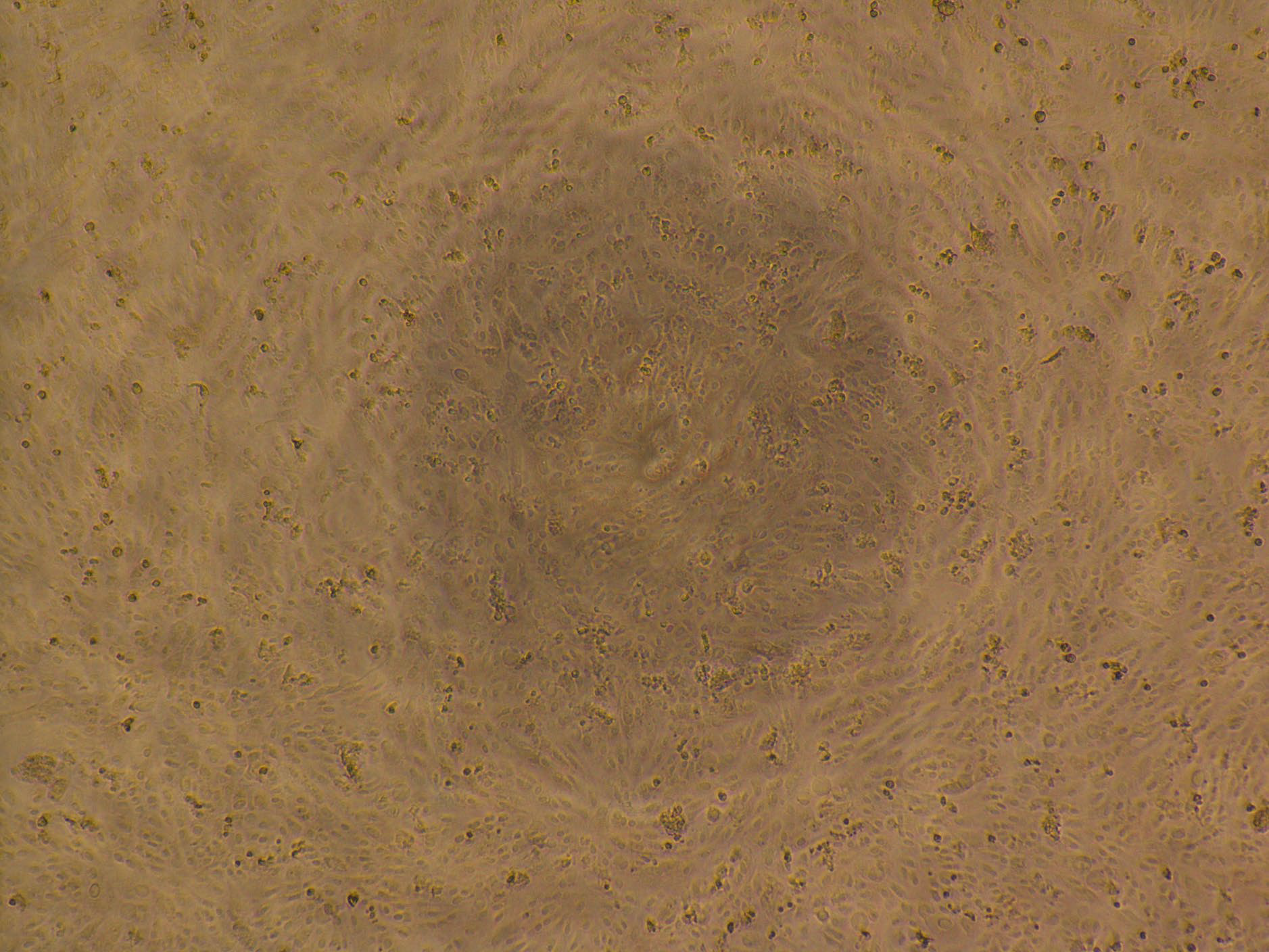 |
| ST075446 |  | ST097570 |  |
| ST076534 |  | ST020733 |  |
| ST079949 | 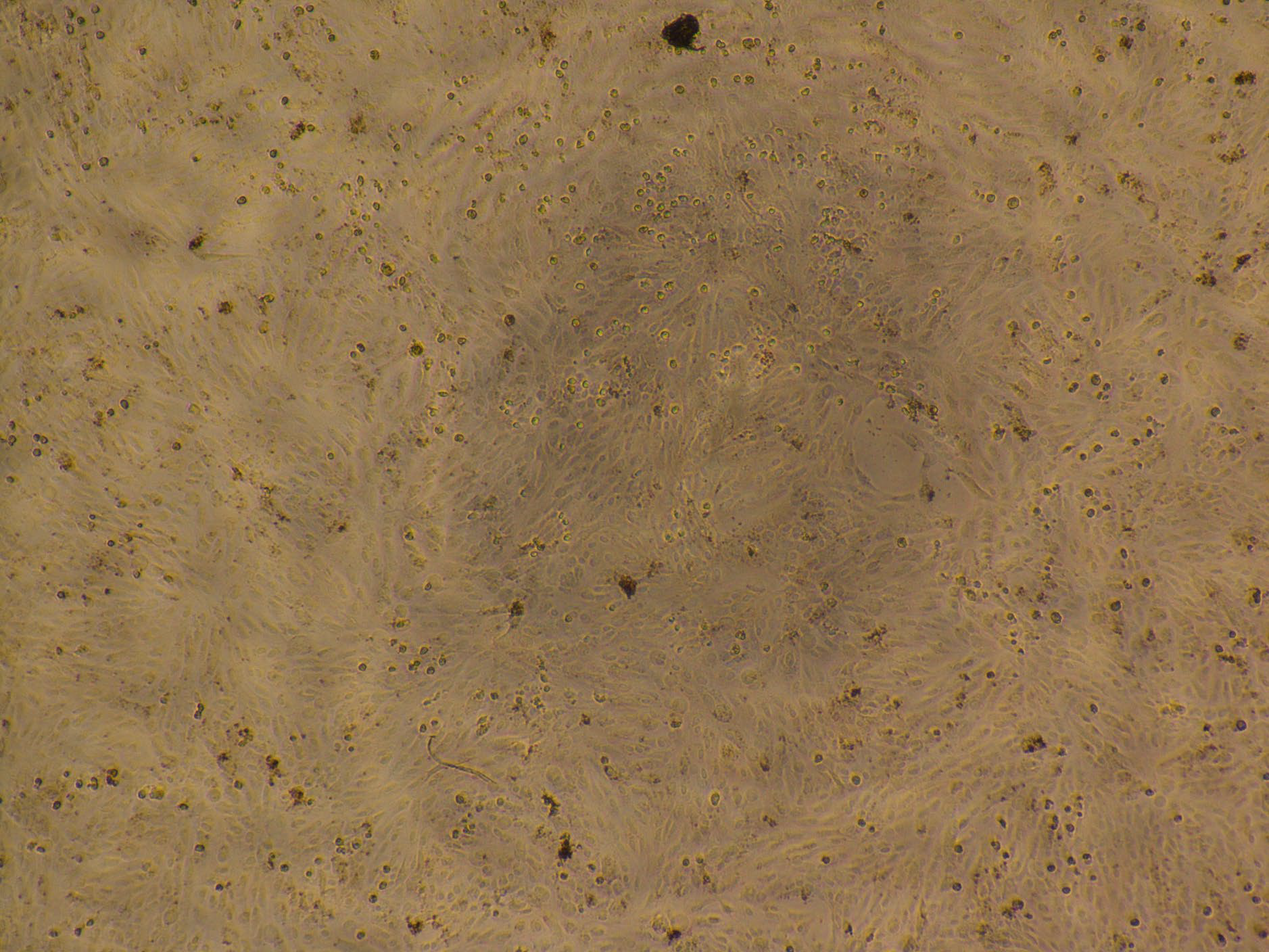 | ST074501 |  |
| ST083092 |  | ST074530 |  |
| ST083658 |  | ST077114 |  |
| ST077135 |  | ST077122 | 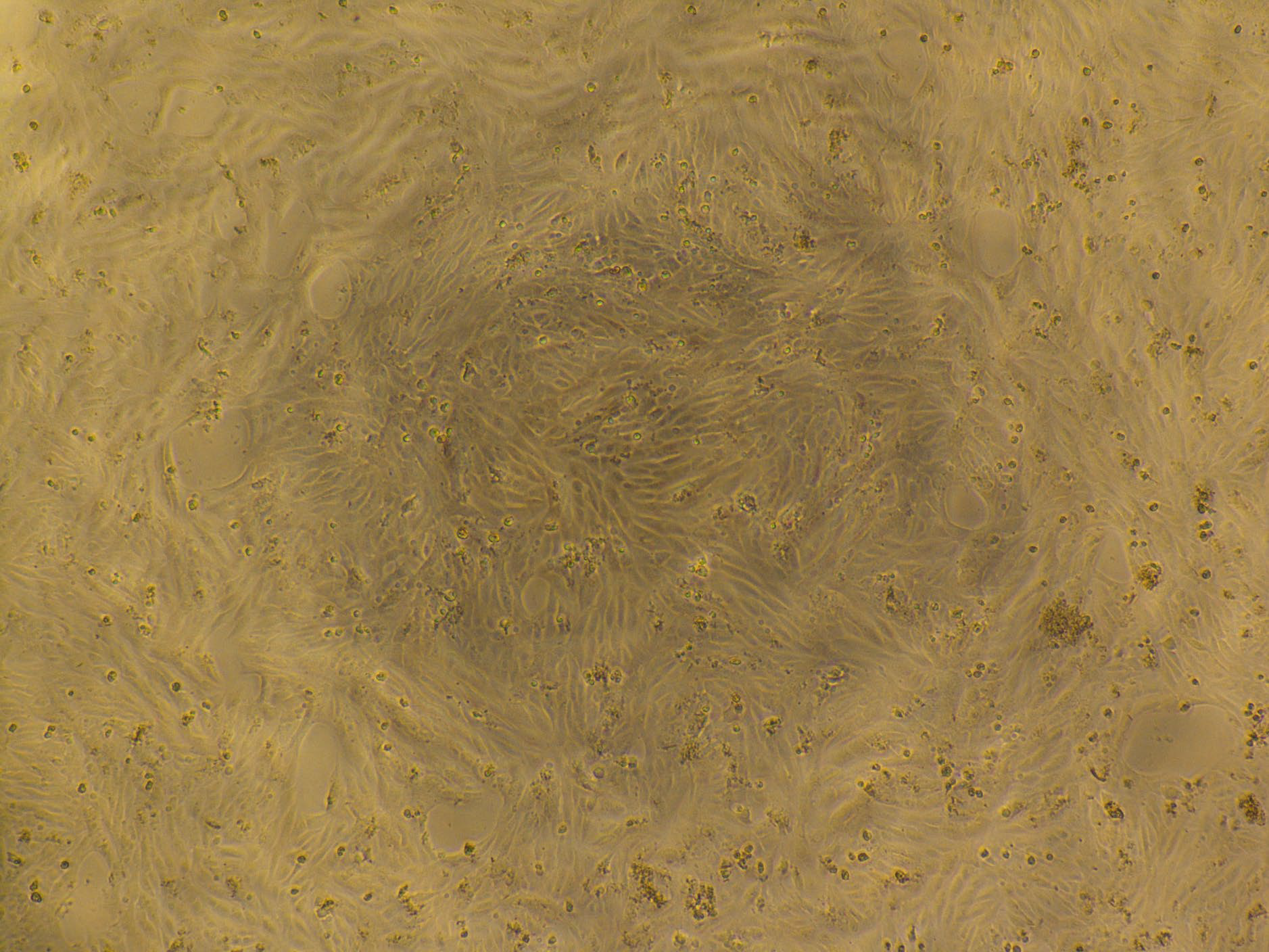 |

**Table S4**: Primary hits selected for further validation against SARS-CoV-2 infection.

| Pharmaceutical Class | No. of compounds with activity against SARS-CoV2 | | | Total no. of compounds for each class |
| --- | --- | --- | --- | --- |
|  | FDA-approved drugs | ACE2 Inhibitors | Flavonoids |  |
| Antiviral agents | 3 | 2 |  | 5 |
| Antibacterial agents | 9 | 4 |  | 13 |
| Antifungal agents | 3 |  |  | 3 |
| Antiparasitic agents | 2 |  |  | 2 |
| Anticancer agents | 3 | 3 |  | 6 |
| Antihistamines | 5 |  |  | 5 |
| Antihypertensives | 1 | 1 |  | 2 |
| Antiarrhythmic |  | 1 |  | 1 |
| Anticoagulants |  | 1 |  | 1 |
| Antispasmodic | 1 |  |  | 1 |
| Proton pump inhibitors | 2 |  |  | 2 |
| Ion channel blockers | 3 |  |  | 3 |
| Non-steroidal anti-inflammatory drugs | 3 |  |  | 3 |
| Anti-inflammatory agents | 2 | 4 |  | 6 |
| Phosphodiesterase inhibitors | 1 |  |  | 1 |
| Lipid, sterol metabolism inhibitors | 1 | 1 |  | 2 |
| Signalling kinase inhibitors | 2 | 2 |  | 4 |
| Neurotransmitter inhibitors | 3 |  |  | 3 |
| Nucleic acid synthesis inhibitors | 2 |  |  | 2 |
| Histone deacetylase inhibitors | 0 | 1 |  | 1 |
| Flavonoids |  | 1 | 34 | 35 |
| Others | 10 | 10 |  | 20 |
| Total | 56 | 31 | 34 | 121 |

**Table S5**: CC_50_, IC_50_, and SI values observed from validation of primary hits in Vero E6 cells.

| **Compound** | **CC50 (μM)** | **IC50 (μM)** | **Selectivity Index** |
| --- | --- | --- | --- |
| **Citicoline** | > 20 | - | - |
| **Pravastatin Sodium** | > 20 | < 0.01 | >2000 |
| **Tenofovir Alafenamide** | > 20 | - | - |
| **Imatinib Mesylate** | > 20 | < 0.01 | >2000 |
| **Calcitriol** | > 20 | < 0.01 | >2000 |
| **Dexlansoprazole** | > 20 | < 0.01 | >2000 |
| **Prochlorperazine Dimaleate** | > 20 | - | - |

Note: CC_50_ and IC_50_ values were derived from data obtained from cell viability and dose-dependent inhibition studies respectively.

**Table S6**: CC_50_, IC_50_, and SI values observed from validation of primary hits in Huh7 cells.

| Compound | CC50 (μM) | IC50 (μM) | Selectivity Index |
| --- | --- | --- | --- |
| Citicoline | > 20 | - | - |
| Pravastatin Sodium | > 20 | - | - |
| Tenofovir Alafenamide | > 20 | - | - |
| Imatinib Mesylate | > 20 | - | - |
| Calcitriol | 4.692 | - | - |
| Dexlansoprazole | > 20 | 2.479 | > 8.0678 |
| Prochlorperazine Dimaleate | 15.73 | - | - |

Note: CC_50_ and IC_50_ values were derived from data obtained from cell viability and dose-dependent inhibition studies respectively
